# Epithelial–mesenchymal cell state heterogeneity predetermines differential phospho-signaling responses to EGF stimulation

**DOI:** 10.1101/2025.07.05.658545

**Authors:** Felix V. Kohane, Chantelle Johnstone, Daniel P. Neumann, Ihuan Gunawan, Tim Huang, Fatemeh Vafaee, Christine L. Chaffer, John G. Lock

**Affiliations:** School of Biomedical Sciences, Faculty of Medicine and Health, University of New South Wales, Sydney, NSW, Australia; School of Computer Science and Engineering, Faculty of Engineering, University of New South Wales, Sydney, NSW, Australia; School of Biotechnology and Biomolecular Sciences, Faculty of Science, University of New South Wales, Sydney, NSW, Australia; UNSW Data Science Hub, University of New South Wales, Sydney, NSW, Australia; UNSW Artificial Intelligence Institute, University of New South Wales, Sydney, NSW, Australia; Garvan Institute of Medical Research, Darlinghurst, Sydney, NSW, Australia; St. Vincent’s Clinical School, UNSW Medicine, University of New South Wales, Sydney, NSW, Australia; The Kinghorn Cancer Centre, Darlinghurst, Sydney, NSW, Australia; Ingham Institute for Applied Medical Research, Liverpool, NSW, Australia

## Abstract

Understanding why isogenic cancer cells respond differently to equivalent oncogenic stimuli is vital for optimizing anticancer therapies. Emerging evidence suggests that pre-existing differences in cell state may modulate signaling responses to new stimuli, but the interplay of specific cell states and signals remains unclear. We investigated whether epithelial–mesenchymal (E/M) state, a major axis of cancer cell heterogeneity, influences signaling responses to epidermal growth factor (EGF), a critical oncogenic stimulus in non-small cell lung cancer (NSCLC). We imaged >64,000 A549 NSCLC cells labeled for DNA, F-actin and alternate signaling markers (p-AKT-S473, p-AKT-T308, p-ERK or p-S6) after acute stimulation. Quantitative single-cell morphological and spatial profiling defined a stimulus-invariant ‘E/M state landscape’ over which EGF signaling responses were compared. This revealed state-dependent differences in signal-activation magnitudes, dynamics and subcellular routing. AKT responses exhibited phosphosite- and compartment-specific dynamics across states, with epithelial cells showing strong, transient membrane-localized S473 and higher internalized T308, whereas mesenchymal cells displayed weaker but sustained nuclear and ruffle-localized S473. Regression-based computational multiplexing concurrently inferred all signaling responses per cell, mapping state-dependent divergence in multi-molecular signaling trajectories. E/M state thus pre-determines distinctive spatiotemporal profiles of EGF-induced signaling, with implications for signaling functions and anti-signaling therapy responses across E/M state-diverse tumors.

## 1 Introduction

Like many cancers, non-small cell lung cancer (NSCLC) is characterized by pronounced intratumoral heterogeneity. This poses significant challenges to effective treatment [1, 2, 3, 4]. While oncogenic driver mutations, such as those in the epidermal growth factor receptor (EGFR), are well-established therapeutic targets, signaling responses to incident ligand stimuli frequently exhibit substantial variability at the single-cell level, even within genetically identical tumor cell populations [5, 6, 7]. Such variability is commonly attributed to differences in ligand availability or stochastic molecular fluctuations and is considered ‘noise’, yet mounting evidence suggests that pre-existing differences in cell state also shape how normal and transformed cells interpret cues from their microenvironment [8, 9, 10, 11, 12]. Understanding how diversity in the acute cell signaling response arises from pre-existing heterogeneity in cell states is crucial for understanding oncogenic signaling variability in cancer and its impact on the effectiveness of targeted therapies.

The functional outcomes of signaling pathways depend not only on the specific molecular components activated but also on a number of key signaling ‘dimensions’, including signaling magnitude, dynamics, and localization. For example, the magnitude of AKT activity contributes to metabolic adaptation, with insulin inducing stronger responses than either EGF or PDGF, leading to enhanced GLUT4-mediated glucose uptake and the suppression of AMPK activity [13, 14]. In addition to signal strength, dynamic fluctuations and oscillations in AKT activity temporally coordinate with glycolytic flux and AMPK activation, highlighting a link between AKT dynamics and the regulation of metabolic stability in rapidly proliferating cancer cells [13]. Similar dynamic cell fate encoding is present in the MAPK pathway. In breast cancer cells, transient ERK signaling promotes proliferation, whereas sustained ERK activation drives differentiation [15]. Recent single-cell analyses further show that combined ERK and AKT dynamics can predict stochastic cell-division events [16, 17]. Both the strength and kinetics of signaling responses thus encode information influencing clinically relevant cellular fates [6, 18, 19].

Signaling outcomes are also highly dependent on the subcellular spatial location in which activation occurs [20]. Spatial compartmentalization, such as across various organelles, plays a critical role in tuning signaling responses. For instance, endosomes act as platforms for routing AKT and ERK signaling [21, 22, 23, 20], mitochondrial membrane-associated kinases modulate local mitogen-activated protein kinase (MAPK) activity [24], and phosphoinositide 3-kinase (PI3K) signaling is enriched at focal adhesions [25]. Such compartment-specific activities are regulated by the local availability of distinct lipid species, scaffold proteins, kinases, and phosphatases [26, 27]. Spatially restricted signaling can also influence cell fate decisions. For example, nuclear ERK signaling promotes neuronal differentiation, whereas cytosolic ERK triggers myogenic differentiation in muscle precursors [19]. ERK activity at the plasma membrane has a lower activation threshold than in the cytosol, enabling weak stimuli to elicit strong responses, thus illustrating how distinct subcellular domains create unique signaling environments [28]. These studies highlight signaling and its outcomes as highly context-sensitive and tunable across multiple signaling dimensions. Yet, how such multidimensional tuning may be predetermined and modulated by differences in specific cell states remains largely unknown. A key axis of cellular state heterogeneity in epithelial cancers like NSCLC is the epithelial–mesenchymal (E/M) spectrum, within which cells can undergo plastic transitions via the epithelial–mesenchymal transition (EMT) and its reverse, the mesenchymal–epithelial transition (MET) [29]. Cells spanning this continuum exhibit diverse morphologies, gene expression, and functional behaviors, including differences in proliferation, motility, and therapy resistance [30, 31]. E/M plasticity is closely associated with metastatic potential, treatment evasion, and tumor recurrence [32] and is well-established as a source of molecular diversity within tumors [33, 34, 35]. This plasticity is regulated by developmental pathways often aberrantly activated in cancer, including transforming growth factor (TGF)-β, EGF, Wnt, and Notch [36, 37, 38, 39, 40]. While much attention has been paid to how these pathways initiate the EMT [41, 42, 43], relatively little is known about how a cell’s position within the E/M state spectrum influences its capacity to decode and respond to newly incident signaling stimuli.

Here, we investigated whether E/M state modulates early signaling responses to EGF in NSCLC cells, reflecting the clinical significance of EGF signaling (and its therapeutic targeting) in NSCLC. Using immunofluorescence imaging, quantitative single-cell morpho-spatial profiling, and a computational multiplexing framework [44], we mapped phospho-signaling dynamics across thousands of individual A549 cells. Focusing on four key signaling nodes — phosphorylated (p)-AKT-T308, p-AKT-S473, p-ERK, and p-S6 — we characterized phospho-signal magnitudes, dynamics, and subcellular localization across a continuum of E/M phenotypes. By integrating representation learning [45] and pseudotime trajectory analysis, we show that E/M state modulates not only the amplitude and dynamics of signaling, but also its spatial routing and multi-molecular coordination. Our findings reveal that naturally occurring E/M heterogeneity is a major determinant of how cancer cells process incoming oncogenic signals. This underscores the need to incorporate aspects of cell phenotypic diversity (such as E/M state) into models of oncogenic signaling and therapeutic response.

## 2 Methods

### 2.1 Cell culture and EGF stimulation

A549 lung adenocarcinoma cells (American Type Culture Collection, ATCC; # CCL-185) were cultured in Dulbecco’s Modified Eagle Medium (DMEM; Thermo Fisher Scientific) supplemented with 10% fetal bovine serum (FBS; Sigma-Aldrich Pty Ltd). Cells were maintained at 37*^◦^*C, 5% CO_2_, and 21% O_2_. Passaging involved rinsing cells with 1X phosphate-buffered saline (PBS), detachment using Trypsin-EDTA (Thermo Fisher Scientific) for 3 min, centrifugation at 1500 rpm for 5 min, and subsequent reseeding. Cells were passaged twice per week and used at passage numbers 3–20. Cells were routinely tested for mycoplasma.

For EGF activation, 2750 cells per well were seeded in 300 µL media onto 96-well glass bottom plates (Cellvis; #P96-1.5H-N) and allowed to attach for 24 hr. Full growth media was subsequently replaced with serum-free DMEM by gradual dilution and cells were further incubated for 24 hr. Human recombinant EGF (Cell Signaling Technology, CST; #8916) was then added to wells at a final concentration of 20 ng/mL at specified time intervals based on the treatment duration (2, 5, 20, or 60 min). EGF was added such that all wells were fixed at the same end time.

### 2.2 Immunofluorescence labeling

Immunofluorescence was performed at room temperature with an OT-2 liquid-handling robot (Opentrons) using custom pipelines. Cells were fixed with 4% paraformaldehyde (Electron Microscopy Sciences) diluted in tris-buffered saline (TBS) for 20 min, permeabilized with 0.1% Triton X-100 (Sigma-Aldrich) in TBS for 10 min and blocked with Intercept (TBS) blocking buffer (LI-COR) for 15 min. Primary antibodies diluted in Intercept blocking buffer were added to wells for 2 hr and agitated on a rocker. Cells were incubated with anti-mouse or rabbit F(ab*^′^*)_2_ secondary antibodies (Gt; 1:1000; CST #4408; #441), 6-diamidino-2-phenylindole (DAPI; 1:2000; Sigma-Aldrich; #D9532) and phalloidin-647 (1:2000; ATTO-TEC; #AD647-81) diluted in Intercept (TBS) blocking buffer for 30 min. Cells were washed 4× with TBS in between each step and all steps were performed at room temperature. Validated (for immunofluorescence) and widely cited primary antibodies were used including: p-AKT-T308 (Rb; 1:100; CST; #13038; 1199 citations), p-AKT-S473 (Rb; 1:100; CST; #3787; 452 citations), p-ERK (Rb; 1:500; CST; #9101; 9096 citations), p-S6 (Rb; 1:500; CST; #4858; 1668 citations), and E-cadherin (Ms; 1:500; BD Biosciences; #610181; 1882 citations).

Three-channel immunofluorescence was performed via our previously described ‘extensible labeling’ format [44]. As employed herein, all wells were labeled with a common pair of markers (DAPI and phalloidin-647), whereas subsets of wells were labeled with one of four variable phospho-signaling markers: p-AKT-T308, p-AKT-S473, p-ERK, or p-S6. This design enables systematic profiling of phospho-signaling dynamics across distinct cell populations, while maintaining a consistent set of shared marker features that permit downstream computational integration (see Section 2.7 ‘Computational multiplexing’).

### 2.3 Microscopy

Images were captured using a Nikon AX R confocal microscope (Nikon) equipped with a 20× Plan Apochromat air objective (NA 0.75) with resonance scanning and 16× averaging at 2048×2048 resolution. Custom Nikon JOBS protocols were used for automated acquisition of 9 image fields per well, with 3 z-positions per field. Focus was maintained using the perfect focus system with automatic offset detection by image-based autofocusing. The resulting raw data were exported as .nd2 files per well, and maximum projections were computed per image field.

### 2.4 Image analysis

Maximum projected .tif images were imported into CellProfiler [46] for image analysis via custom pipelines. Nuclei masks were generated using the DAPI channel, employing adaptive thresholding and minimum cross-entropy thresholding. Cell body masks were identified using nuclei as seeds for adaptive thresholding and minimum cross-entropy thresholding on phalloidin images. Cytoplasm masks were generated by subtracting nuclei from cell body masks. Cells touching the image borders were excluded from analysis. A total of 1264 quantitative features were measured for the cell body, cytoplasm, and nuclei masks per cell. These included intensity and radial intensity distributions, granularity and texture features for each marker, cell morphology (size, shape), and cell spatial context information per cell, such as neighboring cell counts and percentages of cell boundaries in contact with adjacent cells.

To analyze the subcellular distributions of fluorescence markers, additional segmentation steps were applied to p-AKT images. For the p-AKT-T308 images, both small, high-intensity puncta and larger membrane ruffles were segmented within the cell body. For p-AKT-S473 images, only high-intensity membrane ruffles were segmented. These regions were quantified independently to capture distinct localization patterns, with all subcellular object values assigned to the parent cell. Pairwise pixel intensity correlation analysis (Pearson’s correlation coefficient) was also performed between segmented AKT regions and corresponding DAPI and F-actin channels to assess colocalization and spatial associations between AKT signaling and nuclear or cytoskeletal structures.

### 2.5 Experimental replication

All experiments, including cell plating, EGF stimulation, immunolabeling, and imaging, were performed in two independent biological replicates. Each replicate included the full panel of EGF stimulation timepoints and antibody labeling conditions.

### 2.6 Data analysis

Data analysis was performed using custom pipelines in Konstanz Information Miner (KNIME, v.4.7.3) [47], Python [48] and R [49]. Python analyses relied on ‘pandas’ [50], ‘numpy’ [51], ‘scikit-learn’ [52], ‘matplotlib’ [53], and ‘seaborn’ [54] packages.

#### 2.6.1 Quality control

A two-stage quality control strategy was implemented in KNIME using a custom interactive workflow to remove segmentation artefacts as well as cells visually identified as multinuclear, dying or mitotic – whose altered signaling responses we do not seek to assess herein. In the first stage, mean nuclear DAPI intensity was plotted against cytoplasmic area to identify and exclude bright DAPI aggregates and objects with no associated cytoplasm. In the second stage, morphology, DAPI, and F-actin features were projected into a t-SNE embedding. Clusters corresponding to mitotic and multinucleated cells were identified through image inspection and thereafter excluded. Of the original 73,937 segmented objects, 64,580 cells were retained for analysis (87%).

#### 2.6.2 Data transformation, normalisation, and replicate handling

Feature values were transformed on the basis of their underlying distributions. Most features were log_2_-transformed; texture features containing zero values (e.g., *TextureContrast*, *TextureVariance*) were square-root transformed. Features with approximately normal distributions (e.g., *FormFactor*) were left untransformed. After transformation, all features were z-score normalized per plate for common features (e.g., DAPI, phalloidin, morphology, context features), and per antibody panel for variable phospho-marker features.

Due to the high reproducibility observed between biological replicates (Fig. S2A), singlecell data were combined for all downstream analyses. Prior to pooling, replicate datasets were normalized independently using the procedure described above. After pooling, all features were re-normalized to ensure consistency across the integrated dataset.

### 2.7 Computational multiplexing

To computationally infer missing marker features across different antibody panels, least absolute shrinkage and selection operator (LASSO) regression models [55] were implemented using the ‘glmnet’ R package [56]. A shared set of 540 common features, including morphology, spatial context, DAPI, and F-actin measurements, was used as input to predict each of 241 marker-specific features (intensity, granularity, texture, and radial distribution) per signaling marker. Models were trained separately for each feature, for each signaling marker using stratified 70/30 train-test splits. Ten-fold cross-validation (cv.glmnet, alpha=1, measure=“deviance”) was performed on the training set to determine the optimal regularization parameter (lambda.min), which was then used to evaluate the prediction performance on the held-out test set. One-hot encoded treatment timepoints were included as unpenalized features to account for condition-specific effects. Prediction performance was assessed using explained variance (*R*^2^) of measured versus predicted features in held-out test data. Predicted granularity features were excluded from the final computationally multiplexed dataset due to low predictive accuracy. A full list of input and predicted features is provided in Supplementary Table S1.

### 2.8 Dimensionality reduction

To construct an E/M landscape, a low-dimensional representation of cell morphology and cell spatial context was created using Uniform Manifold Approximation and Projection (UMAP) [57]. A total of 64,580 cells were included. 21 morphological and spatial context features were subjected to principal component analysis (PCA), retaining components that explained 97% of total variance. The resulting seven PCA components were then input into UMAP (neighbors=200, min_dist=1, metric=’euclidean’) to produce a two-dimensional embedding. Features used to generate the E/M landscape are listed in Supplementary Table S2.

To construct a low-dimensional signaling landscape, cells were first stratified by EGF stimulation timepoint and uniformly downsampled resulting in 12,320 cells per group (61,600 in total). Predicted features from computational multiplexing were filtered to retain only those with *R*^2^ ≥ 0.5 across held-out test data, resulting in 476 well-predicted features. Extreme outlier cells were then excluded per treatment group using an interquartile range (IQR) filter (±5 × IQR) applied independently to each predicted feature. This resulted in a final dataset of 59,092 cells. The 476 predicted features were embedded into a two-dimensional signaling state space using PHATE (t=20, decay=40), implemented via the ‘phate’ [58] Python package. Features used to generate the signaling landscape are listed in Supplementary Table S2.

### 2.9 Pseudotime inference

Pseudotime inference was performed using the ‘scFates’ [59] Python package, which models trajectories as principal graphs learned in low-dimensional space.

To infer an EMT pseudotime trajectory, a principal curve was fitted to the two-dimensional E/M UMAP embedding using the ElPiGraph algorithm [60] with 20 nodes and epg_mu=100. A soft assignment matrix, allowing probabilistic mapping of cells to graph segments, was generated using sigma=1, lam=0.01, and n_steps=1. The root node was manually assigned to the node located in the most epithelial region of the UMAP embedding, and pseudotime values were computed using 100 mapping iterations.

To infer signaling pseudotime in the computationally multiplexed signaling landscape, curve fitting was performed separately for control and EGF-treated cells in the two-dimensional PHATE embedding. For the control group, a curve with 10 nodes was fitted (epg_mu=0.2), and for treated cells, 30 nodes were used. In both cases, soft assignment was applied using sigma=0.001, lam=1, n_steps=1. The resulting principal curves were connected to form a unified trajectory. The root node was manually assigned as the node within the control group that was most distant from the EGF-treated cells. Pseudotime was then computed as described above.

### 2.10 Representative cell selection and E/M state classification

To quantitatively define and visualize representative single cells across the EMT trajectory, pseudotime values were first binned into 10 equally spaced intervals. Within each bin, representative cells were selected by identifying the center of density in UMAP space and extracting the 10 nearest neighbors to that centroid using Euclidean distance. Images were cropped to 200×200 pixels with the representative cell centroid used as the center of the crop.

To classify cells as epithelial- or mesenchymal-like cells, EMT pseudotime values were min-max scaled 0–1 and thresholds were set at the 10th and 90th percentiles. Cells in the bottom decile were assigned to the ‘Epithelial’ category, and those in the top decile to the ‘Mesenchymal’ category. All other cells were labeled ‘Uncategorized’. Representative cell images were computed within these classes as above for visualization.

### 2.11 Feature smoothing over signaling pseudotime

Smoothed signaling feature trajectories were computed separately for epithelial and mesenchymal cell categories using generalized additive models (GAM) via the ‘pygam’ [61] Python package. For each subset, feature values were modelled as a function of signaling pseudotime using a linear GAM with 5 spline terms and a smoothing penalty of 0.1. Models were fit on individual cells, and smoothed feature trajectories were then predicted across 100 evenly spaced pseudotime intervals. Resulting vectors were normalized to their value at the start of pseudotime for subsequent visualization as well as for correlation analyses.

### 2.12 Pseudotime feature correlation analysis

Spearman’s correlation (*ρ*) matrices were computed independently for epithelial and mesenchymal cells using the normalized, GAM-smoothed signaling features. Correlation differences between states were calculated by subtracting mesenchymal correlation values from epithelial values. Hierarchical clustering was performed using Ward’s linkage method.

Feature pairs were classified as ‘formed’ if correlations were weak in epithelial (|*ρ*| *<* 0.5) but strong in mesenchymal cells (|*ρ*| ≥ 0.5), and ‘broken’ if the inverse pattern was observed. Pairs that did not meet either criterion were considered ‘unchanged’. This threshold-based approach enabled us to identify signaling relationships that were gained or lost across states, reflecting context-specific rewiring of inter-feature coordination.

### 2.13 Statistical analyses

Transformed intensity and morphological features displayed a normal distribution (Fig. 1L; Fig. S1G). As such, Student’s *t* -tests were used throughout to compute differences between E/M populations or quantified features at specified time points. *P*-value annotations and statistical tests are indicated in figure legends.

**Figure 1:**
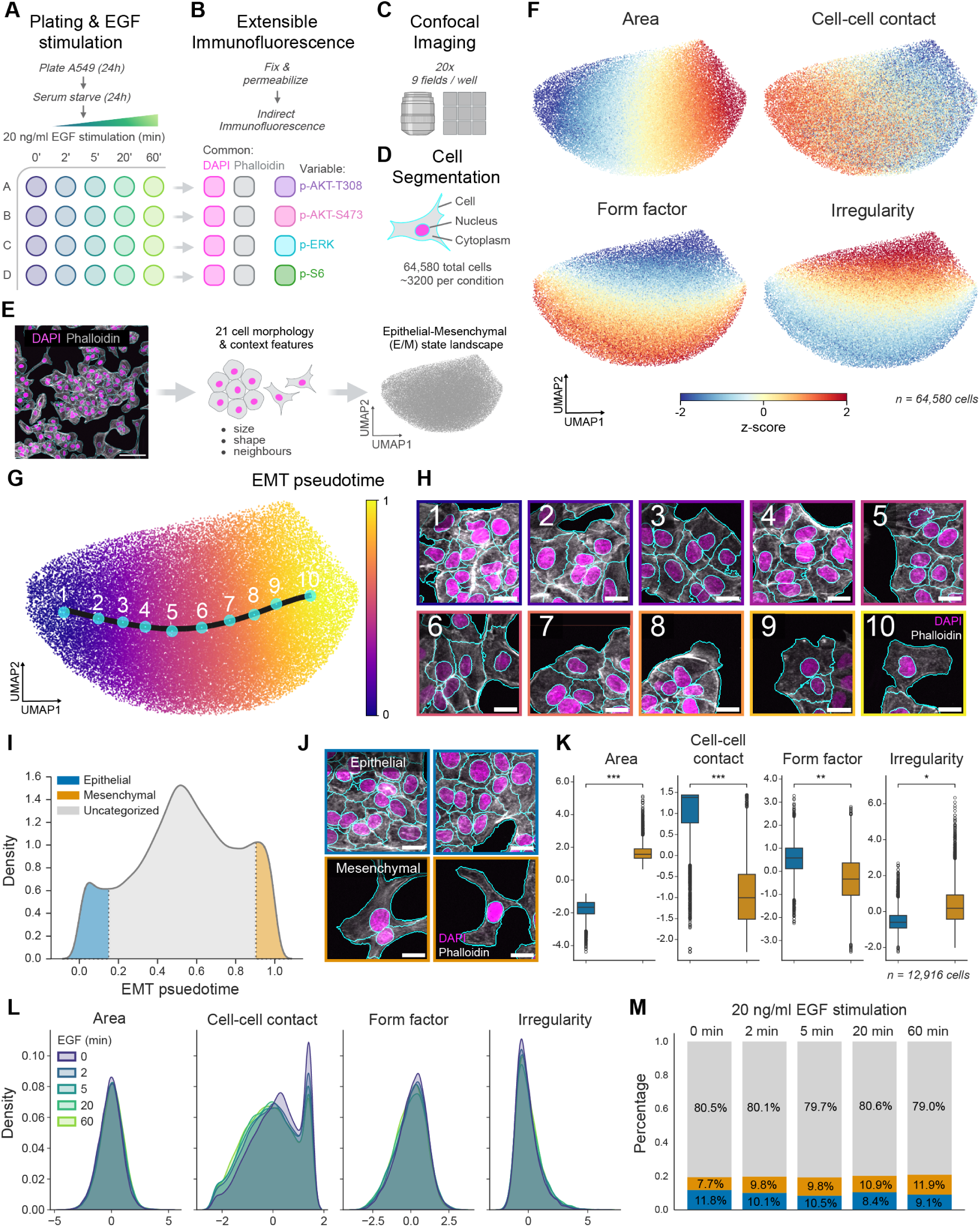
Defining a pre-existing epithelial–mesenchymal landscape in EGF stimulated cells. (A) Experimental workflow. (B) Extensible immunofluorescence labeling. (C) Confocal imaging and (D) cell segmentation. (E) UMAP projection of 21 common morphological and context features defines a continuous landscape of epithelial–mesenchymal (E/M) states. Scale bar = 100 µm. (F) Feature overlays on the landscape reveal an E/M continuum. (G) Pseudotime ordering across the UMAP recapitulates a gradual epithelial–mesenchymal transition. Principal curve shown in black. Ten representative cells identified from binned pseudotime values shown in cyan. (H) Corresponding images of representative cells at pseudotime intervals show progression from epithelial to mesenchymal states. (I) Pseudotime-based classification of epithelial (*<*10th percentile) and mesenchymal (*>*90th percentile) states. (J) Representative epithelial and mesenchymal cell images. (J) Quantification of morphological features in epithelial vs. mesenchymal cells (*n* = 12,916 cells); two-tailed *t* -test (\*\*\**p <* 0.001, \*\**p <* 0.01, \**p <* 0.05). (K) Stability of morphological and context features across the EGF stimulation time course. (L) Proportion of cells classified as epithelial, mesenchymal, or uncategorized across EGF stimulation. All scale bars = 20 µm.

## 3 Results

### 3.1 Morpho-contextual profiling defines the pre-existing epithelial–mesenchymal landscape of A549 cells

To investigate how E/M state influences signaling responses, we applied single-cell quantitative imaging to heterogeneous A549 lung adenocarcinoma cells. Following 24 h attachment and 24 h serum-deprivation, cells were stimulated with 20 ng/mL of EGF for 2, 5, 20, or 60 min prior to fixation (Fig. 1A). Mimicking our previously described ‘extensible immunofluorescence’ (ExIF) labeling and fluorescence marker integration framework [44], all cells were labeled with DAPI and phalloidin (‘common markers’), while subsets were additionally labeled with antibodies against either p-AKT (T308 or S473), p-ERK, or p-S6 (‘variable markers’; Fig. 1B). Confocal imaging was performed, followed by segmentation of single-cell objects (Fig. 1C,D). 1264 quantitative features were measured per cell using CellProfiler [46], spanning whole cell body, cytoplasmic and nuclear domains and including marker intensity, radial intensity distribution, granularity and texture features. After quantitative and visual quality control, the final dataset contained a total of 64,580 cells, constituting ∼3,200 cells per unique treatment and labeling condition.

To delineate ‘pre-existing’ heterogeneity in E/M states, i.e., those that preceded and persisted throughout incident EGF stimulation, we first generated a unified landscape by projecting all cells into a low-dimensional state space using UMAP based on 21 morphological and contextual features measured in all single cells (Fig. 1E). These features captured cell shape, size, spatial context and other properties, enabling construction and visualization of the phenotypic diversity associated with E/M morphologies. Variable (signaling) marker identities (Marker ID) and data from experimental replicate wells (Replicate ID) were well-mixed across the manifold, confirming the robustness and stability of the landscape (Fig. S1A,B).

Inspection of key features across this manifold revealed a clear gradient emblematic of transitions between epithelial-like and mesenchymal-like states. Cells with small areas, high cell–cell contact, and low irregularity—hallmarks of epithelial morphology—were enriched on the left side of the UMAP, whereas larger, more isolated and morphologically irregular mesenchymal-like cells were positioned on the right (Fig. 1F).

To quantify this continuum, we applied pseudotemporal ordering beginning at the epithelial-like end to assign each cell an ‘EMT pseudotime’ score across the manifold (Fig. 1G, landmarks 1–10). Representative cell images quantitatively selected to exemplify pseudotime landmarks 1–10 validated the morphologic progression across pseudotime, from tightly packed cobblestone-like epithelial cells to elongated, irregular mesenchymal cells (Fig. 1H). Identification of morphological or context features that vary with statistical significance over EMT pseudotime (19 of 21 features) enabled quantitative mapping of changes in each feature (Fig. S1C), confirming the visual trends observed above.

By defining the extrema of the EMT pseudotime trajectory (*<*10th or *>*90th percentile of EMT pseudotime, respectively; Fig. 1I), we classified cells into epithelial-like and mesenchymal-like subpopulations and compared their morphological and spatial context profiles. As expected, these groups presented significant differences in key features (Fig. 1J,K; cell area, cell-cell contact, form factor, and irregularity), which is consistent with established E/M phenotypes in A549 cells [62, 63, 64]. To independently validate our morphology-based classification, we repeated the experiment with immunolabeling for E-cadherin, a canonical epithelial marker, and applied the same profiling workflow. E-cadherin expression was enriched at the epithelial end of the manifold, where it localized predominantly to the plasma membrane (Fig. S1D–F). In contrast, mesenchymal-like cells exhibited reduced E-cadherin levels that were predominantly localized internally in punctate compartments. These spatial expression patterns support the validity of our morphology-based classification of epithelial- and mesenchymal-like cells.

Importantly for our study, the E/M states themselves remained highly stable across the EGF stimulation time course, when considering E/M state indicators including the distributions of morphological features, the proportions of epithelial / mesenchymal / intermediate cells (Fig. 1L,M), E-cadherin expression levels and the overall E/M state landscape (Fig. S1G,H). This stability provided a consistent background of E/M state heterogeneity against which to explore how acute EGF signaling responses may differ between pre-existing E/M cell states.

### 3.2 Epithelial–mesenchymal state modulates the magnitude and dynamics of EGF-induced phospho-signaling

Following our characterization and definition (per single cell) of pre-existing heterogeneity in E/M states, we next investigated whether E/M states modulate acute responses to incident EGF stimulation, focusing on four key signaling nodes within the PI3K/AKT and MAPK path-ways: p-AKT (T308 and S473), p-ERK, and p-S6 (Fig. 2A).

**Figure 2:**
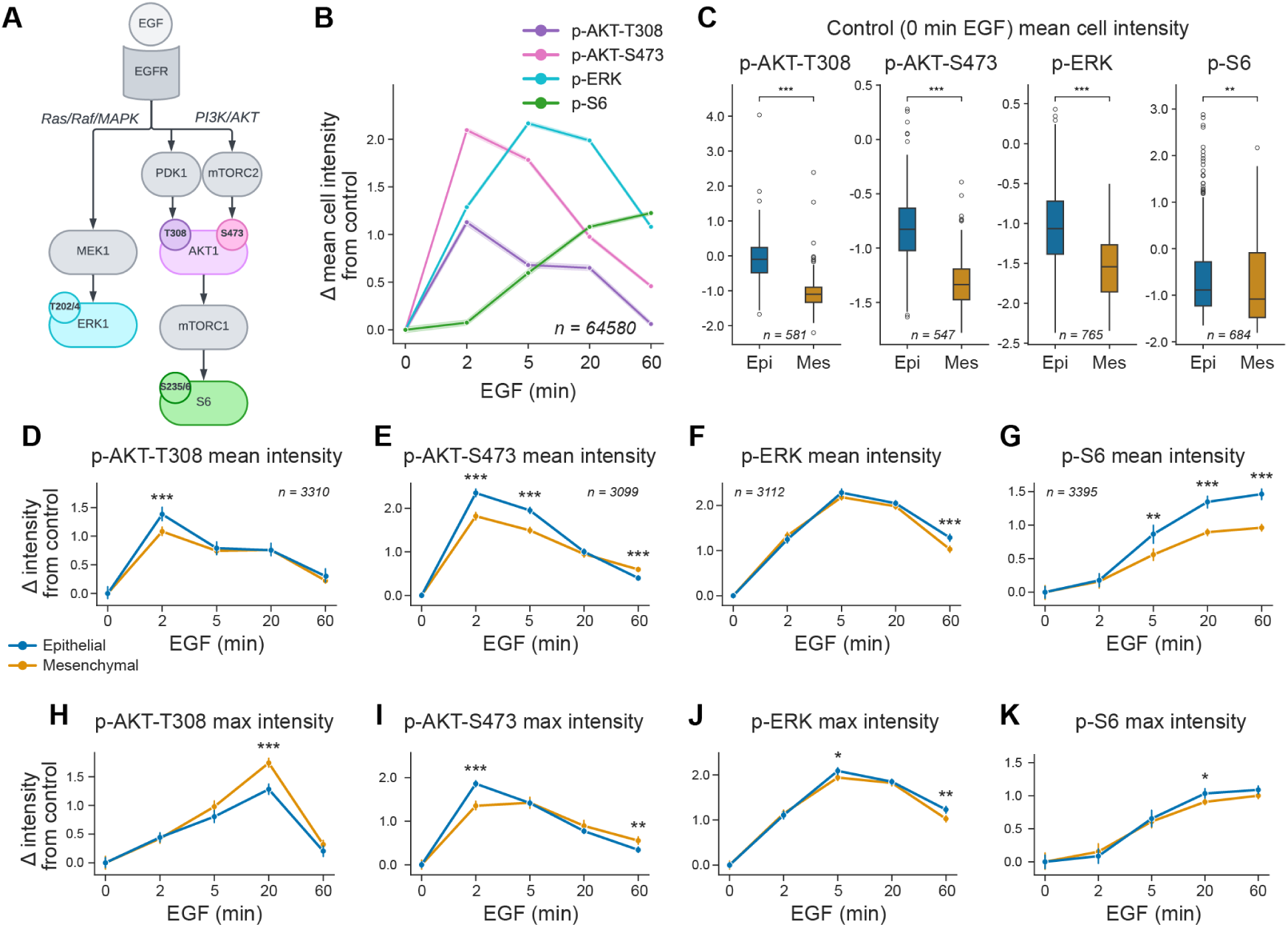
EGF-induced phospho-signaling dynamics differ between epithelial and mesenchymal cells. (A) Schematic of EGFR transduction pathways highlighting key phosphorylation events in the PI3K/AKT and MAPK cascades. (B) Time course of EGF-induced changes in mean phospho-signal intensity relative to control (0 min) (n = 64,580 cells). (C) Basal (0 min EGF) mean cell intensities of phospho-signals in epithelial and mesenchymal cells. Time course of phospho-protein mean (D–G) and maximum (H–K) intensity changes relative to control, grouped by epithelial (blue) and mesenchymal (orange) states. Statistical significance between epithelial and mesenchymal cells was assessed using two-tailed t-tests (***p < 0.001, **p < 0.01, *p < 0.05). Line plots represent the mean ± 95% confidence interval.

We first assessed global EGF-induced responses by quantifying changes in the mean cell intensity of phospho-proteins relative to the unstimulated control (Fig. 2B). All phospho-proteins showed strong activation, with distinct temporal dynamics. p-AKT-T308 and p-AKT-S473 exhibited rapid but transient activation, peaking at 2 min before beginning to decline by 5 min. p-ERK responses peaked later at 5 min and remained moderately elevated at 20 min. In contrast, p-S6 intensity increased gradually over time, reaching maximal levels in this analysis at 60 min. These temporal patterns align with the established topologies of each marker within these canonical pathways, i.e., AKT and ERK activation is rapid and transient while p-S6 reflects downstream integration of sustained mTORC1 signaling [65, 66, 67]. These dynamics were robust and reproducible across biological replicates (Fig. S2A).

Next, we compared baseline (0 min) phosphorylation levels between the previously classified epithelial- and mesenchymal-like cells (Fig. 2C). Mesenchymal cells exhibited significantly lower basal phosphorylation across all markers, suggesting reduced steady-state signaling activity. To assess differences in signaling response dynamics, we normalized intensity values per marker at each timepoint relative to the respective control (0 min) values for each E/M cell state. This revealed distinct temporal signaling profiles between epithelial- and mesenchymal-like cells. Epithelial cells demonstrated stronger early responses in p-AKT-T308 (2 min, Fig. 2D) and p-AKT-S473 (2–5 min, Fig. 2E), indicating rapid and transient AKT activation. In contrast, mesenchymal cells displayed a more prolonged p-AKT-S473 response that remained elevated at 60 min, suggesting sustained PI3K pathway activity. For p-ERK, the overall response dynamics were similar between cell states, but epithelial cells retained higher levels at 60 min (Fig. 2F). p-S6 responses were consistently stronger and more sustained in epithelial cells, continuing to rise at 60 min, whereas mesenchymal cells reached a plateau earlier (Fig. 2G).

Since phospho-signaling can be differentially localized within cells, with functional implications related to locally-available substrates [20, 68], we next compared maximum intensity per-cell signaling values (Fig. 2H–K), emblematic of peak local subcellular signal concentrations, to the mean (whole) cell signaling values detailed above. While epithelial cells exhibited stronger peak mean p-AKT-T308 intensity than mesenchymal cells at 2 min, maximum intensities peaked later at 20 min in both populations with maximum local signal concentration now far higher in mesenchymal cells (Fig. 2H). p-AKT-S473 also saw mesenchymal cells exhibit delayed peak maximum intensity (5 min) compared to mean p-AKT-S473 levels (2 min) (Fig. 2I), while epithelial mean and maximum intensity values peaked concurrently at 2 min. Maximum intensity p-AKT-S473 values were again sustained (at 60 min) at higher levels in mesenchymal than in epithelial cells. Collectively, these differences suggest that while epithelial cells show robust and spatially uniform AKT activation, mesenchymal cells may rely on more focalized, sustained activation, potentially reflecting differential subcellular localization or compartmentalized signaling. This indication is further explored below. Differences in dynamics and magnitude were not observed between mean and maximum intensities of p-ERK or p-S6 (Fig. 2J,K). Relative intensity values (not scaled to starting control (0 min) intensities) for all phospho-proteins across the time course are shown in Fig. S2B.

Together, these findings reveal significant and diverse cell state-dependent differences in phospho-signaling magnitudes, dynamics, and localizations. Epithelial cells display higher basal phosphorylation and more transient yet robust EGF-induced activation, while mesenchymal cells exhibit lower basal activity but sustain longer and potentially more locally concentrated signaling responses, particularly within the PI3K/AKT pathway.

### 3.3 EGF signaling dynamics vary continuously across the epithelial–mesenchymal continuum

Beyond comparing discrete epithelial and mesenchymal states, we asked whether phospho-signaling responses exhibit a continuum of regulation across the E/M spectrum. To do so, we assessed the mean and maximum phospho-intensities for all markers as a function of EMT pseudotime, grouping cells by actual EGF stimulation time (Fig. S3). This approach allows us to examine whether phospho-signaling dependence on E/M state varies in a graded manner or undergoes discrete switches, potentially related to specific E/M states.

Overall, we observed evidence of a continuous spectrum of responses across E/M states for most markers, aligning with the trends seen in the discrete states. p-ERK responses remained relatively stable across the EMT continuum in both mean and maximum intensity, supporting findings that ERK variation has little dependence on E/M heterogeneity from our discrete state comparisons (Fig. S3A). p-S6 responses exhibited a gradual decline over EMT pseudotime, implying coupling to continually varying features of the E/M state spectrum (Fig. S3B).

p-AKT signaling again exhibited the most complex behavior, with state-dependent response dynamics varying depending on stimulation time. Early EGF-induced mean p-AKT-S473 responses (2–5 min) declined progressively along the EMT spectrum, whereas the 60 min response increased towards mesenchymal-like cells (Fig. S3C). Similarly, the maximum intensity p-AKT-T308 responses at the 5 and 20 min timepoints increased over EMT pseudotime, whereas other timepoints decreased (Fig. S3D). Interestingly, maximum p-AKT-S473 intensity at the 20 and 60 min timepoints showed a potential switch-like behavior (at ∼12 in EMT pseudotime), suggesting a sharp shift in AKT regulation beyond a certain EMT pseudotime threshold (Fig. S3E). Overall, these findings suggest that signaling response regulation evolves along a continuum of EMT states. The largely progressive responses of AKT, ERK, and S6 phosphorylation indicate that cells at different EMT stages perceive and respond to EGF in a graded manner. Further characterization of EM state-dependent switch-like behaviors in p-AKT-S473 responses to prolonged EGF stimulation is needed.

Considering the UMAP projection of E/M state space, EMT pseudotime predominantly followed the X-axis (UMAP dimension 1). As such, our analyses thus far have focused on heterogeneity along this axis as a potential determinant of signaling responses. However, considerable variation in cell morphology was also observed along the Y-axis (UMAP dimension 2) of the E/M state space, particularly in shape-related features such as form factor and irregularity (Fig. 1E). To assess whether this alternate axis also contributes to variation in EGF-induced signaling, we applied an unsupervised k-means clustering approach in the 21-dimensional space defined by all E/M morphology and context features, defining 20 phenotypic clusters spanning the landscape. Unlike pseudotime binning, this approach preserves variation across both UMAP axes, allowing us to explore potentially distinct signaling behaviors in different regions of the landscape. Clusters were subsequently ordered based on either their mean X-axis (Fig. S4A) or mean Y-axis (Fig. S4B) coordinates in the UMAP-defined E/M cell state space.

When clusters were ordered along the X-axis, phospho-signaling responses recapitulated the trends observed with EMT pseudotime (as expected), showing progressive patterns of signaling responses depending on the X-axis position within the manifold (Fig. S4C). In contrast, ordering clusters along the Y-axis revealed less coherent or directional trends in signaling (Fig. S4D), indicating that this axis contributes less prominently to determining structured differences in the EGF signaling response. Given the distributions of cell morphological and spatial context features in the E/M landscape (Fig. 1F), this highlights that cell size and spatial context features (e.g., cell-cell contact and cell crowding; predominantly X-axis-associated) exert more influence over signaling dynamics than cell shape features (predominantly Y-axis-associated). This is consistent with prior reports highlighting the importance of local cell density in modulating signaling responses and other cellular behaviors [9, 69].

Together, these findings provide further support for graded, state-dependent modulation of EGF signaling across the EMT spectrum and demonstrate that both continuous (pseudotime-based) and discrete (cluster-based) approaches to defining E/M state reveal consistent EMT-associated variation in signaling pathway activation.

### 3.4 AKT phosphorylation exhibits spatially and temporally distinct localization patterns across cell states

Differences in the magnitude and dynamics of mean and maximum intensity p-AKT responses across epithelial- and mesenchymal-like cells (Fig. 2, Fig. S2) suggested underlying variation in subcellular localization. Visual inspection of stimulated populations revealed that p-AKT-T308 primarily adopted a punctate distribution, particularly at 5 and 20 min post-EGF stimulation, before declining by 60 min (Fig. 3A). This pattern is consistent with AKT recruitment to endomembrane signaling compartments such as endosomes or lysosomes [70, 71, 72, 73, 74, 75]. We also found evidence that p-AKT-T308 localized at membrane ruffles in a small number of cells (Fig. S5A).

**Figure 3:**
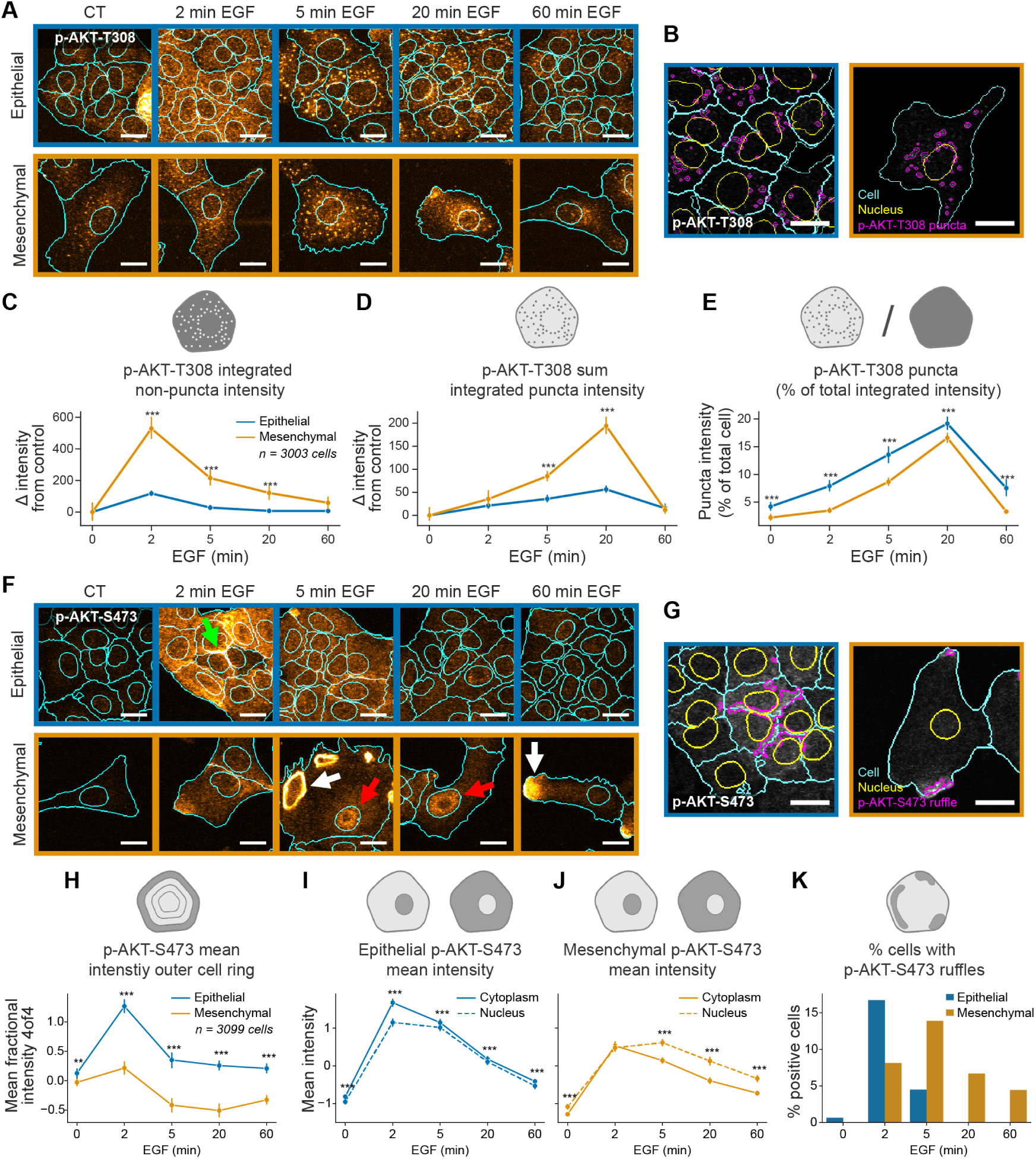
EGF-induced phospho-signaling dynamics differ between epithelial and mesenchymal cells. (A) Representative immunofluorescence images of EGFstimulated p-AKT-T308 in epithelial and mesenchymal cells. (B) Segmentation of p-AKT-T308 signal into punctate and non-punctate compartments. Time course of (C) integrated nonpuncta p-AKT-T308 intensity, (D) sum of integrated puncta p-AKT-T308, and (E) proportion of punctate signal compared to total signal, comparing epithelial and mesenchymal states. (F) Representative immunofluorescence images of EGF-stimulated p-AKT-S473. Green, white, and red arrows indicate uniform membrane, ruffle-localized, and nuclear signal respectively. (G) Segmentation of p-AKT-S473 signal into ruffle regions. (H) Time course of p-AKT-S473 mean outer membrane (outer ring) signal in epithelial and mesenchymal cells. (I–J) Quantification of nuclear and cytoplasmic p-AKT-S473 mean intensities in epithelial (I) and mesenchymal (J) cells. (K) Percentage of cells exhibiting p-AKT-S473 ruffles across time points and E/M states. Statistical significance between epithelial and mesenchymal cells was assessed using a two-tailed t-test (\*\*\**p <* 0.001, \*\**p <* 0.01, \**p <* 0.05). Line plots represent the mean ± 95% confidence interval. All scale bars: 20 *µ*m.

Focusing on the endomembrane spatial patterns, we performed intracellular segmentation of p-AKT-T308 signals into punctate and non-punctate regions (Fig. 3B). Punctate signal intensities peaked later than non-punctate signals in both cell states (Fig. 3C,D), mimicking the temporal trends seen with mean versus maximum intensity p-AKT-T308 signals (Fig. 2), and now clearly indicating delayed compartmentalized activity. Although the temporal profile was similar between E/M states, mesenchymal cells displayed far higher integrated (total) puncta-associated signal intensity, likely due to their larger size and more numerous puncta (Fig. S5B). Interestingly, the proportion of total p-AKT-T308 signal localized to puncta was actually higher in epithelial cells (Fig. 3E), indicating proportionally stronger endomembrane-associated signaling in epithelial states. These findings suggest that distinct pools of p-AKT-T308 (punctate and diffuse) operate with different magnitudes and dynamics across E/M states.

In contrast, p-AKT-S473 displayed distinct localization patterns compared to the T308 site (Fig. 3F). Three spatial distributions were observed: uniform membrane, ruffle-associated, and nuclear (Fig. 3F, green, white, and red arrows, respectively). To resolve the dynamics of these spatially organized pools, we quantified p-AKT-S473 signal at the membrane (outer-ring), nucleus, cytoplasm, and ruffles (Fig. 3G).

Both E/M cell states exhibited similar baseline outer-ring intensities (Fig. 3H). Upon stimulation, epithelial cells showed a sharp 2-minute increase, while mesenchymal cells had a weaker response. By 5 min, outer-ring intensity declined in both states, and continued to decline in mesenchymal cells through 60 min, suggesting re-localization from the membrane. In contrast, epithelial membranous p-AKT-S473 returned to baseline levels.

We next compared nuclear and cytoplasmic p-AKT-S473 intensities. In epithelial cells, cytoplasmic signal exceeded nuclear intensity at early timepoints, with both compartments declining in parallel thereafter (Fig. 3I). In mesenchymal cells, however, p-AKT-S473 signal continued to increase in the nucleus until 5 min, unlike cytoplasmic signals that peaked at 2 min (Fig. 3J). Nuclear signal levels also declined more slowly in mesenchymal cells, indicating enhanced nuclear retention. These compartment-specific patterns were further supported by pixel-level colocalization analyses: epithelial cells exhibited a transient increase in correlation between cytoplasmic p-AKT-S473 and F-actin, peaking at 2 min, while mesenchymal cells showed consistently weaker correlation (Fig. S5B). Conversely, nuclear p-AKT-S473 displayed stronger and more sustained correlation with DAPI in mesenchymal cells, with a peak at 20 min and elevated levels maintained at 60 min (Fig. S5C). In contrast, epithelial cells showed a decline in nuclear correlation at 2 min, consistent with redistribution of signaling to the cytoplasm (Fig. S5D).

Lastly, we quantified the percentage of cells with p-AKT-S473-positive membrane ruffles (Fig. 3K). Epithelial cells showed a transient increase in ruffle-positive cells at 2 min, but by 5 and 20 min, a greater proportion of mesenchymal cells displayed ruffle-localized signal. This pattern was sustained in mesenchymal cells through 60 min, suggesting that ruffle structures may support prolonged p-AKT-S473 signaling uniquely in mesenchymal states.

Taken together, these findings reveal that epithelial and mesenchymal cells differ not only in the magnitude and dynamics of AKT signaling, but also in the spatial organization of differential AKT site-specific phosphorylation in response to EGF. For p-AKT-T308, both cell states exhibited an early diffuse response followed by later accumulation in punctate endomembrane-associated compartments. Epithelial cells displayed a higher proportion of total T308 signal in puncta early in the response, indicating a more rapid localization of AKT to endomembrane structures.

p-AKT-S473 displayed distinct dynamics, with epithelial cells exhibiting a sharp early increase in membrane-localized signal, followed by redistribution within the cytoplasm and nucleus, where intensities remained balanced over time. Mesenchymal cells showed a weaker overall response but preferentially retained p-AKT-S473 in the nucleus and at membrane ruffles, showing more sustained and spatially focused signaling. These differences highlight that E/M state modulates not just the strength but also the subcellular routing of activated AKT, potentially shaping divergent downstream transcriptional outcomes.

### 3.5 Computational multiplexing reconstructs unified multi-marker signaling profiles from separate labeling panels

For a more comprehensive analysis of EGF-induced multi-molecular signaling responses, we next aimed to integrate the independently labeled phospho-signaling data across all cells. We designed a workflow termed ‘computational multiplexing’ that leverages our previously described ExIF framework, where standard 3- or 4-channel IF is performed using different antibody panels in parallel [44] (Fig. 4A). Each independently labeled cell population contains common markers—present in all panels—and variable markers, which differ between panels. This ‘extensible labeling’ process generates non-integrated datasets consisting of multiple 3-plex labeled cell populations, with two common markers (DAPI, phalloidin) and one variable (phospho-signaling) marker per labeling panel (Fig. 4B; note, the schematically depicted structure reflects data for a single timepoint).

**Figure 4:**
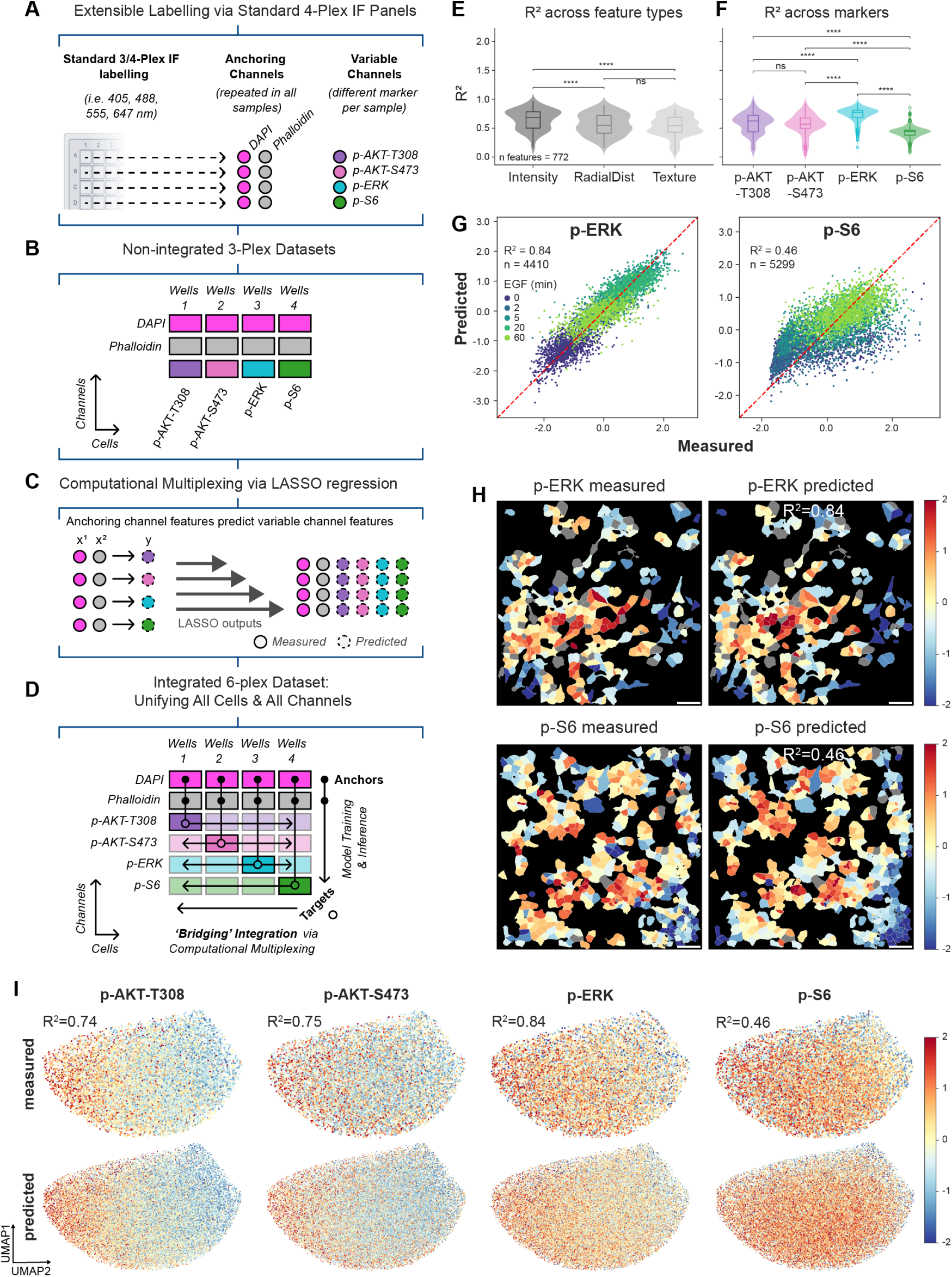
Computational multiplexing integrates independently labeled phospho-signaling data into a unified single-cell dataset. (A) Extensible labeling strategy using standard 3–4 channel immunofluorescence (IF) panels with shared anchoring channels (DAPI, phalloidin) and variable channels (p-AKT-T308, p-AKT-S473, p-ERK, p-S6). (B) Non-integrated 3-plex datasets generated from parallel IF labeling. (C) Computational multiplexing framework. (D) Predicted features are used to unify all cells and markers into an integrated dataset. (E) Prediction performance across feature categories and (F) marker. *P*-values as-sessed using two-tailed t-test with Bonferroni correction (\*\*\**p <* 0.001, \*\**p <* 0.01, \**p <* 0.05). (G) Predicted versus measured mean intensities for p-ERK and p-S6. (H) Spatial mapping of measured and predicted marker intensities in a representative field of view. Scale bars = 100 *µ*m. (I) Distribution of measured and predicted marker intensities for each phospho-protein projected onto the EMT UMAP.

To computationally integrate these separate populations, we trained numerous multi-linear models using LASSO regression [55] to predict all features for each variable (phospho-signaling) marker, using features derived from common markers and other shared cell features (e.g., cell morphology, spatial context) (Fig. 4C). Because the input feature set is shared across all cells, these regression models can then be applied across the entire dataset to predict all (missing) variable marker features within each cell. This creates a fully integrated, multi-molecular dataset (Fig. 4D).

In this case, the common feature set comprised 539 DAPI and F-actin intensity, granularity, texture, and radial distribution features, as well as cell morphology and spatial context information for each cell. The predicted variable marker features (i.e., the outputs of regression models) comprised 241 intensity, texture, and radial-distribution features each for p-AKT-S473, p-AKT-T308, p-ERK, and p-S6. While granularity features were included as input features during model training, they were excluded from the final output feature set due to poor prediction performance during optimization. For each variable marker feature, a single LASSO model was trained using the common feature set as input and ground-truth feature values (from real labeling) as model targets. Prediction accuracy was assessed using *R*^2^, comparing ground-truth and predicted feature values in held-out test sets. After training, each model was applied to the whole dataset, thereby generating the computationally multiplexed dataset.

Comparisons of prediction performance across all variable (target) markers, subset by feature category, revealed that intensity features were best predicted, followed by radial-distribution and texture features (Fig. 4E). We then assessed feature predictions grouped by variable (phospho-signaling) marker (Fig. 4F). p-ERK was found to be the best predicted when all features were considered, which matches the findings of a prior report demonstrating that its levels were best predicted by generic stable cell state features after EGF stimulation [9]. Considering mean phospho-signaling marker intensities specifically, single-cell comparisons of measured (using the cell subset with the labeled marker) versus predicted (in all cells not labeled with the marker) values showed varied correspondence across markers (Fig. 4G,H). For instance, p-ERK mean responses were well-predicted across all EGF stimulation times, whereas p-S6 predictions were less accurate. To further validate our computational multiplexing approach, we visualized the distribution of measured versus predicted mean cell marker intensities across the common E/M state UMAP (Fig. 4I), revealing strong correspondence in predicted signal-strength distributions.

Taken together, computational multiplexing achieves integration of several signaling responses into a unified dataset, enabling exploration of the relationships between E/M states and the complex evolution of multi-molecular phospho-signaling in response to EGF.

### 3.6 Divergent multi-molecular signaling trajectories emerge between epithelial and mesenchymal cells

We next sought to define single-cell positions within the dynamically evolving, multi-molecular signaling response landscape arising from EGF stimulation. We therefore projected a subset of well-predicted signaling features (464 features exceeding 0.5 *R*^2^ for measured versus predicted per-cell values) into a two-dimensional signaling state space using PHATE manifold-learning [58] (Fig. 5A,B). Pseudotime analysis was performed in this multi-molecular ‘signaling landscape’, defining unstimulated cells (0 min) as the pseudotime ‘start’ (Fig. 5C). This enabled reconstruction of multi-signaling progression, which was validated by comparing the closely matched evolution of predicted versus ground-truth signaling features along signaling pseudotime (Fig. 5D).

**Figure 5:**
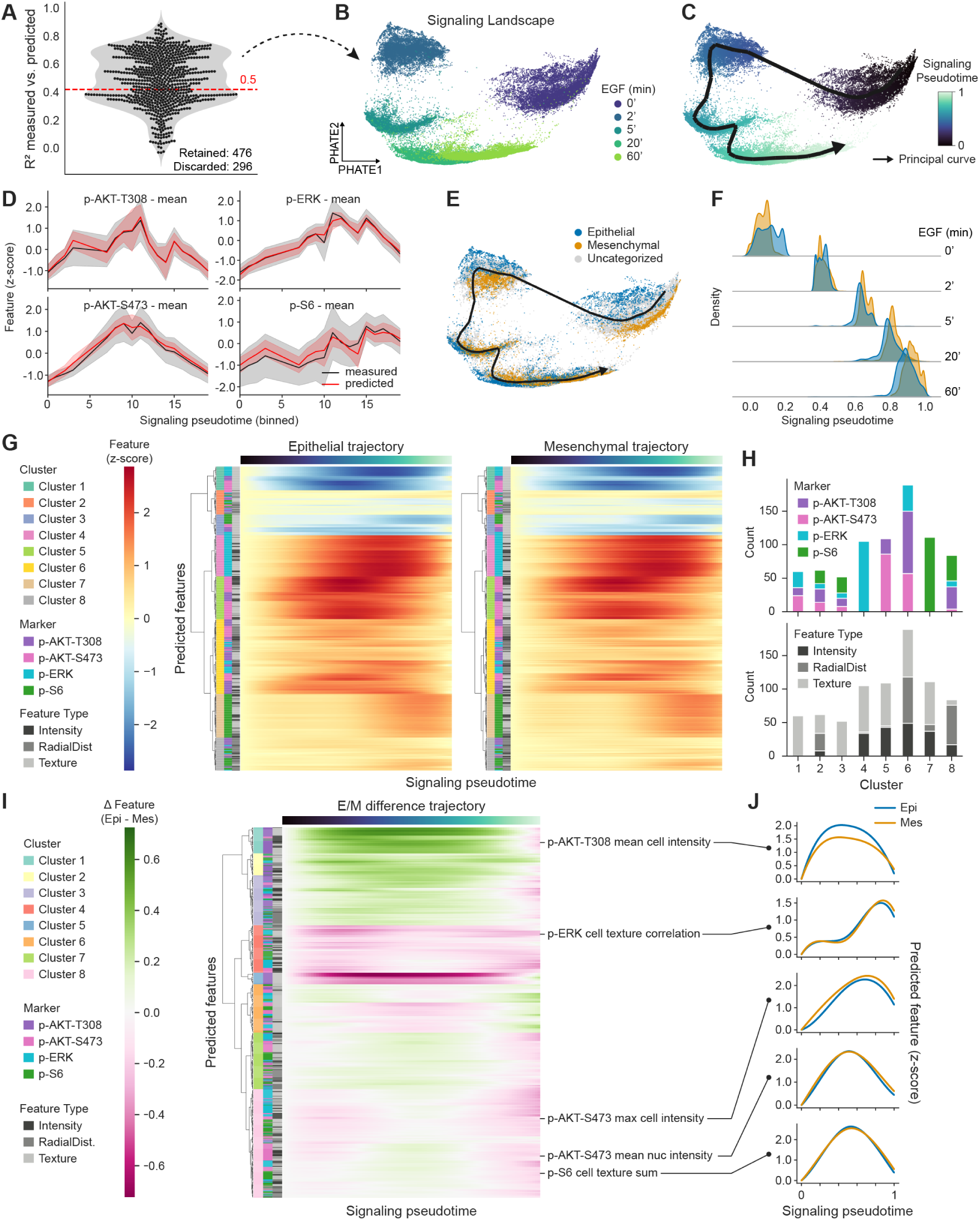
E/M state determines signaling progression through multi-plexed phospho-dynamics. (A) Feature prediction scores (*R*^2^) from LASSO regression models. Dashed line indicates *R*^2^ = 0.5 threshold used to retain well-predicted features. (B) PHATE embedding of computationally multiplexed phospho-signaling features. (C) signaling pseudotime inferred from the PHATE space using unstimulated cells as root. (D) Binned pseudotime values comparing measured and predicted mean intensities of phospho-signals. (E) EMT category overlaid on the signaling landscape. (F) Density plots of epithelial and mesenchymal cells along signaling pseudotime across EGF stimulation timepoints. (G) Heatmaps of smoothed, normalised feature trajectories over pseudotime in epithelial and mesenchymal cells, clustered using Ward hierarchical clustering in the epithelial subset. (H) Cluster composition by marker (top) and feature category (bottom). (I) Difference heatmap computed by subtracting epithelial–mesenchymal (E/M) differences in feature trajectories (Epi – Mes), clustered using Ward method. (J) Examples of smoothed feature trajectories from selected difference clusters.

Focusing on the differences in multi-molecular signal evolution between E/M states (Fig. 5E), we note that at early timepoints (0–2 min) epithelial cells occupied more advanced positions along signaling pseudotime, consistent with their higher signaling baseline and more rapid activation (Fig. 5F). Yet from 5 min onward, mesenchymal cells surpassed epithelial cells along signaling pseudotime, indicating accelerated and sustained signaling activity in this population, potentially linked to elevated AKT activation in the nucleus and/or in ruffles, as evidenced previously (Fig. 3).

To identify features defining these differences, we first clustered signaling features by their temporal trajectories in epithelial cells and applied the same ordering to mesenchymal cells (Fig. 5G). Cluster 5 contained early-responding p-AKT intensity and texture features; Cluster 4 included mid-phase p-ERK features; and Cluster 7 comprised late-responding p-S6 features (Fig. 5H, upper). These dynamics are consistent with our previous analyses of selected intensity features (Fig. 2). Other clusters were more heterogeneous, comprising mixed feature types. Notably, Clusters 1 and 3 consisted exclusively of texture features (Fig. 5H, lower) from all markers, which declined markedly across pseudotime.

To define E/M state-specific differences in signal progression, we computed the difference matrix between epithelial and mesenchymal (from Fig. 5G) trajectories and applied hierarchical clustering to identify groups of phospho-signaling features with similar differences in pseudotemporal dynamics between E/M states (Fig. 5I,J). Here, Clusters 1–3 captured features with stronger early responses in epithelial cells (e.g., mean p-AKT-T308 intensity), while Clusters 4 and 5 included features with greater responses in mesenchymal cells, such as p-ERK texture correlation, a measure of spatial signal regularity. Clusters 7 and 8 were enriched for features elevated in mesenchymal cells at late pseudotime, including maximum and nuclear p-AKT-S473 intensities and p-S6 texture entropy, which reflects the complexity and disorder of signal distribution within the cell. These patterns again support the idea that mesenchymal cells sustain signaling activity over prolonged periods, whereas epithelial cells mount stronger but more transient responses.

Together, these findings reinforce our evidence that E/M state modulates not only the strength, kinetics, duration and intracellular localizations of signaling, but also the coordination of dynamic, multi-molecular response patterns captured through multiplexed analyses.

### 3.7 Epithelial–mesenchymal state reshapes the coordination between phospho-signaling features

To assess how signaling features are interrelated during EGF stimulation in epithelial and mesenchymal-like cells, we examined changes in the correlation structure between all predicted features over signaling pseudotime. Spearman’s correlation matrices were computed separately for each E/M state, hierarchically clustered in the epithelial subset and ordered equivalently in the mesenchymal subset (Fig. 6A). In both cell states, clear modular blocks of correlated features were observed, reflecting coordinated regulation of feature subsets across signaling pseudotime. The global structure of correlations was broadly preserved across states.

**Figure 6:**
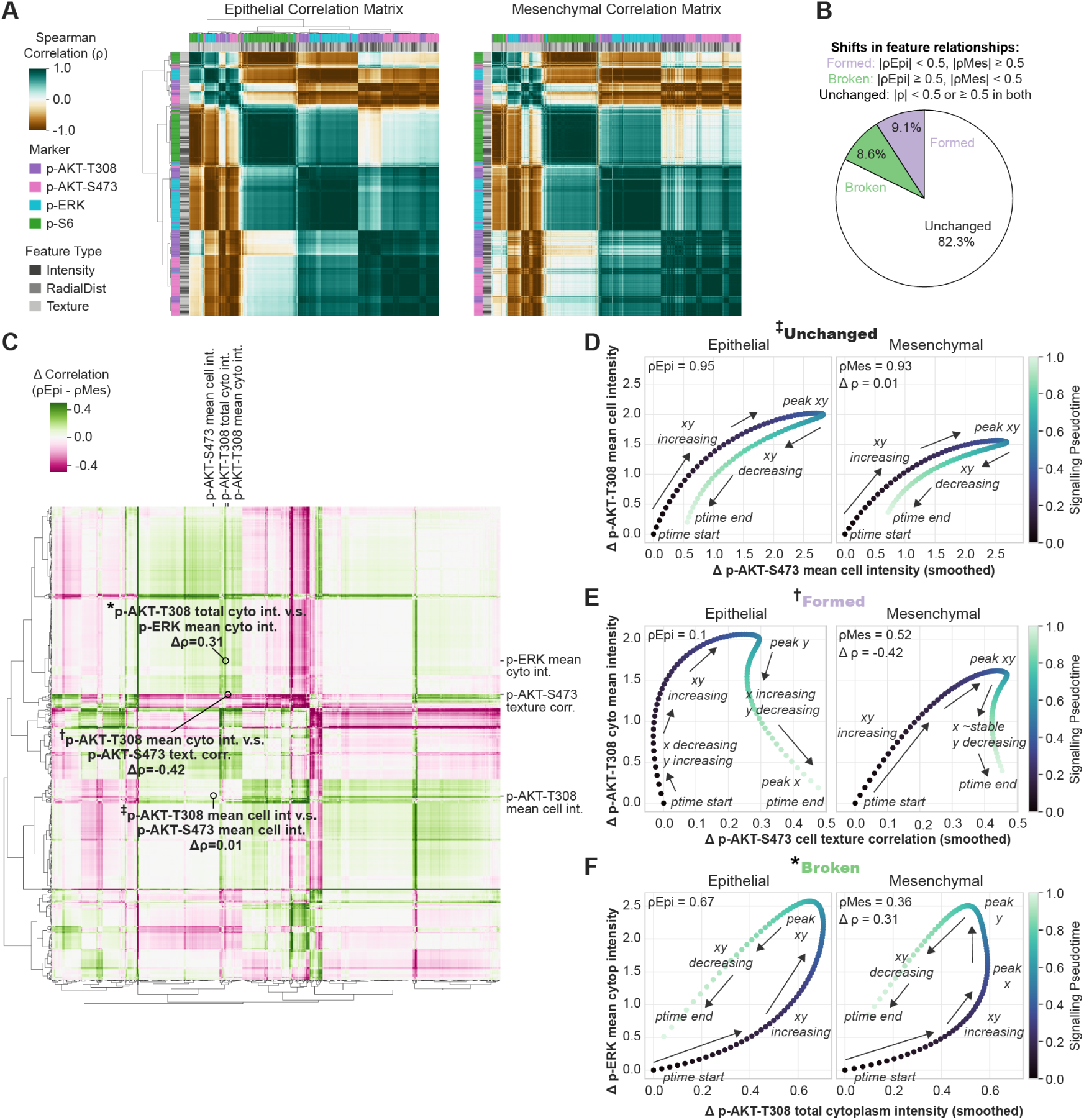
E/M state-specific shifts in feature interdependencies during signaling progression. (A) Spearman correlation matrices of smoothed, normalised signaling features computed separately for epithelial and mesenchymal cells, with hierarchical clustering (Ward) applied to the epithelial matrix and used to order both states. (B) Pie chart showing percentage of feature–feature relationships altered between epithelial and mesenchymal cells. (C) Difference correlation matrix computed by subtracting the mesenchymal matrix from the epithelial matrix. Representative examples of unchanged, formed, and broken relationships are annotated. (D–F) Joint pseudotime scatterplots of representative feature pairs. Features are coloured by signaling pseudotime (dark blue: early; light green: late). Relationships were classified by changes in Spearman correlation between states as per panel B.

Comparing epithelial and mesenchymal matrices, we categorized pairwise relationships as either ‘unchanged’ (no strong change in epithelial and mesenchymal cells), ‘formed’ (weak in epithelial but strong in mesenchymal) or ‘broken’ (strong in epithelial but weak in mesenchymal) (Fig. 6B). This confirmed that the majority of relationships (82.3%) were maintained between cell states, consistent with overall conservation of signaling pathway architecture and inter-feature dependencies. Nonetheless, a sizable fraction of relationships were state-specific, with 9.1% formed and 8.6% broken between epithelial and mesenchymal cells. Of the analyzed markers, p-AKT-T308 features were most involved in both types of relationship changes (total 35.1% changed), suggesting strong divergence in its regulation across E/M state relative to, for example, p-AKT-S473 (total 18.5% changed) (Fig. S6A). Texture features were also enriched in relationship changes, indicating that alterations in subcellular marker distributions were more frequent than changes in overall signal intensity across E/M states (Fig. S6B).

To quantify the shifts in feature relationships between cell states, we then computed a correlation difference matrix (Fig. 6C). Clustering within this matrix revealed groups of signaling features that exhibited consistent shifts in their interdependencies between epithelial and mesenchymal states. These shifts were organized into discrete modules, suggesting that subsets of signaling features undergo coordinated modulation depending on cell state. To illustrate the spectrum of observed changes we selected three feature-pairs representing unchanged, broken or formed relationships (Fig. 6C, circles). Joint pseudotime trajectory scatterplots were used to visualize the co-progression of the paired features, visually capturing their relative temporal dynamics and inter-relationships (Fig. 6D–F).

As an unchanged (between E/M states) example, mean p-AKT-S473 and p-AKT-T308 intensities remained strongly correlated in both states (Fig. 6D). Both features increased with pseudotime, peaking at the same time but with p-AKT-S473 reaching a higher maximum, before both decreased. Despite a similar overall correlation structure, epithelial cells exhibited larger peak responses, while mesenchymal cells maintained higher levels of both features at the end of pseudotime.

A formed relationship was observed between p-AKT-S473 texture correlation (a feature quantifying spatial regularity) and mean cytoplasmic p-AKT-T308 intensity (Fig. 6E). In epithelial cells, these features were uncorrelated: p-AKT-T308 rose early, while p-AKT-S473 texture initially declined, plateaued, and then gradually increased, reaching its peak at the end of pseudotime. In contrast, mesenchymal cells exhibited a moderate positive correlation, with both features rising concurrently and peaking at similar pseudotime values. Notably, the late-phase increase in p-AKT-S473 texture observed in epithelial cells was absent in mesenchymal cells, suggesting a more temporally constrained spatial signaling response.

A broken relationship emerged between total p-AKT-T308 and mean p-ERK intensity in the cytoplasm (Fig. 6F). In epithelial cells, these features showed a coordinated rise, with p-AKT-T308 peaking at a similar time to p-ERK. In mesenchymal cells, p-AKT-T308 peaked earlier and at lower intensity, followed by a continued increase in p-ERK, disrupting their temporal alignment. Both features eventually declined, but mesenchymal cells retained elevated levels at late pseudotime, consistent with their sustained signaling profile.

Taken together, our analyses reveal that epithelial and mesenchymal cell states not only exhibit distinct magnitudes, temporal dynamics, and localizations of EGF-induced phospho-signaling but also differ in the subcellular coordination and integration of these signals. By leveraging computational multiplexing and trajectory inference, we demonstrate that E/M state heterogeneity underlies much of the observed signaling variability, with epithelial cells showing stronger, more rapid responses, and mesenchymal cells displaying slower, more sustained and spatially focused signaling. These findings underscore the critical role of pre-existing cell states in shaping how tumor cells process oncogenic cues, likely leading to divergent functional outcomes in response to the same extracellular signal.

## 4 Discussion

Signaling responses to growth factor stimulation are often highly variable across tumor cell populations, yet the extent to which this signaling heterogeneity is shaped by pre-existing differences in cell state remains poorly resolved [5, 7]. Using quantitative immunofluorescence, morpho-contextual profiling, and computational multiplexing downstream of EGF stimulation in A549 NSCLC cells, we demonstrate that pre-existing E/M states modulate distinctive signaling phenotypes, with characteristic differences in magnitude, dynamics, spatial localization, and coordination of dynamic multi-molecular signaling activity. These findings reveal that E/M state is a major predeterminant of heterogeneous phospho-signaling responses that, in isolation, may otherwise be misinterpreted as stochastic noise. By showing that E/M states shape how cells decode and respond to oncogenic ligand signals, we highlight the potential importance of understanding cell state diversity in the context of therapeutic strategies targeting oncogenic signaling.

Through our quantitative, single-cell resolved analysis of both cell state and signaling, we observed marked differences in baseline signaling as well as in EGF-induced responses between epithelial- and mesenchymal-like cells. Epithelial cells displayed higher basal phosphorylation and robust, transient activation of p-AKT and p-S6. In contrast, mesenchymal cells showed lower baseline activity but sustained activation of p-AKT. These differences were most pronounced for p-AKT and p-S6, whereas p-ERK responses remained relatively conserved across E/M states. This suggests that ERK activation may be decoupled from regulation by E/M status. In support of this, a study in KRAS-mutant NSCLC cells showed that the phosphorylation of AKT, p70S6K (the kinase upstream of p-S6), and PRAS40 (an AKT substrate) was decreased after induction of the EMT, whereas p-ERK levels remained unchanged [76]. The persistence of ERK activity across E/M states in both studies may reflect constitutive KRAS activation in A549 and other NSCLC lines resulting in decoupling of ERK from E/M state control. This is supported by evidence of EGF-induced ERK responsiveness varying with morphological and contextual features in non-transformed systems [9].

A key finding of our study is that E/M state modulates the subcellular compartmentalization and activation kinetics of AKT in a phosphosite-specific manner. Overall, p-AKT-T308 predominantly localized to punctate compartments resembling endosomes, with some ruffle-associated signaling. In contrast, p-AKT-S473 generally showed a broader distribution across the plasma membrane, ruffles, and nucleus. This differential localization likely reflects the fact that AKT is phosphorylated at T308 and S473 by different kinases (PDK1 and mTORC2, respectively), and that these phosphorylation events occur independently and confer distinct substrate preferences and downstream outcomes [77, 78, 68, 79, 80]. When parsed by E/M states, we find that the two phospho-forms of AKT are modulated in an E/M state-dependent manner, potentially reflecting differences in subcellular activation niches or downstream routing mechanisms.

Interestingly, p-AKT-S473 was enriched in nuclear regions in mesenchymal-like cells but not in epithelial cells. Nuclear AKT has historically been interpreted as evidence of translocation following stimulation, although recent findings dispute this view [81]. The functional relevance of nuclear and peri-nuclear-localized AKT2/3 is nonetheless increasingly evident, particularly in relation to control of transcriptional programs driving tumor progression and therapy resistance [20]. In the context of EMT, TGF-*β* stimulation enhances nuclear AKT activity, which potentiates invasive properties by promoting RNF12-mediated degradation of the inhibitory SMAD7, thereby sustaining TGF-*β* signaling [82]. TGF-*β*-induced nuclear AKT also phosphorylates lamin A, which leads to nuclear deformation and genomic instability [83]. Overexpression of nuclear-targeted AKT promotes cancer stem cell-like phenotypes [84]. In line with these examples, our data support a functional role for nuclear-localized AKT in promoting mesenchymal-like phenotypes and behaviours.

Nuclear AKT has also been implicated in therapy resistance. Following DNA damage, p-AKT-S473 accumulates in the nucleus, where it colocalizes with *γ*H2AX and p-ATM-S1981 to facilitate non-homologous end joining repair [85, 86]. This response may be mediated by DNA-PKcs or by PI3K activity acting on nuclear phosphoinositide pools [87]. Our data suggest that mesenchymal-like cells more readily accumulate nuclear p-AKT-S473, raising the possibility that these cells possess a greater capacity for nuclear PI3K/AKT signaling which could confer protection against DNA damage following therapy. Although this was observed in the context of growth factor stimulation rather than genotoxic stress, it remains plausible that mesenchymal-like cells have elevated levels of nuclear-localized PI3K or PDK, or a more efficient mechanism for AKT nuclear translocation. Since AKT lacks a classical nuclear localization signal, its import likely depends on carrier proteins such as TCL1 or RhoB [88, 89]. RhoB is typically downregulated during EMT [90, 91, 92], suggesting that alternative carriers may facilitate nuclear AKT accumulation in mesenchymal cells. Our observation of nuclear p-AKT-S473 enrichment in mesenchymal-like cells is consistent with such context-specific signaling, reinforcing the idea that compartmentalized AKT activity plays diverse roles in cancer progression.

In addition to nuclear localization, we observed kinetic differences including prolonged recruitment of p-AKT-S473 to membrane ruffles in mesenchymal-like cells, in contrast to transient membrane localization in epithelial cells. Aligned with the observed mesenchymal-specific pro-tumorigenic nuclear signaling, sustained p-AKT-S473 recruitment to membrane ruffles has also been linked to cancer cell proliferation, survival, and tumorigenesis [93]. Notably, AKT localization to specific membrane sites (not merely general membrane distribution) is required for the initiation and subsequent formation of invadopodia [94, 95, 20]. These actin-rich protrusive structures mediate extracellular matrix degradation and enable EMT-associated cancer cell invasion [96, 97]. While both epithelial and mesenchymal cells in our study displayed membrane-associated p-AKT-S473, only mesenchymal-like cells exhibited focal enrichment at F-actin-colocalized ruffle sites, consistent with the spatial organization required for invadopodia initiation. This focalization supports a model in which compartmentalized AKT activity underpins a mesenchymal-specific capacity for invasion and metastasis.

Regarding the punctate distribution of p-AKT-T308, we observed that while the total puncta-associated signal was higher in mesenchymal-like cells, the proportion of punctate signal relative to diffuse signal was greater in epithelial cells. Although the precise identity of these puncta remains uncertain, they likely correspond to endosomal and/or lysosomal compartments based on prior reports of such compartment-specific AKT localization and function [20]. At early endosomes, AKT has been shown to phosphorylate and inhibit GSK3*β*, contributing to feedback regulation of clathrin-mediated endocytosis via dynamin-1 [70, 73]. Following growth factor stimulation, lysosome-localized AKT phosphorylates TSC2, relieving its inhibition of mTORC1 to promote local activation of growth and metabolic pathways [98, 99]. These observations suggest that E/M state-dependent differences in p-AKT-T308 localization may shift downstream signaling priorities, ranging from endocytosis regulation to mTORC1-driven nutrient sensing. It is worth noting that compartment-specific AKT functions are largely inferred from colocalization with known interactors, rather than direct local perturbation [20]. As such, their precise roles remain unclear, and future tools enabling spatially resolved functional manipulation will be key to uncovering these local activities [100, 20].

Beyond considering individual phospho-sites, our computational multiplexing approach enabled integrated assessment of multiple phospho-markers across the EGF response landscape. Using manifold learning and pseudotime analysis, we found that epithelial cells progressed rapidly through early phases of signaling, whereas mesenchymal cells dominated later pseudotime intervals. These dynamics suggest that epithelial cells favor acute, robust responses, while mesenchymal cells adopt slower, spatially focused, and sustained signaling. Correlation analysis across signaling features revealed that although overall network structure was preserved between states, ∼18% of pairwise relationships were rewired (either lost or newly formed), highlighting state-specific reconfiguration of signaling subnetworks. This suggests that the EMT may selectively rewire subnetworks within the broader signaling architecture. Comparable findings were reported by Krishnaswamy et al. [101], who used single-cell information-theoretic models to demonstrate dynamic changes in signaling network edges and information flow throughout the EMT in breast cancer cells.

Taken together, our findings have important clinical implications. In tumors, the co-existence of heterogeneous E/M states [102, 103] implies that populations of cancer cells may differentially interpret and respond to oncogenic signals. By extension, this may result in varied sensitivities to therapies targeting these signaling pathways. For example, mesenchymal cells may evade inhibitors targeting membrane-localized signaling by routing AKT activity to nuclear compartments, where signaling is sustained independently of canonical upstream inputs. Such heterogeneity may underlie resistance to PI3K or mTOR inhibitors in EMT-high tumors and supports the rationale for combination strategies that disrupt signaling independent of E/M state [104, 105, 106, 107]. Our study underscores the necessity of modelling and accounting for E/M heterogeneity when developing targeted therapeutic approaches, particularly in cancers with heightened EMT plasticity.

Future studies will be required to assess whether the state-dependent signaling patterns observed here extend to other stimuli relevant to the tumor microenvironment, including cytokines and hormones. Additionally, emerging studies testing the state-dependent effects of targeted pathway inhibitors should now be applied in the context of E/M state and cancer [11]. Such investigations could reveal whether the phospho-dynamics we observed represent generalized regulatory programs shaped by E/M status. The inclusion of EMT models with stronger state contrasts—such as TGF-*β*–induced EMT, sorted epithelial versus mesenchymal populations, or patient-derived systems—would further test the robustness and generalizability of these findings across diverse biological and clinical settings.

Ultimately, we show that naturally occurring E/M cell state heterogeneity in NSCLC cells is a stable predictor of how cells perceive an acute oncogenic growth factor signal, EGF. E/M state heterogeneity results in differential magnitudes, dynamics, spatial organization, and coordination of phospho-signaling responses; all of which can modulate downstream cellular responses. By integrating image-based profiling with computational multiplexing, we thus uncover how cell state shapes both the architecture and progression of multi-molecular signaling responses. These insights emphasize the need to account for phenotypic diversity in therapeutic design and support the use of quantitative imaging approaches to resolve signaling heterogeneity in cancer.

## Data and code availability

Analysis code for all figures will be made available online at: https://github.com/CancerSystemsMicroscopyLab/XXX.

Images and feature tables will be made available at: https://zenodo.org/records/xxxxxxx.

## Acknowledgements

F.V.K. and I.G. are supported by Australian Government Research Training Program (RTP) Scholarships. F.V.K. receives a Top-Up award from SPHERE Cancer CAG and Cancer Institute NSW. J.G.L. is supported by a University of New South Wales Scientia Research Fellowship, a Ramaciotti Biomedical Research Award, an ARC Development Project grant (DP170103599), NHMRC Ideas Grants (GNT1184009, GNT2012848, GNT2028506), and a Tour de Cure Pioneering Grant (RSP-547-FY2023).

## Author contributions

F.V.K. and J.G.L. conceived the study and designed the project. F.V.K. performed all data analyses. C.J. generated the experimental data. T.H., I.G., D.P.N., F.V., F.V.K., and J.G.L. contributed to the early development of the computational multiplexing framework. F.V.K. and J.G.L. led the design and writing of the manuscript, with significant conceptual input from C.L.C.

