## Supplemental Material for "Epithelial–mesenchymal cell state heterogeneity predetermines differential phospho-signaling responses to EGF stimulation" (from another version of this preprint: version 1, 2025-07-09)

### 5 Suppelemental Figures related to manuscript: Epithelial–mesenchymal cell state heterogeneity predetermines differential phospho-signaling responses to EGF stimulation

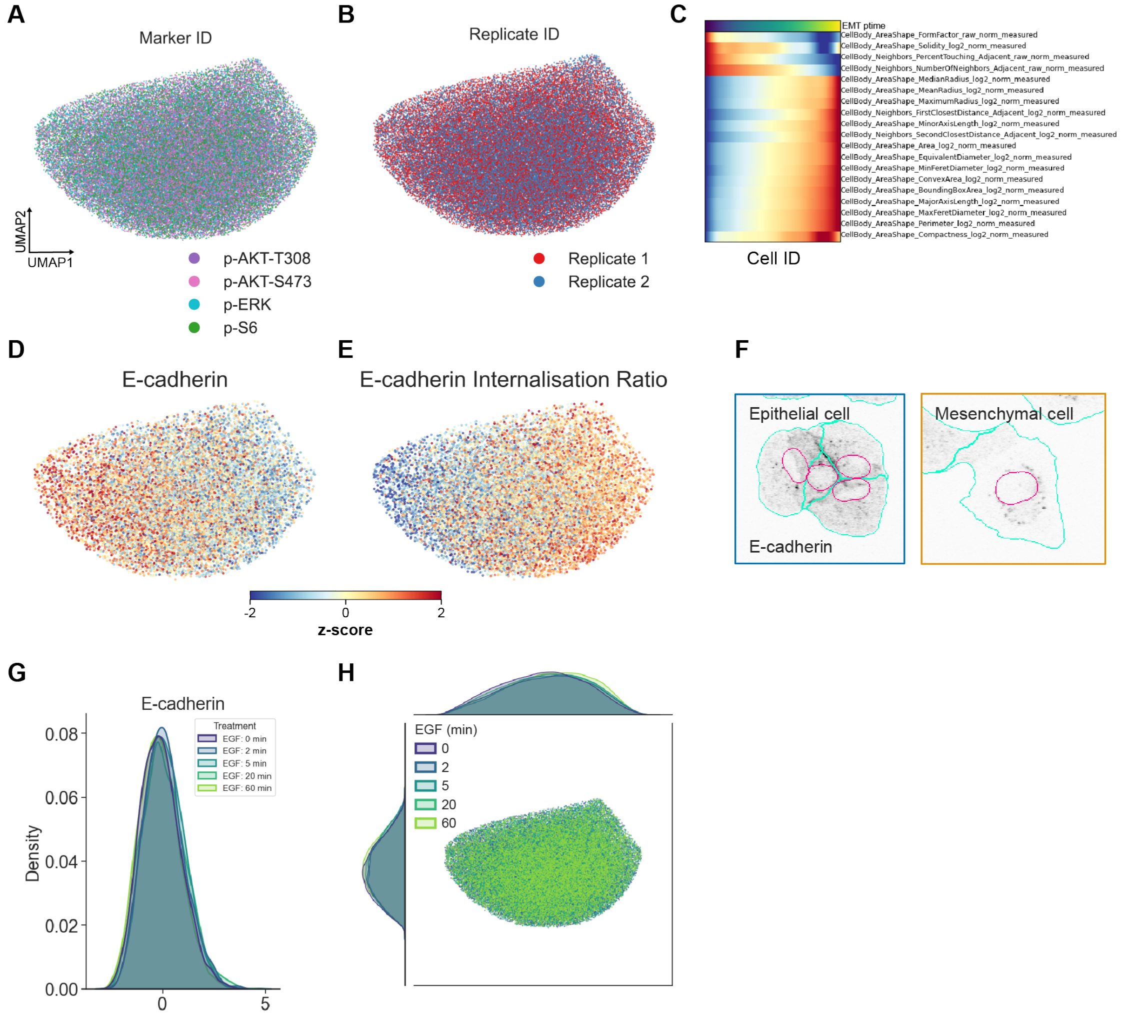


Figure S1: **Construction and validation of the epithelial–mesenchymal (E/M) landscape** UMAP of the E/M landscape colored by marker ID (A) and replicate ID (B) shows thorough mixing and high reproducibility. (C) All E/M state morphological features that significantly vary along EMT pseudotime. Projection of E-cadherin labeled cells into the EMT UMAP, and colored by (D) total levels and (E) internalization ratio (calculated as mean inner cell intensity / mean outer cell intensity; blue implies bias to surface localization, red implies bias to internal localization). (F) Representative cell images of E-cadherin labeled epithelial and mesenchymal cells. EGF stimulation does not alter (G) E-cadherin levels or (H) E/M landscape structure, confirming the stability of this cell state representation across treatments.

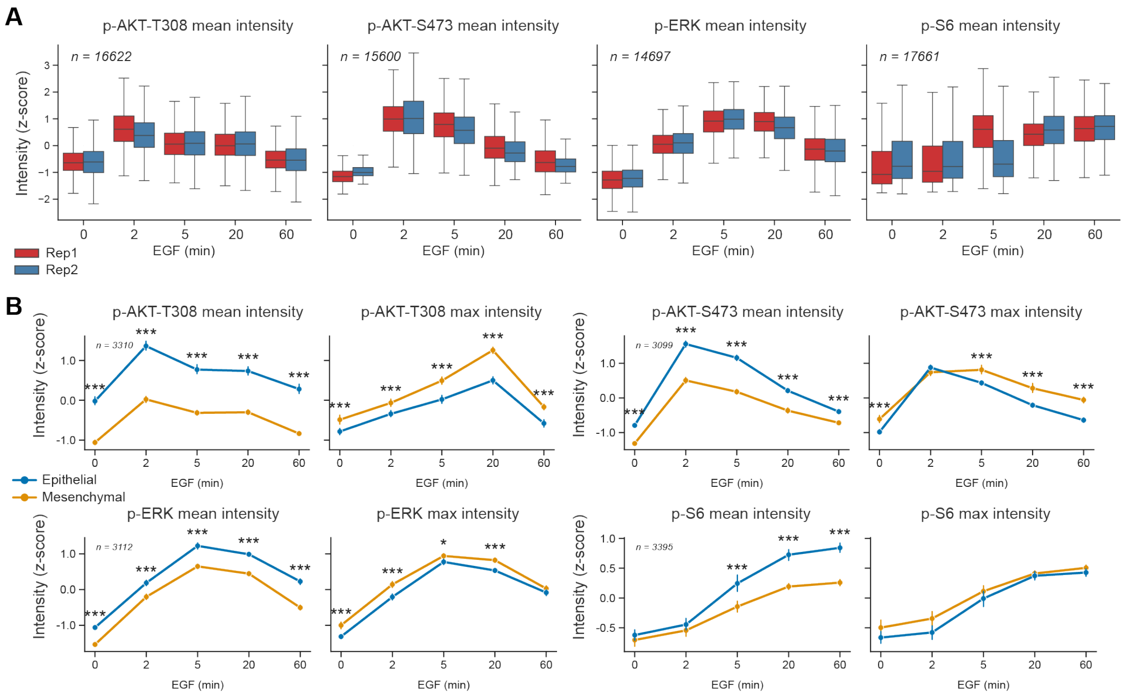


Figure S2: **Reproducibility, spatial dynamics, and raw intensity values for EGF-induced phospho-signal responses** (A) Reproducibility of phospho-signal responses across biological replicates. Boxplots show per-cell mean intensity values for replicate 1 and replicate 2. (B) Raw (non-normalized to control) per-cell intensity values across the EGF time course for all four phosphoproteins. Statistical significance between epithelial and mesenchymal cells was assessed using a two-tailed t-test (***$p < 0.001$, **$p < 0.01$, *$p < 0.05$). Line plots represent the mean $\pm$ 95% confidence interval.

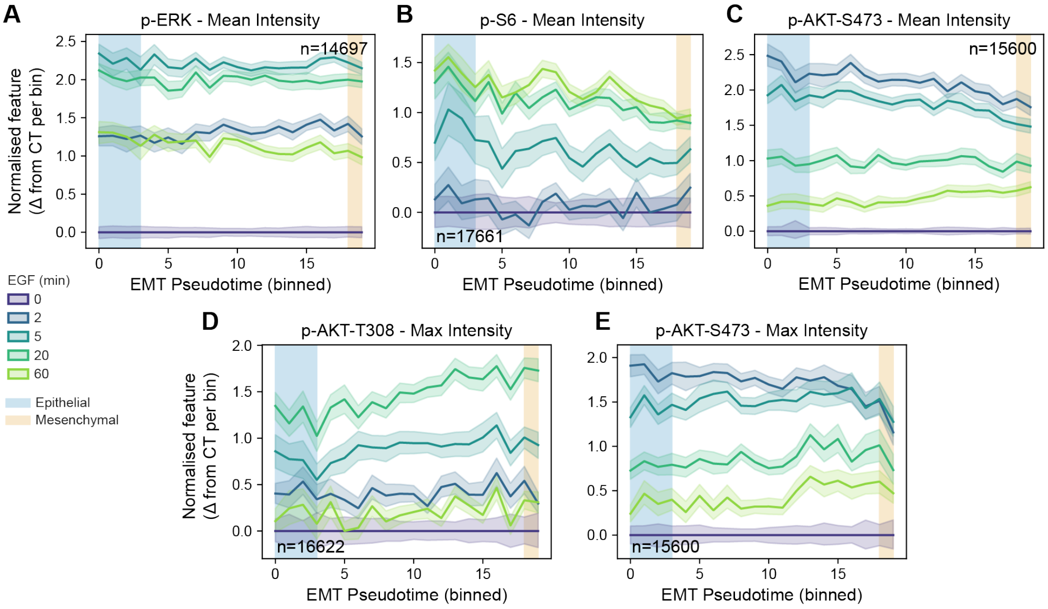


Figure S3: **phospho-signaling responses vary progressively across the EMT continuum** Mean or maximum signal intensity is shown across EMT pseudotime for key phosphoproteins following EGF stimulation. Mean intensity is plotted for (A) p-ERK, (B) p-S6, and (C) p-AKT-S473, while maximum intensity is shown for (D) p-AKT-T308 and (E) p-AKT-S473. Cells were grouped into 20 bins along the EMT pseudotime axis, and within each bin, EGF-stimulated conditions were normalized to the corresponding 0 min control. The resulting feature represents the normalized intensity change from control per bin, allowing direct comparison of EGF-induced signal dynamics across the EMT continuum. Line plots represent mean $\pm$ 95% confidence intervals. Vertical blue and orange shading indicates pseudotime regions corresponding to epithelial- and mesenchymal-like cells, respectively. .

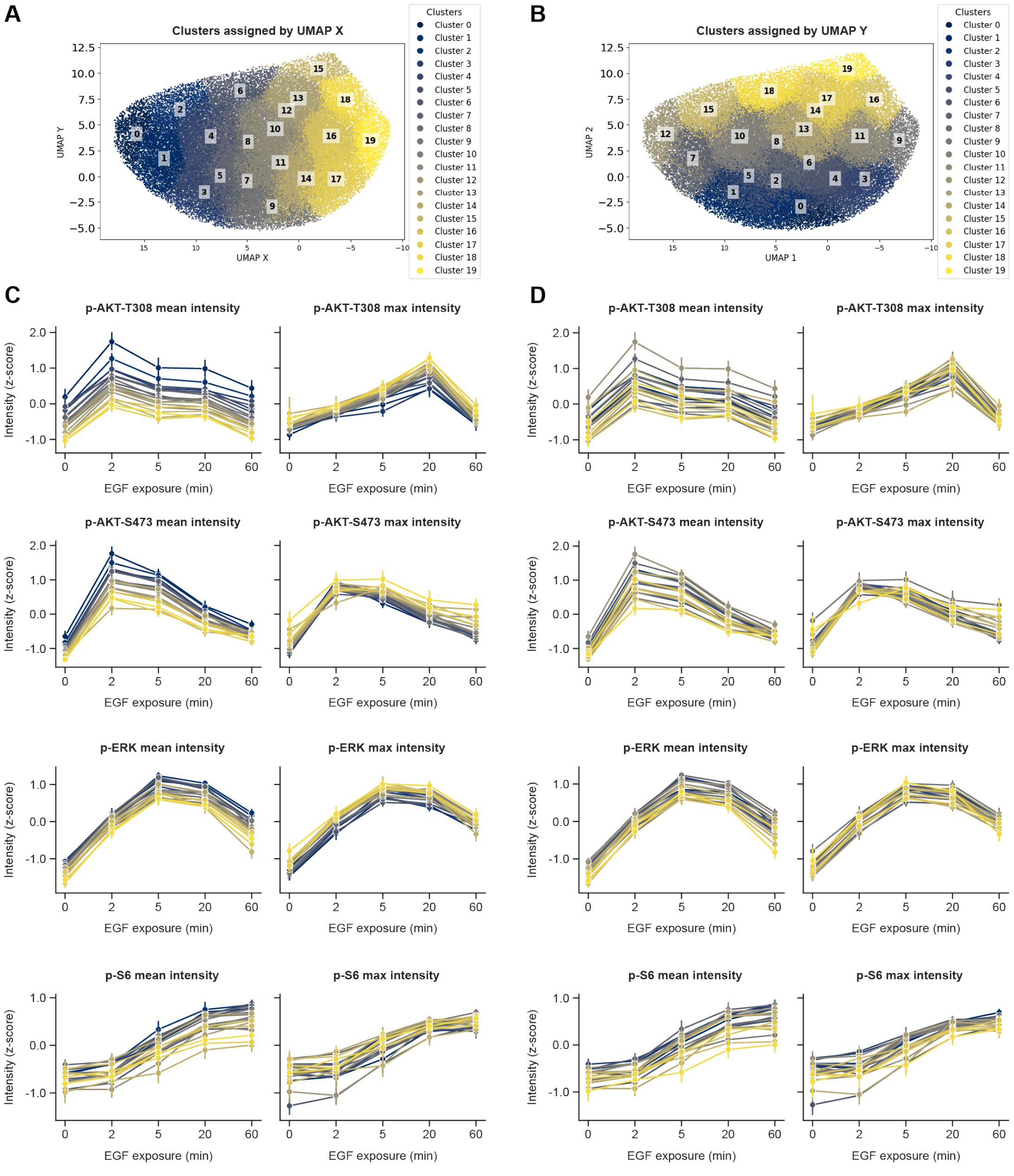


Figure S4: **Cluster-based analysis supports progressive signaling dynamics across the EMT landscape** K-means clustering of E/M morphology features identifies 20 phenotypic clusters spanning the EMT landscape, reordered by mean (A) UMAP X coordinate or (B) UMAP Y coordinate. Mean and maximum intensity EGF time courses grouped by (C) UMAP X cluster or (D) UMAP Y cluster. Line plots show mean $\pm$ 95% confidence intervals.

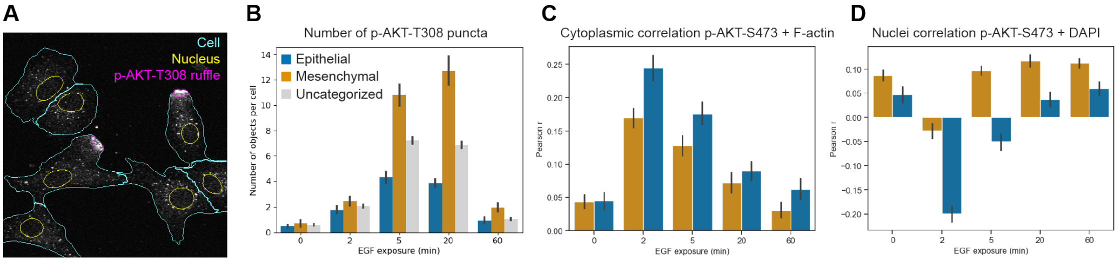


Figure S5: **Additional data for p-AKT subcellular localization** (A) Representative immunofluorescence image showing p-AKT-T308 membrane ruffles. (B) Quantification of p-AKT-T308 puncta per cell across time points and EMT states shows elevated puncta counts in mesenchymal cells at 5–20 min post-EGF. (C) Pixel-level Pearson correlation (Pearson $r$) between p-AKT-S473 and F-actin (phalloidin) in the cytoplasm. (D) Pixel-level Pearson correlation between p-AKT-S473 and DAPI in the nucleus. Data are shown as mean $\pm$ 95% confidence interval across single cells, grouped by EMT classification and EGF treatment time.

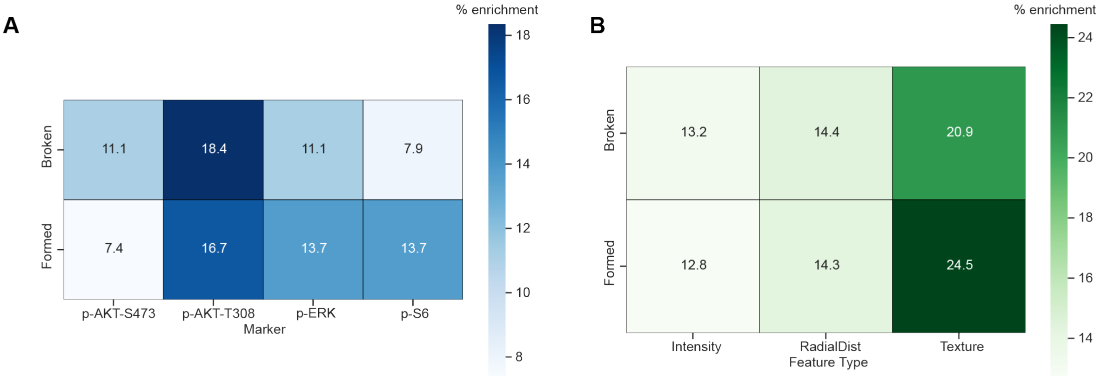


Figure S6: **Marker- and feature-type-specific enrichment of E/M state-dependent signaling relationships** (A) Heatmap showing the percentage of feature–feature pairs involving each signaling marker that exhibited formed (bottom row) or broken (top row) correlations when comparing epithelial and mesenchymal cell states. p-AKT-T308 features were most frequently involved in both formed and broken relationships. (B) Heatmap of the same analysis grouped by feature type. Texture features showed the greatest enrichment for E/M-specific relationships. Values are expressed as percentages of all formed or broken relationships.

Table S1: Features used for Computational Multiplexing. Common features were used as input variables for regression model training, variable features were predicted as outputs.

| **Common features (inputs)** | **Variable features (outputs)** |
|---|---|
| CellBody-AreaShape-Area-log2 | CellBody-Granularity-10-ex555nm-raw |
| CellBody-AreaShape-BoundingBoxArea-log2 | CellBody-Granularity-11-ex555nm-raw |
| CellBody-AreaShape-Compactness-log2 | CellBody-Granularity-12-ex555nm-raw |
| CellBody-AreaShape-ConvexArea-log2 | CellBody-Granularity-13-ex555nm-raw |
| CellBody-AreaShape-Eccentricity-raw | CellBody-Granularity-14-ex555nm-raw |
| CellBody-AreaShape-EquivalentDiameter-log2 | CellBody-Granularity-15-ex555nm-raw |
| CellBody-AreaShape-Extent-raw | CellBody-Granularity-16-ex555nm-raw |
| CellBody-AreaShape-FormFactor-raw | CellBody-Granularity-1-ex555nm-raw |
| CellBody-AreaShape-MajorAxisLength-log2 | CellBody-Granularity-2-ex555nm-raw |
| CellBody-AreaShape-MaxFeretDiameter-log2 | CellBody-Granularity-3-ex555nm-raw |
| CellBody-AreaShape-MaximumRadius-log2 | CellBody-Granularity-4-ex555nm-raw |
| CellBody-AreaShape-MeanRadius-log2 | CellBody-Granularity-5-ex555nm-raw |
| CellBody-AreaShape-MedianRadius-log2 | CellBody-Granularity-6-ex555nm-raw |
| CellBody-AreaShape-MinFeretDiameter-log2 | CellBody-Granularity-7-ex555nm-raw |
| CellBody-AreaShape-MinorAxisLength-log2 | CellBody-Granularity-8-ex555nm-raw |
| CellBody-AreaShape-Perimeter-log2 | CellBody-Granularity-9-ex555nm-raw |
| CellBody-AreaShape-Solidity-log2 | CellBody-Intensity-IntegratedIntensityEdge-ex555nm-log2 |
| CellBody-Granularity-10-ex405nm-raw | CellBody-Intensity-IntegratedIntensity-ex555nm-log2 |
| CellBody-Granularity-10-ex647nm-raw | CellBody-Intensity-LowerQuartileIntensity-ex555nm-log2 |
| CellBody-Granularity-11-ex405nm-raw | CellBody-Intensity-MADIntensity-ex555nm-log2 |
| CellBody-Granularity-11-ex647nm-raw | CellBody-Intensity-MassDisplacement-ex555nm-log2 |
| CellBody-Granularity-12-ex405nm-raw | CellBody-Intensity-MaxIntensityEdge-ex555nm-log2 |
| CellBody-Granularity-12-ex647nm-raw | CellBody-Intensity-MaxIntensity-ex555nm-log2 |

*Continued on next page*

| Common features (inputs) | Variable features (outputs) |
|---|---|
| CellBody-Granularity-13-ex405nm-raw | CellBody-Intensity-MeanIntensityEdge-ex555nm-log2 |
| CellBody-Granularity-13-ex647nm-raw | CellBody-Intensity-MeanIntensity-ex555nm-log2 |
| CellBody-Granularity-14-ex405nm-raw | CellBody-Intensity-MedianIntensity-ex555nm-log2 |
| CellBody-Granularity-14-ex647nm-raw | CellBody-Intensity-MinIntensityEdge-ex555nm-log2 |
| CellBody-Granularity-15-ex405nm-raw | CellBody-Intensity-MinIntensity-ex555nm-log2 |
| CellBody-Granularity-15-ex647nm-raw | CellBody-Intensity-StdIntensityEdge-ex555nm-log2 |
| CellBody-Granularity-16-ex405nm-raw | CellBody-Intensity-StdIntensity-ex555nm-log2 |
| CellBody-Granularity-16-ex647nm-raw | CellBody-Intensity-UpperQuartileIntensity-ex555nm-log2 |
| CellBody-Granularity-1-ex405nm-raw | CellBody-RadialDistribution-FracAtD-ex555nm-1of4-log2 |
| CellBody-Granularity-1-ex647nm-raw | CellBody-RadialDistribution-FracAtD-ex555nm-2of4-log2 |
| CellBody-Granularity-2-ex405nm-raw | CellBody-RadialDistribution-FracAtD-ex555nm-3of4-log2 |
| CellBody-Granularity-2-ex647nm-raw | CellBody-RadialDistribution-FracAtD-ex555nm-4of4-log2 |
| CellBody-Granularity-3-ex405nm-raw | CellBody-RadialDistribution-MeanFrac-ex555nm-1of4-log2 |
| CellBody-Granularity-3-ex647nm-raw | CellBody-RadialDistribution-MeanFrac-ex555nm-2of4-log2 |
| CellBody-Granularity-4-ex405nm-raw | CellBody-RadialDistribution-MeanFrac-ex555nm-3of4-log2 |
| CellBody-Granularity-4-ex647nm-raw | CellBody-RadialDistribution-MeanFrac-ex555nm-4of4-log2 |
| CellBody-Granularity-5-ex405nm-raw | CellBody-RadialDistribution-RadialCV-ex555nm-1of4-log2 |
| CellBody-Granularity-5-ex647nm-raw | CellBody-RadialDistribution-RadialCV-ex555nm-2of4-log2 |

*Continued on next page*

| Common features (inputs) | Variable features (outputs) |
|---|---|
| CellBody-Granularity-6-ex405nm-raw | CellBody-RadialDistribution-RadialCV-ex555nm-3of4-log2 |
| CellBody-Granularity-6-ex647nm-raw | CellBody-RadialDistribution-RadialCV-ex555nm-4of4-log2 |
| CellBody-Granularity-7-ex405nm-raw | CellBody-RadialDistribution-ZernikeMagnitude-ex555nm-0-0-log2 |
| CellBody-Granularity-7-ex647nm-raw | CellBody-RadialDistribution-ZernikeMagnitude-ex555nm-1-1-log2 |
| CellBody-Granularity-8-ex405nm-raw | CellBody-RadialDistribution-ZernikeMagnitude-ex555nm-2-0-log2 |
| CellBody-Granularity-8-ex647nm-raw | CellBody-RadialDistribution-ZernikeMagnitude-ex555nm-2-2-log2 |
| CellBody-Granularity-9-ex405nm-raw | CellBody-RadialDistribution-ZernikeMagnitude-ex555nm-3-1-log2 |
| CellBody-Granularity-9-ex647nm-raw | CellBody-RadialDistribution-ZernikeMagnitude-ex555nm-3-3-log2 |
| CellBody-Intensity-IntegratedIntensityEdge-ex405nm-log2 | CellBody-RadialDistribution-ZernikeMagnitude-ex555nm-4-0-log2 |
| CellBody-Intensity-IntegratedIntensityEdge-ex647nm-log2 | CellBody-RadialDistribution-ZernikeMagnitude-ex555nm-4-2-log2 |
| CellBody-Intensity-IntegratedIntensity-ex405nm-log2 | CellBody-RadialDistribution-ZernikeMagnitude-ex555nm-4-4-log2 |
| CellBody-Intensity-IntegratedIntensity-ex647nm-log2 | CellBody-RadialDistribution-ZernikeMagnitude-ex555nm-5-1-log2 |
| CellBody-Intensity-LowerQuartileIntensity-ex405nm-log2 | CellBody-RadialDistribution-ZernikeMagnitude-ex555nm-5-3-log2 |
| CellBody-Intensity-LowerQuartileIntensity-ex647nm-log2 | CellBody-RadialDistribution-ZernikeMagnitude-ex555nm-5-5-log2 |
| CellBody-Intensity-MADIntensity-ex405nm-log2 | CellBody-RadialDistribution-ZernikeMagnitude-ex555nm-6-0-log2 |
| CellBody-Intensity-MADIntensity-ex647nm-log2 | CellBody-RadialDistribution-ZernikeMagnitude-ex555nm-6-2-log2 |
| CellBody-Intensity-MassDisplacement-ex405nm-log2 | CellBody-RadialDistribution-ZernikeMagnitude-ex555nm-6-4-log2 |

*Continued on next page*

| Common features (inputs) | Variable features (outputs) |
|---|---|
| CellBody-Intensity-MassDisplacement-ex647nm-log2 | CellBody-RadialDistribution-ZernikeMagnitude-ex555nm-6-6-log2 |
| CellBody-Intensity-MaxIntensityEdge-ex405nm-log2 | CellBody-RadialDistribution-ZernikeMagnitude-ex555nm-7-1-log2 |
| CellBody-Intensity-MaxIntensityEdge-ex647nm-log2 | CellBody-RadialDistribution-ZernikeMagnitude-ex555nm-7-3-log2 |
| CellBody-Intensity-MaxIntensity-ex405nm-log2 | CellBody-RadialDistribution-ZernikeMagnitude-ex555nm-7-5-log2 |
| CellBody-Intensity-MaxIntensity-ex647nm-log2 | CellBody-RadialDistribution-ZernikeMagnitude-ex555nm-7-7-log2 |
| CellBody-Intensity-MeanIntensityEdge-ex405nm-log2 | CellBody-RadialDistribution-ZernikeMagnitude-ex555nm-8-0-log2 |
| CellBody-Intensity-MeanIntensityEdge-ex647nm-log2 | CellBody-RadialDistribution-ZernikeMagnitude-ex555nm-8-2-log2 |
| CellBody-Intensity-MeanIntensity-ex405nm-log2 | CellBody-RadialDistribution-ZernikeMagnitude-ex555nm-8-4-log2 |
| CellBody-Intensity-MeanIntensity-ex647nm-log2 | CellBody-RadialDistribution-ZernikeMagnitude-ex555nm-8-6-log2 |
| CellBody-Intensity-MedianIntensity-ex405nm-log2 | CellBody-RadialDistribution-ZernikeMagnitude-ex555nm-8-8-log2 |
| CellBody-Intensity-MedianIntensity-ex647nm-log2 | CellBody-RadialDistribution-ZernikeMagnitude-ex555nm-9-1-log2 |
| CellBody-Intensity-MinIntensityEdge-ex405nm-log2 | CellBody-RadialDistribution-ZernikeMagnitude-ex555nm-9-3-log2 |
| CellBody-Intensity-MinIntensityEdge-ex647nm-log2 | CellBody-RadialDistribution-ZernikeMagnitude-ex555nm-9-5-log2 |
| CellBody-Intensity-MinIntensity-ex405nm-log2 | CellBody-RadialDistribution-ZernikeMagnitude-ex555nm-9-7-log2 |
| CellBody-Intensity-MinIntensity-ex647nm-log2 | CellBody-RadialDistribution-ZernikeMagnitude-ex555nm-9-9-log2 |
| CellBody-Intensity-StdIntensityEdge-ex405nm-log2 | CellBody-Texture-AngularSecondMoment-ex555nm-3-00-256-log2 |
| CellBody-Intensity-StdIntensityEdge-ex647nm-log2 | CellBody-Texture-AngularSecondMoment-ex555nm-3-01-256-log2 |

*Continued on next page*

| Common features (inputs) | Variable features (outputs) |
|---|---|
| CellBody-Intensity-StdIntensity-ex405nm-log2 | CellBody-Texture-AngularSecondMoment-ex555nm-3-02-256-log2 |
| CellBody-Intensity-StdIntensity-ex647nm-log2 | CellBody-Texture-AngularSecondMoment-ex555nm-3-03-256-log2 |
| CellBody-Intensity-UpperQuartileIntensity-ex405nm-log2 | CellBody-Texture-Contrast-ex555nm-3-00-256-sqrt |
| CellBody-Intensity-UpperQuartileIntensity-ex647nm-log2 | CellBody-Texture-Contrast-ex555nm-3-01-256-sqrt |
| CellBody-Neighbors-FirstClosestDistance-Adjacent-log2 | CellBody-Texture-Contrast-ex555nm-3-02-256-sqrt |
| CellBody-Neighbors-NumberOfNeighbors-Adjacent-raw | CellBody-Texture-Contrast-ex555nm-3-03-256-sqrt |
| CellBody-Neighbors-PercentTouching-Adjacent-raw | CellBody-Texture-Correlation-ex555nm-3-00-256-raw |
| CellBody-Neighbors-SecondClosestDistance-Adjacent-log2 | CellBody-Texture-Correlation-ex555nm-3-01-256-raw |
| CellBody-RadialDistribution-FracAtD-ex405nm-1of4-log2 | CellBody-Texture-Correlation-ex555nm-3-02-256-raw |
| CellBody-RadialDistribution-FracAtD-ex405nm-2of4-log2 | CellBody-Texture-Correlation-ex555nm-3-03-256-raw |
| CellBody-RadialDistribution-FracAtD-ex405nm-3of4-log2 | CellBody-Texture-DifferenceEntropy-ex555nm-3-00-256-raw |
| CellBody-RadialDistribution-FracAtD-ex405nm-4of4-log2 | CellBody-Texture-DifferenceEntropy-ex555nm-3-01-256-raw |
| CellBody-RadialDistribution-FracAtD-ex647nm-1of4-log2 | CellBody-Texture-DifferenceEntropy-ex555nm-3-02-256-raw |
| CellBody-RadialDistribution-FracAtD-ex647nm-2of4-log2 | CellBody-Texture-DifferenceEntropy-ex555nm-3-03-256-raw |
| CellBody-RadialDistribution-FracAtD-ex647nm-3of4-log2 | CellBody-Texture-DifferenceVariance-ex555nm-3-00-256-log2 |
| CellBody-RadialDistribution-FracAtD-ex647nm-4of4-log2 | CellBody-Texture-DifferenceVariance-ex555nm-3-01-256-log2 |
| CellBody-RadialDistribution-MeanFrac-ex405nm-1of4-log2 | CellBody-Texture-DifferenceVariance-ex555nm-3-02-256-log2 |

*Continued on next page*

| Common features (inputs) | Variable features (outputs) |
|---|---|
| CellBody-RadialDistribution-MeanFrac-ex405nm-2of4-log2 | CellBody-Texture-DifferenceVariance-ex555nm-3-03-256-log2 |
| CellBody-RadialDistribution-MeanFrac-ex405nm-3of4-log2 | CellBody-Texture-Entropy-ex555nm-3-00-256-raw |
| CellBody-RadialDistribution-MeanFrac-ex405nm-4of4-log2 | CellBody-Texture-Entropy-ex555nm-3-01-256-raw |
| CellBody-RadialDistribution-MeanFrac-ex647nm-1of4-log2 | CellBody-Texture-Entropy-ex555nm-3-02-256-raw |
| CellBody-RadialDistribution-MeanFrac-ex647nm-2of4-log2 | CellBody-Texture-Entropy-ex555nm-3-03-256-raw |
| CellBody-RadialDistribution-MeanFrac-ex647nm-3of4-log2 | CellBody-Texture-InfoMeas1-ex555nm-3-00-256-raw |
| CellBody-RadialDistribution-MeanFrac-ex647nm-4of4-log2 | CellBody-Texture-InfoMeas1-ex555nm-3-01-256-raw |
| CellBody-RadialDistribution-RadialCV-ex405nm-1of4-log2 | CellBody-Texture-InfoMeas1-ex555nm-3-02-256-raw |
| CellBody-RadialDistribution-RadialCV-ex405nm-2of4-log2 | CellBody-Texture-InfoMeas1-ex555nm-3-03-256-raw |
| CellBody-RadialDistribution-RadialCV-ex405nm-3of4-log2 | CellBody-Texture-InfoMeas2-ex555nm-3-00-256-raw |
| CellBody-RadialDistribution-RadialCV-ex405nm-4of4-log2 | CellBody-Texture-InfoMeas2-ex555nm-3-01-256-raw |
| CellBody-RadialDistribution-RadialCV-ex647nm-1of4-log2 | CellBody-Texture-InfoMeas2-ex555nm-3-02-256-raw |
| CellBody-RadialDistribution-RadialCV-ex647nm-2of4-log2 | CellBody-Texture-InfoMeas2-ex555nm-3-03-256-raw |
| CellBody-RadialDistribution-RadialCV-ex647nm-3of4-log2 | CellBody-Texture-InverseDifferenceMoment-ex555nm-3-00-256-raw |
| CellBody-RadialDistribution-RadialCV-ex647nm-4of4-log2 | CellBody-Texture-InverseDifferenceMoment-ex555nm-3-01-256-raw |
| CellBody-RadialDistribution-ZernikeMagnitude-ex405nm-0-0-log2 | CellBody-Texture-InverseDifferenceMoment-ex555nm-3-02-256-raw |
| CellBody-RadialDistribution-ZernikeMagnitude-ex405nm-1-1-log2 | CellBody-Texture-InverseDifferenceMoment-ex555nm-3-03-256-raw |

*Continued on next page*

| Common features (inputs) | Variable features (outputs) |
|---|---|
| CellBody-RadialDistribution-ZernikeMagnitude-ex405nm-2-0-log2 | CellBody-Texture-SumAverage-ex555nm-3-00-256-log2 |
| CellBody-RadialDistribution-ZernikeMagnitude-ex405nm-2-2-log2 | CellBody-Texture-SumAverage-ex555nm-3-01-256-log2 |
| CellBody-RadialDistribution-ZernikeMagnitude-ex405nm-3-1-log2 | CellBody-Texture-SumAverage-ex555nm-3-02-256-log2 |
| CellBody-RadialDistribution-ZernikeMagnitude-ex405nm-3-3-log2 | CellBody-Texture-SumAverage-ex555nm-3-03-256-log2 |
| CellBody-RadialDistribution-ZernikeMagnitude-ex405nm-4-0-log2 | CellBody-Texture-SumEntropy-ex555nm-3-00-256-raw |
| CellBody-RadialDistribution-ZernikeMagnitude-ex405nm-4-2-log2 | CellBody-Texture-SumEntropy-ex555nm-3-01-256-raw |
| CellBody-RadialDistribution-ZernikeMagnitude-ex405nm-4-4-log2 | CellBody-Texture-SumEntropy-ex555nm-3-02-256-raw |
| CellBody-RadialDistribution-ZernikeMagnitude-ex405nm-5-1-log2 | CellBody-Texture-SumEntropy-ex555nm-3-03-256-raw |
| CellBody-RadialDistribution-ZernikeMagnitude-ex405nm-5-3-log2 | CellBody-Texture-SumVariance-ex555nm-3-00-256-sqrt |
| CellBody-RadialDistribution-ZernikeMagnitude-ex405nm-5-5-log2 | CellBody-Texture-SumVariance-ex555nm-3-01-256-sqrt |
| CellBody-RadialDistribution-ZernikeMagnitude-ex405nm-6-0-log2 | CellBody-Texture-SumVariance-ex555nm-3-02-256-sqrt |
| CellBody-RadialDistribution-ZernikeMagnitude-ex405nm-6-2-log2 | CellBody-Texture-SumVariance-ex555nm-3-03-256-sqrt |
| CellBody-RadialDistribution-ZernikeMagnitude-ex405nm-6-4-log2 | CellBody-Texture-Variance-ex555nm-3-00-256-sqrt |
| CellBody-RadialDistribution-ZernikeMagnitude-ex405nm-6-6-log2 | CellBody-Texture-Variance-ex555nm-3-01-256-sqrt |
| CellBody-RadialDistribution-ZernikeMagnitude-ex405nm-7-1-log2 | CellBody-Texture-Variance-ex555nm-3-02-256-sqrt |
| CellBody-RadialDistribution-ZernikeMagnitude-ex405nm-7-3-log2 | CellBody-Texture-Variance-ex555nm-3-03-256-sqrt |
| CellBody-RadialDistribution-ZernikeMagnitude-ex405nm-7-5-log2 | Cytoplasm-Granularity-10-ex555nm-raw |

*Continued on next page*

| Common features (inputs) | Variable features (outputs) |
|---|---|
| CellBody-RadialDistribution-ZernikeMagnitude-ex405nm-7-7-log2 | Cytoplasm-Granularity-11-ex555nm-raw |
| CellBody-RadialDistribution-ZernikeMagnitude-ex405nm-8-0-log2 | Cytoplasm-Granularity-12-ex555nm-raw |
| CellBody-RadialDistribution-ZernikeMagnitude-ex405nm-8-2-log2 | Cytoplasm-Granularity-13-ex555nm-raw |
| CellBody-RadialDistribution-ZernikeMagnitude-ex405nm-8-4-log2 | Cytoplasm-Granularity-14-ex555nm-raw |
| CellBody-RadialDistribution-ZernikeMagnitude-ex405nm-8-6-log2 | Cytoplasm-Granularity-15-ex555nm-raw |
| CellBody-RadialDistribution-ZernikeMagnitude-ex405nm-8-8-log2 | Cytoplasm-Granularity-16-ex555nm-raw |
| CellBody-RadialDistribution-ZernikeMagnitude-ex405nm-9-1-log2 | Cytoplasm-Granularity-1-ex555nm-raw |
| CellBody-RadialDistribution-ZernikeMagnitude-ex405nm-9-3-log2 | Cytoplasm-Granularity-2-ex555nm-raw |
| CellBody-RadialDistribution-ZernikeMagnitude-ex405nm-9-5-log2 | Cytoplasm-Granularity-3-ex555nm-raw |
| CellBody-RadialDistribution-ZernikeMagnitude-ex405nm-9-7-log2 | Cytoplasm-Granularity-4-ex555nm-raw |
| CellBody-RadialDistribution-ZernikeMagnitude-ex405nm-9-9-log2 | Cytoplasm-Granularity-5-ex555nm-raw |
| CellBody-RadialDistribution-ZernikeMagnitude-ex647nm-0-0-log2 | Cytoplasm-Granularity-6-ex555nm-raw |
| CellBody-RadialDistribution-ZernikeMagnitude-ex647nm-1-1-log2 | Cytoplasm-Granularity-7-ex555nm-raw |
| CellBody-RadialDistribution-ZernikeMagnitude-ex647nm-2-0-log2 | Cytoplasm-Granularity-8-ex555nm-raw |
| CellBody-RadialDistribution-ZernikeMagnitude-ex647nm-2-2-log2 | Cytoplasm-Granularity-9-ex555nm-raw |
| CellBody-RadialDistribution-ZernikeMagnitude-ex647nm-3-1-log2 | Cytoplasm-Intensity-IntegratedIntensityEdge-ex555nm-log2 |
| CellBody-RadialDistribution-ZernikeMagnitude-ex647nm-3-3-log2 | Cytoplasm-Intensity-IntegratedIntensity-ex555nm-log2 |

*Continued on next page*

| Common features (inputs) | Variable features (outputs) |
|---|---|
| CellBody-RadialDistribution-ZernikeMagnitude-ex647nm-4-0-log2 | Cytoplasm-Intensity-LowerQuartileIntensity-ex555nm-log2 |
| CellBody-RadialDistribution-ZernikeMagnitude-ex647nm-4-2-log2 | Cytoplasm-Intensity-MADIntensity-ex555nm-log2 |
| CellBody-RadialDistribution-ZernikeMagnitude-ex647nm-4-4-log2 | Cytoplasm-Intensity-MassDisplacement-ex555nm-log2 |
| CellBody-RadialDistribution-ZernikeMagnitude-ex647nm-5-1-log2 | Cytoplasm-Intensity-MaxIntensityEdge-ex555nm-log2 |
| CellBody-RadialDistribution-ZernikeMagnitude-ex647nm-5-3-log2 | Cytoplasm-Intensity-MaxIntensity-ex555nm-log2 |
| CellBody-RadialDistribution-ZernikeMagnitude-ex647nm-5-5-log2 | Cytoplasm-Intensity-MeanIntensityEdge-ex555nm-log2 |
| CellBody-RadialDistribution-ZernikeMagnitude-ex647nm-6-0-log2 | Cytoplasm-Intensity-MeanIntensity-ex555nm-log2 |
| CellBody-RadialDistribution-ZernikeMagnitude-ex647nm-6-2-log2 | Cytoplasm-Intensity-MedianIntensity-ex555nm-log2 |
| CellBody-RadialDistribution-ZernikeMagnitude-ex647nm-6-4-log2 | Cytoplasm-Intensity-MinIntensityEdge-ex555nm-log2 |
| CellBody-RadialDistribution-ZernikeMagnitude-ex647nm-6-6-log2 | Cytoplasm-Intensity-MinIntensity-ex555nm-log2 |
| CellBody-RadialDistribution-ZernikeMagnitude-ex647nm-7-1-log2 | Cytoplasm-Intensity-StdIntensityEdge-ex555nm-log2 |
| CellBody-RadialDistribution-ZernikeMagnitude-ex647nm-7-3-log2 | Cytoplasm-Intensity-StdIntensity-ex555nm-log2 |
| CellBody-RadialDistribution-ZernikeMagnitude-ex647nm-7-5-log2 | Cytoplasm-Intensity-UpperQuartileIntensity-ex555nm-log2 |
| CellBody-RadialDistribution-ZernikeMagnitude-ex647nm-7-7-log2 | Nuclei-Granularity-10-ex555nm-raw |
| CellBody-RadialDistribution-ZernikeMagnitude-ex647nm-8-0-log2 | Nuclei-Granularity-11-ex555nm-raw |
| CellBody-RadialDistribution-ZernikeMagnitude-ex647nm-8-2-log2 | Nuclei-Granularity-12-ex555nm-raw |
| CellBody-RadialDistribution-ZernikeMagnitude-ex647nm-8-4-log2 | Nuclei-Granularity-13-ex555nm-raw |

*Continued on next page*

| **Common features (inputs)** | **Variable features (outputs)** |
|---|---|
| CellBody-RadialDistribution-ZernikeMagnitude-ex647nm-8-6-log2 | Nuclei-Granularity-14-ex555nm-raw |
| CellBody-RadialDistribution-ZernikeMagnitude-ex647nm-8-8-log2 | Nuclei-Granularity-15-ex555nm-raw |
| CellBody-RadialDistribution-ZernikeMagnitude-ex647nm-9-1-log2 | Nuclei-Granularity-16-ex555nm-raw |
| CellBody-RadialDistribution-ZernikeMagnitude-ex647nm-9-3-log2 | Nuclei-Granularity-1-ex555nm-raw |
| CellBody-RadialDistribution-ZernikeMagnitude-ex647nm-9-5-log2 | Nuclei-Granularity-2-ex555nm-raw |
| CellBody-RadialDistribution-ZernikeMagnitude-ex647nm-9-7-log2 | Nuclei-Granularity-3-ex555nm-raw |
| CellBody-RadialDistribution-ZernikeMagnitude-ex647nm-9-9-log2 | Nuclei-Granularity-4-ex555nm-raw |
| CellBody-Texture-AngularSecondMoment-ex405nm-3-00-256-log2 | Nuclei-Granularity-5-ex555nm-raw |
| CellBody-Texture-AngularSecondMoment-ex405nm-3-01-256-log2 | Nuclei-Granularity-6-ex555nm-raw |
| CellBody-Texture-AngularSecondMoment-ex405nm-3-02-256-log2 | Nuclei-Granularity-7-ex555nm-raw |
| CellBody-Texture-AngularSecondMoment-ex405nm-3-03-256-log2 | Nuclei-Granularity-8-ex555nm-raw |
| CellBody-Texture-AngularSecondMoment-ex647nm-3-00-256-log2 | Nuclei-Granularity-9-ex555nm-raw |
| CellBody-Texture-AngularSecondMoment-ex647nm-3-01-256-log2 | Nuclei-Intensity-IntegratedIntensityEdge-ex555nm-log2 |
| CellBody-Texture-AngularSecondMoment-ex647nm-3-02-256-log2 | Nuclei-Intensity-IntegratedIntensity-ex555nm-log2 |
| CellBody-Texture-AngularSecondMoment-ex647nm-3-03-256-log2 | Nuclei-Intensity-LowerQuartileIntensity-ex555nm-log2 |
| CellBody-Texture-Contrast-ex405nm-3-00-256-sqrt | Nuclei-Intensity-MADIntensity-ex555nm-log2 |
| CellBody-Texture-Contrast-ex405nm-3-01-256-sqrt | Nuclei-Intensity-MassDisplacement-ex555nm-log2 |

*Continued on next page*

| Common features (inputs) | Variable features (outputs) |
|---|---|
| CellBody-Texture-Contrast-ex405nm-3-02-256-sqrt | Nuclei-Intensity-MaxIntensityEdge-ex555nm-log2 |
| CellBody-Texture-Contrast-ex405nm-3-03-256-sqrt | Nuclei-Intensity-MaxIntensity-ex555nm-log2 |
| CellBody-Texture-Contrast-ex647nm-3-00-256-sqrt | Nuclei-Intensity-MeanIntensityEdge-ex555nm-log2 |
| CellBody-Texture-Contrast-ex647nm-3-01-256-sqrt | Nuclei-Intensity-MeanIntensity-ex555nm-log2 |
| CellBody-Texture-Contrast-ex647nm-3-02-256-sqrt | Nuclei-Intensity-MedianIntensity-ex555nm-log2 |
| CellBody-Texture-Contrast-ex647nm-3-03-256-sqrt | Nuclei-Intensity-MinIntensityEdge-ex555nm-log2 |
| CellBody-Texture-Correlation-ex405nm-3-00-256-raw | Nuclei-Intensity-MinIntensity-ex555nm-log2 |
| CellBody-Texture-Correlation-ex405nm-3-01-256-raw | Nuclei-Intensity-StdIntensityEdge-ex555nm-log2 |
| CellBody-Texture-Correlation-ex405nm-3-02-256-raw | Nuclei-Intensity-StdIntensity-ex555nm-log2 |
| CellBody-Texture-Correlation-ex405nm-3-03-256-raw | Nuclei-Intensity-UpperQuartileIntensity-ex555nm-log2 |
| CellBody-Texture-Correlation-ex647nm-3-00-256-raw | Nuclei-Texture-AngularSecondMoment-ex555nm-3-00-256-log2 |
| CellBody-Texture-Correlation-ex647nm-3-01-256-raw | Nuclei-Texture-AngularSecondMoment-ex555nm-3-01-256-log2 |
| CellBody-Texture-Correlation-ex647nm-3-02-256-raw | Nuclei-Texture-AngularSecondMoment-ex555nm-3-02-256-log2 |
| CellBody-Texture-Correlation-ex647nm-3-03-256-raw | Nuclei-Texture-AngularSecondMoment-ex555nm-3-03-256-log2 |
| CellBody-Texture-DifferenceEntropy-ex405nm-3-00-256-raw | Nuclei-Texture-Contrast-ex555nm-3-00-256-sqrt |
| CellBody-Texture-DifferenceEntropy-ex405nm-3-01-256-raw | Nuclei-Texture-Contrast-ex555nm-3-01-256-sqrt |
| CellBody-Texture-DifferenceEntropy-ex405nm-3-02-256-raw | Nuclei-Texture-Contrast-ex555nm-3-02-256-sqrt |

*Continued on next page*

| Common features (inputs) | Variable features (outputs) |
|---|---|
| CellBody-Texture-DifferenceEntropy-ex405nm-3-03-256-raw | Nuclei-Texture-Contrast-ex555nm-3-03-256-sqrt |
| CellBody-Texture-DifferenceEntropy-ex647nm-3-00-256-raw | Nuclei-Texture-Correlation-ex555nm-3-00-256-raw |
| CellBody-Texture-DifferenceEntropy-ex647nm-3-01-256-raw | Nuclei-Texture-Correlation-ex555nm-3-01-256-raw |
| CellBody-Texture-DifferenceEntropy-ex647nm-3-02-256-raw | Nuclei-Texture-Correlation-ex555nm-3-02-256-raw |
| CellBody-Texture-DifferenceEntropy-ex647nm-3-03-256-raw | Nuclei-Texture-Correlation-ex555nm-3-03-256-raw |
| CellBody-Texture-DifferenceVariance-ex405nm-3-00-256-log2 | Nuclei-Texture-DifferenceEntropy-ex555nm-3-00-256-raw |
| CellBody-Texture-DifferenceVariance-ex405nm-3-01-256-log2 | Nuclei-Texture-DifferenceEntropy-ex555nm-3-01-256-raw |
| CellBody-Texture-DifferenceVariance-ex405nm-3-02-256-log2 | Nuclei-Texture-DifferenceEntropy-ex555nm-3-02-256-raw |
| CellBody-Texture-DifferenceVariance-ex405nm-3-03-256-log2 | Nuclei-Texture-DifferenceEntropy-ex555nm-3-03-256-raw |
| CellBody-Texture-DifferenceVariance-ex647nm-3-00-256-log2 | Nuclei-Texture-DifferenceVariance-ex555nm-3-00-256-log2 |
| CellBody-Texture-DifferenceVariance-ex647nm-3-01-256-log2 | Nuclei-Texture-DifferenceVariance-ex555nm-3-01-256-log2 |
| CellBody-Texture-DifferenceVariance-ex647nm-3-02-256-log2 | Nuclei-Texture-DifferenceVariance-ex555nm-3-02-256-log2 |
| CellBody-Texture-DifferenceVariance-ex647nm-3-03-256-log2 | Nuclei-Texture-DifferenceVariance-ex555nm-3-03-256-log2 |
| CellBody-Texture-Entropy-ex405nm-3-00-256-raw | Nuclei-Texture-Entropy-ex555nm-3-00-256-raw |
| CellBody-Texture-Entropy-ex405nm-3-01-256-raw | Nuclei-Texture-Entropy-ex555nm-3-01-256-raw |
| CellBody-Texture-Entropy-ex405nm-3-02-256-raw | Nuclei-Texture-Entropy-ex555nm-3-02-256-raw |
| CellBody-Texture-Entropy-ex405nm-3-03-256-raw | Nuclei-Texture-Entropy-ex555nm-3-03-256-raw |

*Continued on next page*

| Common features (inputs) | Variable features (outputs) |
|---|---|
| CellBody-Texture-Entropy-ex647nm-3-00-256-raw | Nuclei-Texture-InfoMeas1-ex555nm-3-00-256-raw |
| CellBody-Texture-Entropy-ex647nm-3-01-256-raw | Nuclei-Texture-InfoMeas1-ex555nm-3-01-256-raw |
| CellBody-Texture-Entropy-ex647nm-3-02-256-raw | Nuclei-Texture-InfoMeas1-ex555nm-3-02-256-raw |
| CellBody-Texture-Entropy-ex647nm-3-03-256-raw | Nuclei-Texture-InfoMeas1-ex555nm-3-03-256-raw |
| CellBody-Texture-InfoMeas1-ex405nm-3-00-256-raw | Nuclei-Texture-InfoMeas2-ex555nm-3-00-256-raw |
| CellBody-Texture-InfoMeas1-ex405nm-3-01-256-raw | Nuclei-Texture-InfoMeas2-ex555nm-3-01-256-raw |
| CellBody-Texture-InfoMeas1-ex405nm-3-02-256-raw | Nuclei-Texture-InfoMeas2-ex555nm-3-02-256-raw |
| CellBody-Texture-InfoMeas1-ex405nm-3-03-256-raw | Nuclei-Texture-InfoMeas2-ex555nm-3-03-256-raw |
| CellBody-Texture-InfoMeas1-ex647nm-3-00-256-raw | Nuclei-Texture-InverseDifferenceMoment-ex555nm-3-00-256-raw |
| CellBody-Texture-InfoMeas1-ex647nm-3-01-256-raw | Nuclei-Texture-InverseDifferenceMoment-ex555nm-3-01-256-raw |
| CellBody-Texture-InfoMeas1-ex647nm-3-02-256-raw | Nuclei-Texture-InverseDifferenceMoment-ex555nm-3-02-256-raw |
| CellBody-Texture-InfoMeas1-ex647nm-3-03-256-raw | Nuclei-Texture-InverseDifferenceMoment-ex555nm-3-03-256-raw |
| CellBody-Texture-InfoMeas2-ex405nm-3-00-256-raw | Nuclei-Texture-SumAverage-ex555nm-3-00-256-log2 |
| CellBody-Texture-InfoMeas2-ex405nm-3-01-256-raw | Nuclei-Texture-SumAverage-ex555nm-3-01-256-log2 |
| CellBody-Texture-InfoMeas2-ex405nm-3-02-256-raw | Nuclei-Texture-SumAverage-ex555nm-3-02-256-log2 |
| CellBody-Texture-InfoMeas2-ex405nm-3-03-256-raw | Nuclei-Texture-SumAverage-ex555nm-3-03-256-log2 |
| CellBody-Texture-InfoMeas2-ex647nm-3-00-256-raw | Nuclei-Texture-SumEntropy-ex555nm-3-00-256-raw |

*Continued on next page*

| Common features (inputs) | Variable features (outputs) |
|---|---|
| CellBody-Texture-InfoMeas2-ex647nm-3-01-256-raw | Nuclei-Texture-SumEntropy-ex555nm-3-01-256-raw |
| CellBody-Texture-InfoMeas2-ex647nm-3-02-256-raw | Nuclei-Texture-SumEntropy-ex555nm-3-02-256-raw |
| CellBody-Texture-InfoMeas2-ex647nm-3-03-256-raw | Nuclei-Texture-SumEntropy-ex555nm-3-03-256-raw |
| CellBody-Texture-InverseDifferenceMoment-ex405nm-3-00-256-raw | Nuclei-Texture-SumVariance-ex555nm-3-00-256-sqrt |
| CellBody-Texture-InverseDifferenceMoment-ex405nm-3-01-256-raw | Nuclei-Texture-SumVariance-ex555nm-3-01-256-sqrt |
| CellBody-Texture-InverseDifferenceMoment-ex405nm-3-02-256-raw | Nuclei-Texture-SumVariance-ex555nm-3-02-256-sqrt |
| CellBody-Texture-InverseDifferenceMoment-ex405nm-3-03-256-raw | Nuclei-Texture-SumVariance-ex555nm-3-03-256-sqrt |
| CellBody-Texture-InverseDifferenceMoment-ex647nm-3-00-256-raw | Nuclei-Texture-Variance-ex555nm-3-00-256-sqrt |
| CellBody-Texture-InverseDifferenceMoment-ex647nm-3-01-256-raw | Nuclei-Texture-Variance-ex555nm-3-01-256-sqrt |
| CellBody-Texture-InverseDifferenceMoment-ex647nm-3-02-256-raw | Nuclei-Texture-Variance-ex555nm-3-02-256-sqrt |
| CellBody-Texture-InverseDifferenceMoment-ex647nm-3-03-256-raw | Nuclei-Texture-Variance-ex555nm-3-03-256-sqrt |
| CellBody-Texture-SumAverage-ex405nm-3-00-256-log2 | Ratio-NC-ex555nm-IntegInt-log2 |
| CellBody-Texture-SumAverage-ex405nm-3-01-256-log2 | Ratio-NC-ex555nm-MeanInt-log2 |
| CellBody-Texture-SumAverage-ex405nm-3-02-256-log2 | - |
| CellBody-Texture-SumAverage-ex405nm-3-03-256-log2 | - |
| CellBody-Texture-SumAverage-ex647nm-3-00-256-log2 | - |
| CellBody-Texture-SumAverage-ex647nm-3-01-256-log2 | - |

*Continued on next page*

| Common features (inputs) | Variable features (outputs) |
|---|---|
| CellBody-Texture-SumAverage-ex647nm-3-02-256-log2 | - |
| CellBody-Texture-SumAverage-ex647nm-3-03-256-log2 | - |
| CellBody-Texture-SumEntropy-ex405nm-3-00-256-raw | - |
| CellBody-Texture-SumEntropy-ex405nm-3-01-256-raw | - |
| CellBody-Texture-SumEntropy-ex405nm-3-02-256-raw | - |
| CellBody-Texture-SumEntropy-ex405nm-3-03-256-raw | - |
| CellBody-Texture-SumEntropy-ex647nm-3-00-256-raw | - |
| CellBody-Texture-SumEntropy-ex647nm-3-01-256-raw | - |
| CellBody-Texture-SumEntropy-ex647nm-3-02-256-raw | - |
| CellBody-Texture-SumEntropy-ex647nm-3-03-256-raw | - |
| CellBody-Texture-SumVariance-ex405nm-3-00-256-sqrt | - |
| CellBody-Texture-SumVariance-ex405nm-3-01-256-sqrt | - |
| CellBody-Texture-SumVariance-ex405nm-3-02-256-sqrt | - |
| CellBody-Texture-SumVariance-ex405nm-3-03-256-sqrt | - |
| CellBody-Texture-SumVariance-ex647nm-3-00-256-sqrt | - |
| CellBody-Texture-SumVariance-ex647nm-3-01-256-sqrt | - |
| CellBody-Texture-SumVariance-ex647nm-3-02-256-sqrt | - |

*Continued on next page*

| Common features (inputs) | Variable features (outputs) |
|---|---|
| CellBody-Texture-SumVariance-ex647nm-3-03-256-sqrt | - |
| CellBody-Texture-Variance-ex405nm-3-00-256-sqrt | - |
| CellBody-Texture-Variance-ex405nm-3-01-256-sqrt | - |
| CellBody-Texture-Variance-ex405nm-3-02-256-sqrt | - |
| CellBody-Texture-Variance-ex405nm-3-03-256-sqrt | - |
| CellBody-Texture-Variance-ex647nm-3-00-256-sqrt | - |
| CellBody-Texture-Variance-ex647nm-3-01-256-sqrt | - |
| CellBody-Texture-Variance-ex647nm-3-02-256-sqrt | - |
| CellBody-Texture-Variance-ex647nm-3-03-256-sqrt | - |
| Cytoplasm-AreaShape-Area-log2 | - |
| Cytoplasm-AreaShape-BoundingBoxArea-log2 | - |
| Cytoplasm-AreaShape-Compactness-log2 | - |
| Cytoplasm-AreaShape-ConvexArea-log2 | - |
| Cytoplasm-AreaShape-Eccentricity-raw | - |
| Cytoplasm-AreaShape-EquivalentDiameter-log2 | - |
| Cytoplasm-AreaShape-Extent-raw | - |
| Cytoplasm-AreaShape-FormFactor-raw | - |
| Cytoplasm-AreaShape-MajorAxisLength-log2 | - |
| Cytoplasm-AreaShape-MaxFeretDiameter-log2 | - |
| Cytoplasm-AreaShape-MaximumRadius-log2 | - |
| Cytoplasm-AreaShape-MeanRadius-log2 | - |
| Cytoplasm-AreaShape-MedianRadius-log2 | - |

*Continued on next page*

| Common features (inputs) | Variable features (outputs) |
|---|---|
| Cytoplasm-AreaShape-MinFeretDiameter-log2 | - |
| Cytoplasm-AreaShape-MinorAxisLength-log2 | - |
| Cytoplasm-AreaShape-Perimeter-log2 | - |
| Cytoplasm-AreaShape-Solidity-log2 | - |
| Cytoplasm-Granularity-10-ex405nm-raw | - |
| Cytoplasm-Granularity-10-ex647nm-raw | - |
| Cytoplasm-Granularity-11-ex405nm-raw | - |
| Cytoplasm-Granularity-11-ex647nm-raw | - |
| Cytoplasm-Granularity-12-ex405nm-raw | - |
| Cytoplasm-Granularity-12-ex647nm-raw | - |
| Cytoplasm-Granularity-13-ex405nm-raw | - |
| Cytoplasm-Granularity-13-ex647nm-raw | - |
| Cytoplasm-Granularity-14-ex405nm-raw | - |
| Cytoplasm-Granularity-14-ex647nm-raw | - |
| Cytoplasm-Granularity-15-ex405nm-raw | - |
| Cytoplasm-Granularity-15-ex647nm-raw | - |
| Cytoplasm-Granularity-16-ex405nm-raw | - |
| Cytoplasm-Granularity-16-ex647nm-raw | - |
| Cytoplasm-Granularity-1-ex405nm-raw | - |
| Cytoplasm-Granularity-1-ex647nm-raw | - |
| Cytoplasm-Granularity-2-ex405nm-raw | - |
| Cytoplasm-Granularity-2-ex647nm-raw | - |
| Cytoplasm-Granularity-3-ex405nm-raw | - |
| Cytoplasm-Granularity-3-ex647nm-raw | - |
| Cytoplasm-Granularity-4-ex405nm-raw | - |
| Cytoplasm-Granularity-4-ex647nm-raw | - |
| Cytoplasm-Granularity-5-ex405nm-raw | - |
| Cytoplasm-Granularity-5-ex647nm-raw | - |
| Cytoplasm-Granularity-6-ex405nm-raw | - |
| Cytoplasm-Granularity-6-ex647nm-raw | - |
| Cytoplasm-Granularity-7-ex405nm-raw | - |
| Cytoplasm-Granularity-7-ex647nm-raw | - |
| Cytoplasm-Granularity-8-ex405nm-raw | - |
| Cytoplasm-Granularity-8-ex647nm-raw | - |
| Cytoplasm-Granularity-9-ex405nm-raw | - |

*Continued on next page*

| Common features (inputs) | Variable features (outputs) |
| --- | --- |
| Cytoplasm-Granularity-9-ex647nm-raw | - |
| Cytoplasm-Intensity-IntegratedIntensityEdge-ex405nm-log2 | - |
| Cytoplasm-Intensity-IntegratedIntensityEdge-ex647nm-log2 | - |
| Cytoplasm-Intensity-IntegratedIntensity-ex405nm-log2 | - |
| Cytoplasm-Intensity-IntegratedIntensity-ex647nm-log2 | - |
| Cytoplasm-Intensity-LowerQuartileIntensity-ex405nm-log2 | - |
| Cytoplasm-Intensity-LowerQuartileIntensity-ex647nm-log2 | - |
| Cytoplasm-Intensity-MADIntensity-ex405nm-log2 | - |
| Cytoplasm-Intensity-MADIntensity-ex647nm-log2 | - |
| Cytoplasm-Intensity-MassDisplacement-ex405nm-log2 | - |
| Cytoplasm-Intensity-MassDisplacement-ex647nm-log2 | - |
| Cytoplasm-Intensity-MaxIntensityEdge-ex405nm-log2 | - |
| Cytoplasm-Intensity-MaxIntensityEdge-ex647nm-log2 | - |
| Cytoplasm-Intensity-MaxIntensity-ex405nm-log2 | - |
| Cytoplasm-Intensity-MaxIntensity-ex647nm-log2 | - |
| Cytoplasm-Intensity-MeanIntensityEdge-ex405nm-log2 | - |
| Cytoplasm-Intensity-MeanIntensityEdge-ex647nm-log2 | - |

*Continued on next page*

| Common features (inputs) | Variable features (outputs) |
|---|---|
| Cytoplasm-Intensity-MeanIntensity-ex405nm-log2 | - |
| Cytoplasm-Intensity-MeanIntensity-ex647nm-log2 | - |
| Cytoplasm-Intensity-MedianIntensity-ex405nm-log2 | - |
| Cytoplasm-Intensity-MedianIntensity-ex647nm-log2 | - |
| Cytoplasm-Intensity-MinIntensityEdge-ex405nm-log2 | - |
| Cytoplasm-Intensity-MinIntensityEdge-ex647nm-log2 | - |
| Cytoplasm-Intensity-MinIntensity-ex405nm-log2 | - |
| Cytoplasm-Intensity-MinIntensity-ex647nm-log2 | - |
| Cytoplasm-Intensity-StdIntensityEdge-ex405nm-log2 | - |
| Cytoplasm-Intensity-StdIntensityEdge-ex647nm-log2 | - |
| Cytoplasm-Intensity-StdIntensity-ex405nm-log2 | - |
| Cytoplasm-Intensity-StdIntensity-ex647nm-log2 | - |
| Cytoplasm-Intensity-UpperQuartileIntensity-ex405nm-log2 | - |
| Cytoplasm-Intensity-UpperQuartileIntensity-ex647nm-log2 | - |
| Nuclei-AreaShape-Area-log2 | - |
| Nuclei-AreaShape-BoundingBoxArea-log2 | - |
| Nuclei-AreaShape-Compactness-log2 | - |
| Nuclei-AreaShape-ConvexArea-log2 | - |
| Nuclei-AreaShape-Eccentricity-raw | - |
| Nuclei-AreaShape-EquivalentDiameter-log2 | - |

*Continued on next page*

| Common features (inputs) | Variable features (outputs) |
|---|---|
| Nuclei-AreaShape-Extent-raw | - |
| Nuclei-AreaShape-FormFactor-raw | - |
| Nuclei-AreaShape-MajorAxisLength-log2 | - |
| Nuclei-AreaShape-MaxFeretDiameter-log2 | - |
| Nuclei-AreaShape-MaximumRadius-log2 | - |
| Nuclei-AreaShape-MeanRadius-log2 | - |
| Nuclei-AreaShape-MedianRadius-log2 | - |
| Nuclei-AreaShape-MinFeretDiameter-log2 | - |
| Nuclei-AreaShape-MinorAxisLength-log2 | - |
| Nuclei-AreaShape-Perimeter-log2 | - |
| Nuclei-AreaShape-Solidity-log2 | - |
| Nuclei-Granularity-10-ex405nm-raw | - |
| Nuclei-Granularity-10-ex647nm-raw | - |
| Nuclei-Granularity-11-ex405nm-raw | - |
| Nuclei-Granularity-11-ex647nm-raw | - |
| Nuclei-Granularity-12-ex405nm-raw | - |
| Nuclei-Granularity-12-ex647nm-raw | - |
| Nuclei-Granularity-13-ex405nm-raw | - |
| Nuclei-Granularity-13-ex647nm-raw | - |
| Nuclei-Granularity-14-ex405nm-raw | - |
| Nuclei-Granularity-14-ex647nm-raw | - |
| Nuclei-Granularity-15-ex405nm-raw | - |
| Nuclei-Granularity-15-ex647nm-raw | - |
| Nuclei-Granularity-16-ex405nm-raw | - |
| Nuclei-Granularity-16-ex647nm-raw | - |
| Nuclei-Granularity-1-ex405nm-raw | - |
| Nuclei-Granularity-1-ex647nm-raw | - |
| Nuclei-Granularity-2-ex405nm-raw | - |
| Nuclei-Granularity-2-ex647nm-raw | - |
| Nuclei-Granularity-3-ex405nm-raw | - |
| Nuclei-Granularity-3-ex647nm-raw | - |
| Nuclei-Granularity-4-ex405nm-raw | - |
| Nuclei-Granularity-4-ex647nm-raw | - |
| Nuclei-Granularity-5-ex405nm-raw | - |
| Nuclei-Granularity-5-ex647nm-raw | - |
| Nuclei-Granularity-6-ex405nm-raw | - |

*Continued on next page*

| Common features (inputs) | Variable features (outputs) |
|---|---|
| Nuclei-Granularity-6-ex647nm-raw | - |
| Nuclei-Granularity-7-ex405nm-raw | - |
| Nuclei-Granularity-7-ex647nm-raw | - |
| Nuclei-Granularity-8-ex405nm-raw | - |
| Nuclei-Granularity-8-ex647nm-raw | - |
| Nuclei-Granularity-9-ex405nm-raw | - |
| Nuclei-Granularity-9-ex647nm-raw | - |
| Nuclei-Intensity-IntegratedIntensityEdge-ex405nm-log2 | - |
| Nuclei-Intensity-IntegratedIntensityEdge-ex647nm-log2 | - |
| Nuclei-Intensity-IntegratedIntensity-ex405nm-log2 | - |
| Nuclei-Intensity-IntegratedIntensity-ex647nm-log2 | - |
| Nuclei-Intensity-LowerQuartileIntensity-ex405nm-log2 | - |
| Nuclei-Intensity-LowerQuartileIntensity-ex647nm-log2 | - |
| Nuclei-Intensity-MADIntensity-ex405nm-log2 | - |
| Nuclei-Intensity-MADIntensity-ex647nm-log2 | - |
| Nuclei-Intensity-MassDisplacement-ex405nm-log2 | - |
| Nuclei-Intensity-MassDisplacement-ex647nm-log2 | - |
| Nuclei-Intensity-MaxIntensityEdge-ex405nm-log2 | - |
| Nuclei-Intensity-MaxIntensityEdge-ex647nm-log2 | - |
| Nuclei-Intensity-MaxIntensity-ex405nm-log2 | - |
| Nuclei-Intensity-MaxIntensity-ex647nm-log2 | - |
| Nuclei-Intensity-MeanIntensityEdge-ex405nm-log2 | - |

*Continued on next page*

| Common features (inputs) | Variable features (outputs) |
|---|---|
| Nuclei-Intensity-MeanIntensityEdge-ex647nm-log2 | - |
| Nuclei-Intensity-MeanIntensity-ex405nm-log2 | - |
| Nuclei-Intensity-MeanIntensity-ex647nm-log2 | - |
| Nuclei-Intensity-MedianIntensity-ex405nm-log2 | - |
| Nuclei-Intensity-MedianIntensity-ex647nm-log2 | - |
| Nuclei-Intensity-MinIntensityEdge-ex405nm-log2 | - |
| Nuclei-Intensity-MinIntensityEdge-ex647nm-log2 | - |
| Nuclei-Intensity-MinIntensity-ex405nm-log2 | - |
| Nuclei-Intensity-MinIntensity-ex647nm-log2 | - |
| Nuclei-Intensity-StdIntensityEdge-ex405nm-log2 | - |
| Nuclei-Intensity-StdIntensityEdge-ex647nm-log2 | - |
| Nuclei-Intensity-StdIntensity-ex405nm-log2 | - |
| Nuclei-Intensity-StdIntensity-ex647nm-log2 | - |
| Nuclei-Intensity-UpperQuartileIntensity-ex405nm-log2 | - |
| Nuclei-Intensity-UpperQuartileIntensity-ex647nm-log2 | - |
| Nuclei-Texture-AngularSecondMoment-ex405nm-3-00-256-log2 | - |
| Nuclei-Texture-AngularSecondMoment-ex405nm-3-01-256-log2 | - |
| Nuclei-Texture-AngularSecondMoment-ex405nm-3-02-256-log2 | - |
| Nuclei-Texture-AngularSecondMoment-ex405nm-3-03-256-log2 | - |
| Nuclei-Texture-AngularSecondMoment-ex647nm-3-00-256-log2 | - |

*Continued on next page*

| Common features (inputs) | Variable features (outputs) |
|---|---|
| Nuclei-Texture-AngularSecondMoment-ex647nm-3-01-256-log2 | - |
| Nuclei-Texture-AngularSecondMoment-ex647nm-3-02-256-log2 | - |
| Nuclei-Texture-AngularSecondMoment-ex647nm-3-03-256-log2 | - |
| Nuclei-Texture-Contrast-ex405nm-3-00-256-sqrt | - |
| Nuclei-Texture-Contrast-ex405nm-3-01-256-sqrt | - |
| Nuclei-Texture-Contrast-ex405nm-3-02-256-sqrt | - |
| Nuclei-Texture-Contrast-ex405nm-3-03-256-sqrt | - |
| Nuclei-Texture-Contrast-ex647nm-3-00-256-sqrt | - |
| Nuclei-Texture-Contrast-ex647nm-3-01-256-sqrt | - |
| Nuclei-Texture-Contrast-ex647nm-3-02-256-sqrt | - |
| Nuclei-Texture-Contrast-ex647nm-3-03-256-sqrt | - |
| Nuclei-Texture-Correlation-ex405nm-3-00-256-raw | - |
| Nuclei-Texture-Correlation-ex405nm-3-01-256-raw | - |
| Nuclei-Texture-Correlation-ex405nm-3-02-256-raw | - |
| Nuclei-Texture-Correlation-ex405nm-3-03-256-raw | - |
| Nuclei-Texture-Correlation-ex647nm-3-00-256-raw | - |
| Nuclei-Texture-Correlation-ex647nm-3-01-256-raw | - |

*Continued on next page*

| Common features (inputs) | Variable features (outputs) |
|---|---|
| Nuclei-Texture-Correlation-ex647nm-3-02-256-raw | - |
| Nuclei-Texture-Correlation-ex647nm-3-03-256-raw | - |
| Nuclei-Texture-DifferenceEntropy-ex405nm-3-00-256-raw | - |
| Nuclei-Texture-DifferenceEntropy-ex405nm-3-01-256-raw | - |
| Nuclei-Texture-DifferenceEntropy-ex405nm-3-02-256-raw | - |
| Nuclei-Texture-DifferenceEntropy-ex405nm-3-03-256-raw | - |
| Nuclei-Texture-DifferenceEntropy-ex647nm-3-00-256-raw | - |
| Nuclei-Texture-DifferenceEntropy-ex647nm-3-01-256-raw | - |
| Nuclei-Texture-DifferenceEntropy-ex647nm-3-02-256-raw | - |
| Nuclei-Texture-DifferenceEntropy-ex647nm-3-03-256-raw | - |
| Nuclei-Texture-DifferenceVariance-ex405nm-3-00-256-log2 | - |
| Nuclei-Texture-DifferenceVariance-ex405nm-3-01-256-log2 | - |
| Nuclei-Texture-DifferenceVariance-ex405nm-3-02-256-log2 | - |
| Nuclei-Texture-DifferenceVariance-ex405nm-3-03-256-log2 | - |
| Nuclei-Texture-DifferenceVariance-ex647nm-3-00-256-log2 | - |
| Nuclei-Texture-DifferenceVariance-ex647nm-3-01-256-log2 | - |
| Nuclei-Texture-DifferenceVariance-ex647nm-3-02-256-log2 | - |

*Continued on next page*

| Common features (inputs) | Variable features (outputs) |
|---|---|
| Nuclei-Texture-DifferenceVariance-ex647nm-3-03-256-log2 | - |
| Nuclei-Texture-Entropy-ex405nm-3-00-256-raw | - |
| Nuclei-Texture-Entropy-ex405nm-3-01-256-raw | - |
| Nuclei-Texture-Entropy-ex405nm-3-02-256-raw | - |
| Nuclei-Texture-Entropy-ex405nm-3-03-256-raw | - |
| Nuclei-Texture-Entropy-ex647nm-3-00-256-raw | - |
| Nuclei-Texture-Entropy-ex647nm-3-01-256-raw | - |
| Nuclei-Texture-Entropy-ex647nm-3-02-256-raw | - |
| Nuclei-Texture-Entropy-ex647nm-3-03-256-raw | - |
| Nuclei-Texture-InfoMeas1-ex405nm-3-00-256-raw | - |
| Nuclei-Texture-InfoMeas1-ex405nm-3-01-256-raw | - |
| Nuclei-Texture-InfoMeas1-ex405nm-3-02-256-raw | - |
| Nuclei-Texture-InfoMeas1-ex647nm-3-00-256-raw | - |
| Nuclei-Texture-InfoMeas1-ex647nm-3-01-256-raw | - |
| Nuclei-Texture-InfoMeas1-ex647nm-3-02-256-raw | - |
| Nuclei-Texture-InfoMeas1-ex647nm-3-03-256-raw | - |
| Nuclei-Texture-InfoMeas2-ex405nm-3-00-256-raw | - |

*Continued on next page*

| Common features (inputs) | Variable features (outputs) |
|---|---|
| Nuclei-Texture-InfoMeas2-ex405nm-3-01-256-raw | - |
| Nuclei-Texture-InfoMeas2-ex405nm-3-02-256-raw | - |
| Nuclei-Texture-InfoMeas2-ex405nm-3-03-256-raw | - |
| Nuclei-Texture-InfoMeas2-ex647nm-3-00-256-raw | - |
| Nuclei-Texture-InfoMeas2-ex647nm-3-01-256-raw | - |
| Nuclei-Texture-InfoMeas2-ex647nm-3-02-256-raw | - |
| Nuclei-Texture-InfoMeas2-ex647nm-3-03-256-raw | - |
| Nuclei-Texture-InverseDifferenceMoment-ex405nm-3-00-256-raw | - |
| Nuclei-Texture-InverseDifferenceMoment-ex405nm-3-01-256-raw | - |
| Nuclei-Texture-InverseDifferenceMoment-ex405nm-3-02-256-raw | - |
| Nuclei-Texture-InverseDifferenceMoment-ex405nm-3-03-256-raw | - |
| Nuclei-Texture-InverseDifferenceMoment-ex647nm-3-00-256-raw | - |
| Nuclei-Texture-InverseDifferenceMoment-ex647nm-3-01-256-raw | - |
| Nuclei-Texture-InverseDifferenceMoment-ex647nm-3-02-256-raw | - |
| Nuclei-Texture-InverseDifferenceMoment-ex647nm-3-03-256-raw | - |
| Nuclei-Texture-SumAverage-ex405nm-3-00-256-log2 | - |
| Nuclei-Texture-SumAverage-ex405nm-3-01-256-log2 | - |

*Continued on next page*

| Common features (inputs) | Variable features (outputs) |
|---|---|
| Nuclei-Texture-SumAverage-ex405nm-3-02-256-log2 | - |
| Nuclei-Texture-SumAverage-ex405nm-3-03-256-log2 | - |
| Nuclei-Texture-SumAverage-ex647nm-3-00-256-log2 | - |
| Nuclei-Texture-SumAverage-ex647nm-3-01-256-log2 | - |
| Nuclei-Texture-SumAverage-ex647nm-3-02-256-log2 | - |
| Nuclei-Texture-SumAverage-ex647nm-3-03-256-log2 | - |
| Nuclei-Texture-SumEntropy-ex405nm-3-00-256-raw | - |
| Nuclei-Texture-SumEntropy-ex405nm-3-01-256-raw | - |
| Nuclei-Texture-SumEntropy-ex405nm-3-02-256-raw | - |
| Nuclei-Texture-SumEntropy-ex405nm-3-03-256-raw | - |
| Nuclei-Texture-SumEntropy-ex647nm-3-00-256-raw | - |
| Nuclei-Texture-SumEntropy-ex647nm-3-01-256-raw | - |
| Nuclei-Texture-SumEntropy-ex647nm-3-02-256-raw | - |
| Nuclei-Texture-SumEntropy-ex647nm-3-03-256-raw | - |
| Nuclei-Texture-SumVariance-ex405nm-3-00-256-sqrt | - |
| Nuclei-Texture-SumVariance-ex405nm-3-01-256-sqrt | - |
| Nuclei-Texture-SumVariance-ex405nm-3-02-256-sqrt | - |

*Continued on next page*

| Common features (inputs) | Variable features (outputs) |
|---|---|
| Nuclei-Texture-SumVariance-ex405nm-3-03-256-sqrt | - |
| Nuclei-Texture-SumVariance-ex647nm-3-00-256-sqrt | - |
| Nuclei-Texture-SumVariance-ex647nm-3-01-256-sqrt | - |
| Nuclei-Texture-SumVariance-ex647nm-3-02-256-sqrt | - |
| Nuclei-Texture-SumVariance-ex647nm-3-03-256-sqrt | - |
| Nuclei-Texture-Variance-ex405nm-3-00-256-sqrt | - |
| Nuclei-Texture-Variance-ex405nm-3-01-256-sqrt | - |
| Nuclei-Texture-Variance-ex405nm-3-02-256-sqrt | - |
| Nuclei-Texture-Variance-ex405nm-3-03-256-sqrt | - |
| Nuclei-Texture-Variance-ex647nm-3-00-256-sqrt | - |
| Nuclei-Texture-Variance-ex647nm-3-01-256-sqrt | - |
| Nuclei-Texture-Variance-ex647nm-3-02-256-sqrt | - |
| Nuclei-Texture-Variance-ex647nm-3-03-256-sqrt | - |
| Ratio-NC-ex647nm-IntegInt-log2 | - |
| Ratio-NC-ex647nm-MeanInt-log2 | - |
| X007-Categ-TreatmentTime-0.min | - |
| X007-Categ-TreatmentTime-2.min | - |
| X007-Categ-TreatmentTime-20.min | - |
| X007-Categ-TreatmentTime-5.min | - |
| X007-Categ-TreatmentTime-60.min | - |

*Continued on next page*

Table S2: Features used for dimensionality reduction of epithelial–mesenchymal (E/M) and signaling landscapes

| **Features used for E/M landscape UMAP** | **Features used for signaling landscape PHATE** |
|---|---|
| CellBody-AreaShape-Area-log2 | p-ERK-CellBody-Intensity-IntegratedIntensityEdge-ex555nm-log2-predicted |
| CellBody-AreaShape-BoundingBoxArea-log2 | p-ERK-CellBody-Intensity-IntegratedIntensity-ex555nm-log2-predicted |
| CellBody-AreaShape-Compactness-log2 | p-ERK-CellBody-Intensity-LowerQuartileIntensity-ex555nm-log2-predicted |
| CellBody-AreaShape-ConvexArea-log2 | p-ERK-CellBody-Intensity-MADIntensity-ex555nm-log2-predicted |
| CellBody-AreaShape-Eccentricity-raw | p-ERK-CellBody-Intensity-MaxIntensityEdge-ex555nm-log2-predicted |
| CellBody-AreaShape-EquivalentDiameter-log2 | p-ERK-CellBody-Intensity-MaxIntensity-ex555nm-log2-predicted |
| CellBody-AreaShape-Extent-raw | p-ERK-CellBody-Intensity-MeanIntensityEdge-ex555nm-log2-predicted |
| CellBody-AreaShape-FormFactor-raw | p-ERK-CellBody-Intensity-MeanIntensity-ex555nm-log2-predicted |
| CellBody-AreaShape-MajorAxisLength-log2 | p-ERK-CellBody-Intensity-MedianIntensity-ex555nm-log2-predicted |
| CellBody-AreaShape-MaxFeretDiameter-log2 | p-ERK-CellBody-Intensity-MinIntensityEdge-ex555nm-log2-predicted |
| CellBody-AreaShape-MaximumRadius-log2 | p-ERK-CellBody-Intensity-MinIntensity-ex555nm-log2-predicted |
| CellBody-AreaShape-MeanRadius-log2 | p-ERK-CellBody-Intensity-StdIntensityEdge-ex555nm-log2-predicted |
| CellBody-AreaShape-MedianRadius-log2 | p-ERK-CellBody-Intensity-StdIntensity-ex555nm-log2-predicted |

*Continued on next page*

| Features used for E/M landscape UMAP | Features used for signaling landscape PHATE |
|---|---|
| CellBody-AreaShape-MinFeretDiameter-log2 | p-ERK-CellBody-Intensity-UpperQuartileIntensity-ex555nm-log2-predicted |
| CellBody-AreaShape-MinorAxisLength-log2 | p-ERK-CellBody-RadialDistribution-FracAtD-ex555nm-1of4-log2-predicted |
| CellBody-AreaShape-Perimeter-log2 | p-ERK-CellBody-RadialDistribution-FracAtD-ex555nm-2of4-log2-predicted |
| CellBody-AreaShape-Solidity-log2 | p-ERK-CellBody-RadialDistribution-FracAtD-ex555nm-3of4-log2-predicted |
| | p-ERK-CellBody-RadialDistribution-FracAtD-ex555nm-4of4-log2-predicted |
| | p-ERK-CellBody-RadialDistribution-MeanFrac-ex555nm-1of4-log2-predicted |
| | p-ERK-CellBody-RadialDistribution-MeanFrac-ex555nm-2of4-log2-predicted |
| | p-ERK-CellBody-RadialDistribution-MeanFrac-ex555nm-3of4-log2-predicted |
| | p-ERK-CellBody-RadialDistribution-MeanFrac-ex555nm-4of4-log2-predicted |
| | p-ERK-CellBody-RadialDistribution-RadialCV-ex555nm-4of4-log2-predicted |
| | p-ERK-CellBody-RadialDistribution-ZernikeMagnitude-ex555nm-0-0-log2-predicted |
| | p-ERK-CellBody-RadialDistribution-ZernikeMagnitude-ex555nm-1-1-log2-predicted |
| | p-ERK-CellBody-RadialDistribution-ZernikeMagnitude-ex555nm-2-0-log2-predicted |

*Continued on next page*

| Features used for E/M landscape UMAP | Features used for signaling landscape PHATE |
|---|---|
| | p-ERK-CellBody-RadialDistribution-ZernikeMagnitude-ex555nm-2-2-log2-predicted |
| | p-ERK-CellBody-RadialDistribution-ZernikeMagnitude-ex555nm-3-1-log2-predicted |
| | p-ERK-CellBody-RadialDistribution-ZernikeMagnitude-ex555nm-3-3-log2-predicted |
| | p-ERK-CellBody-RadialDistribution-ZernikeMagnitude-ex555nm-4-2-log2-predicted |
| | p-ERK-CellBody-RadialDistribution-ZernikeMagnitude-ex555nm-4-4-log2-predicted |
| | p-ERK-CellBody-RadialDistribution-ZernikeMagnitude-ex555nm-5-1-log2-predicted |
| | p-ERK-CellBody-RadialDistribution-ZernikeMagnitude-ex555nm-5-3-log2-predicted |
| | p-ERK-CellBody-RadialDistribution-ZernikeMagnitude-ex555nm-5-5-log2-predicted |
| | p-ERK-CellBody-RadialDistribution-ZernikeMagnitude-ex555nm-6-4-log2-predicted |
| | p-ERK-CellBody-RadialDistribution-ZernikeMagnitude-ex555nm-6-6-log2-predicted |

*Continued on next page*

| Features used for E/M landscape UMAP | Features used for signaling landscape PHATE |
|---|---|
| | p-ERK-CellBody-RadialDistribution-ZernikeMagnitude-ex555nm-7-1-log2-predicted |
| | p-ERK-CellBody-RadialDistribution-ZernikeMagnitude-ex555nm-7-5-log2-predicted |
| | p-ERK-CellBody-RadialDistribution-ZernikeMagnitude-ex555nm-7-7-log2-predicted |
| | p-ERK-CellBody-RadialDistribution-ZernikeMagnitude-ex555nm-8-6-log2-predicted |
| | p-ERK-CellBody-RadialDistribution-ZernikeMagnitude-ex555nm-8-8-log2-predicted |
| | p-ERK-CellBody-RadialDistribution-ZernikeMagnitude-ex555nm-9-1-log2-predicted |
| | p-ERK-CellBody-RadialDistribution-ZernikeMagnitude-ex555nm-9-7-log2-predicted |
| | p-ERK-CellBody-RadialDistribution-ZernikeMagnitude-ex555nm-9-9-log2-predicted |
| | p-ERK-CellBody-Texture-AngularSecondMoment-ex555nm-3-00-256-log2-predicted |
| | p-ERK-CellBody-Texture-AngularSecondMoment-ex555nm-3-01-256-log2-predicted |

*Continued on next page*

| Features used for E/M landscape UMAP | Features used for signaling landscape PHATE |
| --- | --- |
| | p-ERK-CellBody-Texture-AngularSecondMoment-ex555nm-3-02-256-log2-predicted |
| | p-ERK-CellBody-Texture-AngularSecondMoment-ex555nm-3-03-256-log2-predicted |
| | p-ERK-CellBody-Texture-Contrast-ex555nm-3-00-256-sqrt-predicted |
| | p-ERK-CellBody-Texture-Contrast-ex555nm-3-01-256-sqrt-predicted |
| | p-ERK-CellBody-Texture-Contrast-ex555nm-3-02-256-sqrt-predicted |
| | p-ERK-CellBody-Texture-Contrast-ex555nm-3-03-256-sqrt-predicted |
| | p-ERK-CellBody-Texture-Correlation-ex555nm-3-00-256-raw-predicted |
| | p-ERK-CellBody-Texture-Correlation-ex555nm-3-01-256-raw-predicted |
| | p-ERK-CellBody-Texture-Correlation-ex555nm-3-02-256-raw-predicted |
| | p-ERK-CellBody-Texture-Correlation-ex555nm-3-03-256-raw-predicted |
| | p-ERK-CellBody-Texture-DifferenceEntropy-ex555nm-3-00-256-raw-predicted |
| | p-ERK-CellBody-Texture-DifferenceEntropy-ex555nm-3-01-256-raw-predicted |
| | p-ERK-CellBody-Texture-DifferenceEntropy-ex555nm-3-02-256-raw-predicted |
| | p-ERK-CellBody-Texture-DifferenceEntropy-ex555nm-3-03-256-raw-predicted |
| | p-ERK-CellBody-Texture-DifferenceVariance-ex555nm-3-00-256-log2-predicted |

*Continued on next page*

| Features used for E/M landscape UMAP | Features used for signaling landscape PHATE |
|---|---|
| | p-ERK-CellBody-Texture-DifferenceVariance-ex555nm-3-01-256-log2-predicted |
| | p-ERK-CellBody-Texture-DifferenceVariance-ex555nm-3-02-256-log2-predicted |
| | p-ERK-CellBody-Texture-DifferenceVariance-ex555nm-3-03-256-log2-predicted |
| | p-ERK-CellBody-Texture-Entropy-ex555nm-3-00-256-raw-predicted |
| | p-ERK-CellBody-Texture-Entropy-ex555nm-3-01-256-raw-predicted |
| | p-ERK-CellBody-Texture-Entropy-ex555nm-3-02-256-raw-predicted |
| | p-ERK-CellBody-Texture-Entropy-ex555nm-3-03-256-raw-predicted |
| | p-ERK-CellBody-Texture-InfoMeas1-ex555nm-3-00-256-raw-predicted |
| | p-ERK-CellBody-Texture-InfoMeas1-ex555nm-3-01-256-raw-predicted |
| | p-ERK-CellBody-Texture-InfoMeas1-ex555nm-3-02-256-raw-predicted |
| | p-ERK-CellBody-Texture-InfoMeas1-ex555nm-3-03-256-raw-predicted |
| | p-ERK-CellBody-Texture-InfoMeas2-ex555nm-3-00-256-raw-predicted |
| | p-ERK-CellBody-Texture-InfoMeas2-ex555nm-3-01-256-raw-predicted |
| | p-ERK-CellBody-Texture-InfoMeas2-ex555nm-3-02-256-raw-predicted |
| | p-ERK-CellBody-Texture-InfoMeas2-ex555nm-3-03-256-raw-predicted |
| | p-ERK-CellBody-Texture-InverseDifferenceMoment-ex555nm-3-00-256-raw-predicted |

*Continued on next page*

| Features used for E/M landscape UMAP | Features used for signaling landscape PHATE |
|---|---|
| | p-ERK-CellBody-Texture-InverseDifferenceMoment-ex555nm-3-01-256-raw-predicted |
| | p-ERK-CellBody-Texture-InverseDifferenceMoment-ex555nm-3-02-256-raw-predicted |
| | p-ERK-CellBody-Texture-InverseDifferenceMoment-ex555nm-3-03-256-raw-predicted |
| | p-ERK-CellBody-Texture-SumAverage-ex555nm-3-00-256-log2-predicted |
| | p-ERK-CellBody-Texture-SumAverage-ex555nm-3-01-256-log2-predicted |
| | p-ERK-CellBody-Texture-SumAverage-ex555nm-3-02-256-log2-predicted |
| | p-ERK-CellBody-Texture-SumAverage-ex555nm-3-03-256-log2-predicted |
| | p-ERK-CellBody-Texture-SumEntropy-ex555nm-3-00-256-raw-predicted |
| | p-ERK-CellBody-Texture-SumEntropy-ex555nm-3-01-256-raw-predicted |
| | p-ERK-CellBody-Texture-SumEntropy-ex555nm-3-02-256-raw-predicted |
| | p-ERK-CellBody-Texture-SumEntropy-ex555nm-3-03-256-raw-predicted |
| | p-ERK-CellBody-Texture-SumVariance-ex555nm-3-00-256-sqrt-predicted |
| | p-ERK-CellBody-Texture-SumVariance-ex555nm-3-01-256-sqrt-predicted |
| | p-ERK-CellBody-Texture-SumVariance-ex555nm-3-02-256-sqrt-predicted |
| | p-ERK-CellBody-Texture-SumVariance-ex555nm-3-03-256-sqrt-predicted |

*Continued on next page*

| Features used for E/M landscape UMAP | Features used for signaling landscape PHATE |
| --- | --- |
| | p-ERK-CellBody-Texture-Variance-ex555nm-3-00-256-sqrt-predicted |
| | p-ERK-CellBody-Texture-Variance-ex555nm-3-01-256-sqrt-predicted |
| | p-ERK-CellBody-Texture-Variance-ex555nm-3-02-256-sqrt-predicted |
| | p-ERK-CellBody-Texture-Variance-ex555nm-3-03-256-sqrt-predicted |
| | p-ERK-Cytoplasm-Intensity-IntegratedIntensityEdge-ex555nm-log2-predicted |
| | p-ERK-Cytoplasm-Intensity-IntegratedIntensity-ex555nm-log2-predicted |
| | p-ERK-Cytoplasm-Intensity-LowerQuartileIntensity-ex555nm-log2-predicted |
| | p-ERK-Cytoplasm-Intensity-MADIntensity-ex555nm-log2-predicted |
| | p-ERK-Cytoplasm-Intensity-MaxIntensityEdge-ex555nm-log2-predicted |
| | p-ERK-Cytoplasm-Intensity-MaxIntensity-ex555nm-log2-predicted |
| | p-ERK-Cytoplasm-Intensity-MeanIntensityEdge-ex555nm-log2-predicted |
| | p-ERK-Cytoplasm-Intensity-MeanIntensity-ex555nm-log2-predicted |
| | p-ERK-Cytoplasm-Intensity-MedianIntensity-ex555nm-log2-predicted |
| | p-ERK-Cytoplasm-Intensity-MinIntensityEdge-ex555nm-log2-predicted |
| | p-ERK-Cytoplasm-Intensity-MinIntensity-ex555nm-log2-predicted |

*Continued on next page*

| Features used for E/M landscape UMAP | Features used for signaling landscape PHATE |
| --- | --- |
| | p-ERK-Cytoplasm-Intensity-StdIntensityEdge-ex555nm-log2-predicted |
| | p-ERK-Cytoplasm-Intensity-StdIntensity-ex555nm-log2-predicted |
| | p-ERK-Cytoplasm-Intensity-UpperQuartileIntensity-ex555nm-log2-predicted |
| | p-ERK-Nuclei-Intensity-IntegratedIntensityEdge-ex555nm-log2-predicted |
| | p-ERK-Nuclei-Intensity-IntegratedIntensity-ex555nm-log2-predicted |
| | p-ERK-Nuclei-Intensity-LowerQuartileIntensity-ex555nm-log2-predicted |
| | p-ERK-Nuclei-Intensity-MADIntensity-ex555nm-log2-predicted |
| | p-ERK-Nuclei-Intensity-MaxIntensityEdge-ex555nm-log2-predicted |
| | p-ERK-Nuclei-Intensity-MaxIntensity-ex555nm-log2-predicted |
| | p-ERK-Nuclei-Intensity-MeanIntensityEdge-ex555nm-log2-predicted |
| | p-ERK-Nuclei-Intensity-MeanIntensity-ex555nm-log2-predicted |
| | p-ERK-Nuclei-Intensity-MedianIntensity-ex555nm-log2-predicted |
| | p-ERK-Nuclei-Intensity-MinIntensityEdge-ex555nm-log2-predicted |
| | p-ERK-Nuclei-Intensity-MinIntensity-ex555nm-log2-predicted |
| | p-ERK-Nuclei-Intensity-StdIntensityEdge-ex555nm-log2-predicted |

*Continued on next page*

| Features used for E/M landscape UMAP | Features used for signaling landscape PHATE |
|---|---|
| | p-ERK-Nuclei-Intensity-StdIntensity-ex555nm-log2-predicted |
| | p-ERK-Nuclei-Intensity-UpperQuartileIntensity-ex555nm-log2-predicted |
| | p-ERK-Nuclei-Texture-AngularSecondMoment-ex555nm-3-00-256-log2-predicted |
| | p-ERK-Nuclei-Texture-AngularSecondMoment-ex555nm-3-01-256-log2-predicted |
| | p-ERK-Nuclei-Texture-AngularSecondMoment-ex555nm-3-02-256-log2-predicted |
| | p-ERK-Nuclei-Texture-AngularSecondMoment-ex555nm-3-03-256-log2-predicted |
| | p-ERK-Nuclei-Texture-Contrast-ex555nm-3-00-256-sqrt-predicted |
| | p-ERK-Nuclei-Texture-Contrast-ex555nm-3-01-256-sqrt-predicted |
| | p-ERK-Nuclei-Texture-Contrast-ex555nm-3-02-256-sqrt-predicted |
| | p-ERK-Nuclei-Texture-Contrast-ex555nm-3-03-256-sqrt-predicted |
| | p-ERK-Nuclei-Texture-Correlation-ex555nm-3-00-256-raw-predicted |
| | p-ERK-Nuclei-Texture-Correlation-ex555nm-3-02-256-raw-predicted |
| | p-ERK-Nuclei-Texture-DifferenceEntropy-ex555nm-3-00-256-raw-predicted |

*Continued on next page*

| Features used for E/M landscape UMAP | Features used for signaling landscape PHATE |
|---|---|
| | p-ERK-Nuclei-Texture-DifferenceEntropy-ex555nm-3-01-256-raw-predicted |
| | p-ERK-Nuclei-Texture-DifferenceEntropy-ex555nm-3-02-256-raw-predicted |
| | p-ERK-Nuclei-Texture-DifferenceEntropy-ex555nm-3-03-256-raw-predicted |
| | p-ERK-Nuclei-Texture-DifferenceVariance-ex555nm-3-00-256-log2-predicted |
| | p-ERK-Nuclei-Texture-DifferenceVariance-ex555nm-3-01-256-log2-predicted |
| | p-ERK-Nuclei-Texture-DifferenceVariance-ex555nm-3-02-256-log2-predicted |
| | p-ERK-Nuclei-Texture-DifferenceVariance-ex555nm-3-03-256-log2-predicted |
| | p-ERK-Nuclei-Texture-Entropy-ex555nm-3-00-256-raw-predicted |
| | p-ERK-Nuclei-Texture-Entropy-ex555nm-3-01-256-raw-predicted |
| | p-ERK-Nuclei-Texture-Entropy-ex555nm-3-02-256-raw-predicted |
| | p-ERK-Nuclei-Texture-Entropy-ex555nm-3-03-256-raw-predicted |
| | p-ERK-Nuclei-Texture-InfoMeas2-ex555nm-3-00-256-raw-predicted |
| | p-ERK-Nuclei-Texture-InfoMeas2-ex555nm-3-01-256-raw-predicted |
| | p-ERK-Nuclei-Texture-InfoMeas2-ex555nm-3-02-256-raw-predicted |
| | p-ERK-Nuclei-Texture-InfoMeas2-ex555nm-3-03-256-raw-predicted |
| | p-ERK-Nuclei-Texture-InverseDifferenceMoment-ex555nm-3-00-256-raw-predicted |

*Continued on next page*

| Features used for E/M landscape UMAP | Features used for signaling landscape PHATE |
|---|---|
| | p-ERK-Nuclei-Texture-InverseDifferenceMoment-ex555nm-3-01-256-raw-predicted |
| | p-ERK-Nuclei-Texture-InverseDifferenceMoment-ex555nm-3-02-256-raw-predicted |
| | p-ERK-Nuclei-Texture-InverseDifferenceMoment-ex555nm-3-03-256-raw-predicted |
| | p-ERK-Nuclei-Texture-SumAverage-ex555nm-3-00-256-log2-predicted |
| | p-ERK-Nuclei-Texture-SumAverage-ex555nm-3-01-256-log2-predicted |
| | p-ERK-Nuclei-Texture-SumAverage-ex555nm-3-02-256-log2-predicted |
| | p-ERK-Nuclei-Texture-SumAverage-ex555nm-3-03-256-log2-predicted |
| | p-ERK-Nuclei-Texture-SumEntropy-ex555nm-3-00-256-raw-predicted |
| | p-ERK-Nuclei-Texture-SumEntropy-ex555nm-3-01-256-raw-predicted |
| | p-ERK-Nuclei-Texture-SumEntropy-ex555nm-3-02-256-raw-predicted |
| | p-ERK-Nuclei-Texture-SumEntropy-ex555nm-3-03-256-raw-predicted |
| | p-ERK-Nuclei-Texture-SumVariance-ex555nm-3-00-256-sqrt-predicted |
| | p-ERK-Nuclei-Texture-SumVariance-ex555nm-3-01-256-sqrt-predicted |
| | p-ERK-Nuclei-Texture-SumVariance-ex555nm-3-02-256-sqrt-predicted |
| | p-ERK-Nuclei-Texture-SumVariance-ex555nm-3-03-256-sqrt-predicted |

*Continued on next page*

| Features used for E/M landscape UMAP | Features used for signaling landscape PHATE |
| --- | --- |
| | p-ERK-Nuclei-Texture-Variance-ex555nm-3-00-256-sqrt-predicted |
| | p-ERK-Nuclei-Texture-Variance-ex555nm-3-01-256-sqrt-predicted |
| | p-ERK-Nuclei-Texture-Variance-ex555nm-3-02-256-sqrt-predicted |
| | p-ERK-Nuclei-Texture-Variance-ex555nm-3-03-256-sqrt-predicted |
| | p-ERK-Ratio-NC-ex555nm-IntegInt-log2-predicted |
| | p-ERK-Ratio-NC-ex555nm-MeanInt-log2-predicted |
| | p-S6-CellBody-Intensity-IntegratedIntensityEdge-ex555nm-log2-predicted |
| | p-S6-CellBody-Intensity-IntegratedIntensity-ex555nm-log2-predicted |
| | p-S6-CellBody-Intensity-LowerQuartileIntensity-ex555nm-log2-predicted |
| | p-S6-CellBody-Intensity-MeanIntensityEdge-ex555nm-log2-predicted |
| | p-S6-CellBody-Intensity-MinIntensityEdge-ex555nm-log2-predicted |
| | p-S6-CellBody-Intensity-StdIntensityEdge-ex555nm-log2-predicted |
| | p-S6-CellBody-RadialDistribution-FracAtD-ex555nm-1of4-log2-predicted |
| | p-S6-CellBody-RadialDistribution-FracAtD-ex555nm-2of4-log2-predicted |
| | p-S6-CellBody-RadialDistribution-FracAtD-ex555nm-3of4-log2-predicted |

*Continued on next page*

| Features used for E/M landscape UMAP | Features used for signaling landscape PHATE |
| --- | --- |
| | p-S6-CellBody-RadialDistribution-FracAtD-ex555nm-4of4-log2-predicted |
| | p-S6-CellBody-RadialDistribution-MeanFrac-ex555nm-1of4-log2-predicted |
| | p-S6-CellBody-RadialDistribution-MeanFrac-ex555nm-2of4-log2-predicted |
| | p-S6-CellBody-RadialDistribution-MeanFrac-ex555nm-3of4-log2-predicted |
| | p-S6-CellBody-RadialDistribution-MeanFrac-ex555nm-4of4-log2-predicted |
| | p-S6-CellBody-RadialDistribution-ZernikeMagnitude-ex555nm-1-1-log2-predicted |
| | p-S6-CellBody-RadialDistribution-ZernikeMagnitude-ex555nm-2-0-log2-predicted |
| | p-S6-CellBody-RadialDistribution-ZernikeMagnitude-ex555nm-2-2-log2-predicted |
| | p-S6-CellBody-RadialDistribution-ZernikeMagnitude-ex555nm-4-4-log2-predicted |
| | p-S6-CellBody-RadialDistribution-ZernikeMagnitude-ex555nm-6-6-log2-predicted |
| | p-S6-CellBody-RadialDistribution-ZernikeMagnitude-ex555nm-7-7-log2-predicted |
| | p-S6-CellBody-RadialDistribution-ZernikeMagnitude-ex555nm-8-8-log2-predicted |

*Continued on next page*

| Features used for E/M landscape UMAP | Features used for signaling landscape PHATE |
|---|---|
| | p-S6-CellBody-RadialDistribution-ZernikeMagnitude-ex555nm-9-9-log2-predicted |
| | p-S6-Cytoplasm-Intensity-IntegratedIntensityEdge-ex555nm-log2-predicted |
| | p-S6-Cytoplasm-Intensity-IntegratedIntensity-ex555nm-log2-predicted |
| | p-S6-Cytoplasm-Intensity-LowerQuartileIntensity-ex555nm-log2-predicted |
| | p-S6-Cytoplasm-Intensity-MeanIntensityEdge-ex555nm-log2-predicted |
| | p-S6-Cytoplasm-Intensity-MinIntensity-ex555nm-log2-predicted |
| | p-S6-Nuclei-Intensity-IntegratedIntensityEdge-ex555nm-log2-predicted |
| | p-S6-Nuclei-Intensity-IntegratedIntensity-ex555nm-log2-predicted |
| | p-S6-Ratio-NC-ex555nm-IntegInt-log2-predicted |
| | p-S6-Ratio-NC-ex555nm-MeanInt-log2-predicted |
| | p-AKT-T308-CellBody-Intensity-IntegratedIntensityEdge-ex555nm-log2-predicted |
| | p-AKT-T308-CellBody-Intensity-IntegratedIntensity-ex555nm-log2-predicted |
| | p-AKT-T308-CellBody-Intensity-LowerQuartileIntensity-ex555nm-log2-predicted |

*Continued on next page*

| Features used for E/M landscape UMAP | Features used for signaling landscape PHATE |
|---|---|
| | p-AKT-T308-CellBody-Intensity-MADIntensity-ex555nm-log2-predicted |
| | p-AKT-T308-CellBody-Intensity-MaxIntensity-ex555nm-log2-predicted |
| | p-AKT-T308-CellBody-Intensity-MeanIntensityEdge-ex555nm-log2-predicted |
| | p-AKT-T308-CellBody-Intensity-MeanIntensity-ex555nm-log2-predicted |
| | p-AKT-T308-CellBody-Intensity-MedianIntensity-ex555nm-log2-predicted |
| | p-AKT-T308-CellBody-Intensity-MinIntensityEdge-ex555nm-log2-predicted |
| | p-AKT-T308-CellBody-Intensity-MinIntensity-ex555nm-log2-predicted |
| | p-AKT-T308-CellBody-Intensity-StdIntensityEdge-ex555nm-log2-predicted |
| | p-AKT-T308-CellBody-Intensity-StdIntensity-ex555nm-log2-predicted |
| | p-AKT-T308-CellBody-Intensity-UpperQuartileIntensity-ex555nm-log2-predicted |
| | p-AKT-T308-CellBody-RadialDistribution-FracAtD-ex555nm-1of4-log2-predicted |
| | p-AKT-T308-CellBody-RadialDistribution-FracAtD-ex555nm-2of4-log2-predicted |
| | p-AKT-T308-CellBody-RadialDistribution-FracAtD-ex555nm-3of4-log2-predicted |
| | p-AKT-T308-CellBody-RadialDistribution-FracAtD-ex555nm-4of4-log2-predicted |
| | p-AKT-T308-CellBody-RadialDistribution-MeanFrac-ex555nm-1of4-log2-predicted |
| | p-AKT-T308-CellBody-RadialDistribution-MeanFrac-ex555nm-2of4-log2-predicted |

*Continued on next page*

| Features used for E/M landscape UMAP | Features used for signaling landscape PHATE |
|---|---|
| | p-AKT-T308-CellBody-RadialDistribution-MeanFrac-ex555nm-3of4-log2-predicted |
| | p-AKT-T308-CellBody-RadialDistribution-MeanFrac-ex555nm-4of4-log2-predicted |
| | p-AKT-T308-CellBody-RadialDistribution-RadialCV-ex555nm-4of4-log2-predicted |
| | p-AKT-T308-CellBody-RadialDistribution-ZernikeMagnitude-ex555nm-0-0-log2-predicted |
| | p-AKT-T308-CellBody-RadialDistribution-ZernikeMagnitude-ex555nm-1-1-log2-predicted |
| | p-AKT-T308-CellBody-RadialDistribution-ZernikeMagnitude-ex555nm-2-0-log2-predicted |
| | p-AKT-T308-CellBody-RadialDistribution-ZernikeMagnitude-ex555nm-2-2-log2-predicted |
| | p-AKT-T308-CellBody-RadialDistribution-ZernikeMagnitude-ex555nm-3-1-log2-predicted |
| | p-AKT-T308-CellBody-RadialDistribution-ZernikeMagnitude-ex555nm-3-3-log2-predicted |
| | p-AKT-T308-CellBody-RadialDistribution-ZernikeMagnitude-ex555nm-4-2-log2-predicted |
| | p-AKT-T308-CellBody-RadialDistribution-ZernikeMagnitude-ex555nm-4-4-log2-predicted |

*Continued on next page*

| Features used for E/M landscape UMAP | Features used for signaling landscape PHATE |
|---|---|
| | p-AKT-T308-CellBody-RadialDistribution-ZernikeMagnitude-ex555nm-5-3-log2-predicted |
| | p-AKT-T308-CellBody-RadialDistribution-ZernikeMagnitude-ex555nm-5-5-log2-predicted |
| | p-AKT-T308-CellBody-RadialDistribution-ZernikeMagnitude-ex555nm-6-4-log2-predicted |
| | p-AKT-T308-CellBody-RadialDistribution-ZernikeMagnitude-ex555nm-6-6-log2-predicted |
| | p-AKT-T308-CellBody-RadialDistribution-ZernikeMagnitude-ex555nm-7-5-log2-predicted |
| | p-AKT-T308-CellBody-RadialDistribution-ZernikeMagnitude-ex555nm-7-7-log2-predicted |
| | p-AKT-T308-CellBody-RadialDistribution-ZernikeMagnitude-ex555nm-8-6-log2-predicted |
| | p-AKT-T308-CellBody-RadialDistribution-ZernikeMagnitude-ex555nm-8-8-log2-predicted |
| | p-AKT-T308-CellBody-RadialDistribution-ZernikeMagnitude-ex555nm-9-7-log2-predicted |
| | p-AKT-T308-CellBody-RadialDistribution-ZernikeMagnitude-ex555nm-9-9-log2-predicted |

*Continued on next page*

| Features used for E/M landscape UMAP | Features used for signaling landscape PHATE |
|---|---|
| | p-AKT-T308-CellBody-Texture-AngularSecondMoment-ex555nm-3-00-256-log2-predicted |
| | p-AKT-T308-CellBody-Texture-AngularSecondMoment-ex555nm-3-01-256-log2-predicted |
| | p-AKT-T308-CellBody-Texture-AngularSecondMoment-ex555nm-3-02-256-log2-predicted |
| | p-AKT-T308-CellBody-Texture-AngularSecondMoment-ex555nm-3-03-256-log2-predicted |
| | p-AKT-T308-CellBody-Texture-Contrast-ex555nm-3-00-256-sqrt-predicted |
| | p-AKT-T308-CellBody-Texture-Contrast-ex555nm-3-01-256-sqrt-predicted |
| | p-AKT-T308-CellBody-Texture-Contrast-ex555nm-3-02-256-sqrt-predicted |
| | p-AKT-T308-CellBody-Texture-Contrast-ex555nm-3-03-256-sqrt-predicted |
| | p-AKT-T308-CellBody-Texture-DifferenceEntropy-ex555nm-3-00-256-raw-predicted |
| | p-AKT-T308-CellBody-Texture-DifferenceEntropy-ex555nm-3-01-256-raw-predicted |
| | p-AKT-T308-CellBody-Texture-DifferenceEntropy-ex555nm-3-02-256-raw-predicted |
| | p-AKT-T308-CellBody-Texture-DifferenceEntropy-ex555nm-3-03-256-raw-predicted |

*Continued on next page*

| Features used for E/M landscape UMAP | Features used for signaling landscape PHATE |
|---|---|
| | p-AKT-T308-CellBody-Texture-DifferenceVariance-ex555nm-3-00-256-log2-predicted |
| | p-AKT-T308-CellBody-Texture-DifferenceVariance-ex555nm-3-01-256-log2-predicted |
| | p-AKT-T308-CellBody-Texture-DifferenceVariance-ex555nm-3-02-256-log2-predicted |
| | p-AKT-T308-CellBody-Texture-DifferenceVariance-ex555nm-3-03-256-log2-predicted |
| | p-AKT-T308-CellBody-Texture-Entropy-ex555nm-3-00-256-raw-predicted |
| | p-AKT-T308-CellBody-Texture-Entropy-ex555nm-3-01-256-raw-predicted |
| | p-AKT-T308-CellBody-Texture-Entropy-ex555nm-3-02-256-raw-predicted |
| | p-AKT-T308-CellBody-Texture-Entropy-ex555nm-3-03-256-raw-predicted |
| | p-AKT-T308-CellBody-Texture-InfoMeas2-ex555nm-3-00-256-raw-predicted |
| | p-AKT-T308-CellBody-Texture-InfoMeas2-ex555nm-3-01-256-raw-predicted |
| | p-AKT-T308-CellBody-Texture-InfoMeas2-ex555nm-3-02-256-raw-predicted |
| | p-AKT-T308-CellBody-Texture-InfoMeas2-ex555nm-3-03-256-raw-predicted |
| | p-AKT-T308-CellBody-Texture-InverseDifferenceMoment-ex555nm-3-00-256-raw-predicted |

*Continued on next page*

| Features used for E/M landscape UMAP | Features used for signaling landscape PHATE |
|---|---|
| | p-AKT-T308-CellBody-Texture-InverseDifferenceMoment-ex555nm-3-01-256-raw-predicted |
| | p-AKT-T308-CellBody-Texture-InverseDifferenceMoment-ex555nm-3-02-256-raw-predicted |
| | p-AKT-T308-CellBody-Texture-InverseDifferenceMoment-ex555nm-3-03-256-raw-predicted |
| | p-AKT-T308-CellBody-Texture-SumAverage-ex555nm-3-00-256-log2-predicted |
| | p-AKT-T308-CellBody-Texture-SumAverage-ex555nm-3-01-256-log2-predicted |
| | p-AKT-T308-CellBody-Texture-SumAverage-ex555nm-3-02-256-log2-predicted |
| | p-AKT-T308-CellBody-Texture-SumAverage-ex555nm-3-03-256-log2-predicted |
| | p-AKT-T308-CellBody-Texture-SumEntropy-ex555nm-3-00-256-raw-predicted |
| | p-AKT-T308-CellBody-Texture-SumEntropy-ex555nm-3-01-256-raw-predicted |
| | p-AKT-T308-CellBody-Texture-SumEntropy-ex555nm-3-02-256-raw-predicted |
| | p-AKT-T308-CellBody-Texture-SumEntropy-ex555nm-3-03-256-raw-predicted |
| | p-AKT-T308-Cytoplasm-Intensity-IntegratedIntensityEdge-ex555nm-log2-predicted |
| | p-AKT-T308-Cytoplasm-Intensity-IntegratedIntensity-ex555nm-log2-predicted |

*Continued on next page*

| Features used for E/M landscape UMAP | Features used for signaling landscape PHATE |
|---|---|
| | p-AKT-T308-Cytoplasm-Intensity-LowerQuartileIntensity-ex555nm-log2-predicted |
| | p-AKT-T308-Cytoplasm-Intensity-MADIntensity-ex555nm-log2-predicted |
| | p-AKT-T308-Cytoplasm-Intensity-MaxIntensity-ex555nm-log2-predicted |
| | p-AKT-T308-Cytoplasm-Intensity-MeanIntensityEdge-ex555nm-log2-predicted |
| | p-AKT-T308-Cytoplasm-Intensity-MeanIntensity-ex555nm-log2-predicted |
| | p-AKT-T308-Cytoplasm-Intensity-MedianIntensity-ex555nm-log2-predicted |
| | p-AKT-T308-Cytoplasm-Intensity-MinIntensityEdge-ex555nm-log2-predicted |
| | p-AKT-T308-Cytoplasm-Intensity-MinIntensity-ex555nm-log2-predicted |
| | p-AKT-T308-Cytoplasm-Intensity-StdIntensityEdge-ex555nm-log2-predicted |
| | p-AKT-T308-Cytoplasm-Intensity-StdIntensity-ex555nm-log2-predicted |
| | p-AKT-T308-Cytoplasm-Intensity-UpperQuartileIntensity-ex555nm-log2-predicted |
| | p-AKT-T308-Nuclei-Intensity-IntegratedIntensityEdge-ex555nm-log2-predicted |
| | p-AKT-T308-Nuclei-Intensity-IntegratedIntensity-ex555nm-log2-predicted |
| | p-AKT-T308-Nuclei-Intensity-LowerQuartileIntensity-ex555nm-log2-predicted |

*Continued on next page*

| Features used for E/M landscape UMAP | Features used for signaling landscape PHATE |
|---|---|
| | p-AKT-T308-Nuclei-Intensity-MADIntensity-ex555nm-log2-predicted |
| | p-AKT-T308-Nuclei-Intensity-MeanIntensityEdge-ex555nm-log2-predicted |
| | p-AKT-T308-Nuclei-Intensity-MeanIntensity-ex555nm-log2-predicted |
| | p-AKT-T308-Nuclei-Intensity-MedianIntensity-ex555nm-log2-predicted |
| | p-AKT-T308-Nuclei-Intensity-MinIntensityEdge-ex555nm-log2-predicted |
| | p-AKT-T308-Nuclei-Intensity-MinIntensity-ex555nm-log2-predicted |
| | p-AKT-T308-Nuclei-Intensity-StdIntensity-ex555nm-log2-predicted |
| | p-AKT-T308-Nuclei-Intensity-UpperQuartileIntensity-ex555nm-log2-predicted |
| | p-AKT-T308-Nuclei-Texture-AngularSecondMoment-ex555nm-3-00-256-log2-predicted |
| | p-AKT-T308-Nuclei-Texture-AngularSecondMoment-ex555nm-3-01-256-log2-predicted |
| | p-AKT-T308-Nuclei-Texture-AngularSecondMoment-ex555nm-3-02-256-log2-predicted |
| | p-AKT-T308-Nuclei-Texture-AngularSecondMoment-ex555nm-3-03-256-log2-predicted |
| | p-AKT-T308-Nuclei-Texture-Contrast-ex555nm-3-00-256-sqrt-predicted |

*Continued on next page*

| Features used for E/M landscape UMAP | Features used for signaling landscape PHATE |
|---|---|
| | p-AKT-T308-Nuclei-Texture-Contrast-ex555nm-3-01-256-sqrt-predicted |
| | p-AKT-T308-Nuclei-Texture-Contrast-ex555nm-3-02-256-sqrt-predicted |
| | p-AKT-T308-Nuclei-Texture-Contrast-ex555nm-3-03-256-sqrt-predicted |
| | p-AKT-T308-Nuclei-Texture-DifferenceEntropy-ex555nm-3-00-256-raw-predicted |
| | p-AKT-T308-Nuclei-Texture-DifferenceEntropy-ex555nm-3-01-256-raw-predicted |
| | p-AKT-T308-Nuclei-Texture-DifferenceEntropy-ex555nm-3-02-256-raw-predicted |
| | p-AKT-T308-Nuclei-Texture-DifferenceEntropy-ex555nm-3-03-256-raw-predicted |
| | p-AKT-T308-Nuclei-Texture-DifferenceVariance-ex555nm-3-00-256-log2-predicted |
| | p-AKT-T308-Nuclei-Texture-DifferenceVariance-ex555nm-3-01-256-log2-predicted |
| | p-AKT-T308-Nuclei-Texture-DifferenceVariance-ex555nm-3-02-256-log2-predicted |
| | p-AKT-T308-Nuclei-Texture-DifferenceVariance-ex555nm-3-03-256-log2-predicted |
| | p-AKT-T308-Nuclei-Texture-Entropy-ex555nm-3-00-256-raw-predicted |

*Continued on next page*

| Features used for E/M landscape UMAP | Features used for signaling landscape PHATE |
| --- | --- |
| | p-AKT-T308-Nuclei-Texture-Entropy-ex555nm-3-01-256-raw-predicted |
| | p-AKT-T308-Nuclei-Texture-Entropy-ex555nm-3-02-256-raw-predicted |
| | p-AKT-T308-Nuclei-Texture-Entropy-ex555nm-3-03-256-raw-predicted |
| | p-AKT-T308-Nuclei-Texture-InverseDifferenceMoment-ex555nm-3-00-256-raw-predicted |
| | p-AKT-T308-Nuclei-Texture-InverseDifferenceMoment-ex555nm-3-01-256-raw-predicted |
| | p-AKT-T308-Nuclei-Texture-InverseDifferenceMoment-ex555nm-3-02-256-raw-predicted |
| | p-AKT-T308-Nuclei-Texture-InverseDifferenceMoment-ex555nm-3-03-256-raw-predicted |
| | p-AKT-T308-Nuclei-Texture-SumAverage-ex555nm-3-00-256-log2-predicted |
| | p-AKT-T308-Nuclei-Texture-SumAverage-ex555nm-3-01-256-log2-predicted |
| | p-AKT-T308-Nuclei-Texture-SumAverage-ex555nm-3-02-256-log2-predicted |
| | p-AKT-T308-Nuclei-Texture-SumAverage-ex555nm-3-03-256-log2-predicted |
| | p-AKT-T308-Nuclei-Texture-SumEntropy-ex555nm-3-00-256-raw-predicted |
| | p-AKT-T308-Nuclei-Texture-SumEntropy-ex555nm-3-01-256-raw-predicted |
| | p-AKT-T308-Nuclei-Texture-SumEntropy-ex555nm-3-02-256-raw-predicted |

*Continued on next page*

| Features used for E/M landscape UMAP | Features used for signaling landscape PHATE |
|---|---|
| | p-AKT-T308-Nuclei-Texture-SumEntropy-ex555nm-3-03-256-raw-predicted |
| | p-AKT-T308-Ratio-NC-ex555nm-IntegInt-log2-predicted |
| | p-AKT-T308-Ratio-NC-ex555nm-MeanInt-log2-predicted |
| | p-AKT-S473-CellBody-Intensity-IntegratedIntensityEdge-ex555nm-log2-predicted |
| | p-AKT-S473-CellBody-Intensity-IntegratedIntensity-ex555nm-log2-predicted |
| | p-AKT-S473-CellBody-Intensity-LowerQuartileIntensity-ex555nm-log2-predicted |
| | p-AKT-S473-CellBody-Intensity-MADIntensity-ex555nm-log2-predicted |
| | p-AKT-S473-CellBody-Intensity-MaxIntensityEdge-ex555nm-log2-predicted |
| | p-AKT-S473-CellBody-Intensity-MaxIntensity-ex555nm-log2-predicted |
| | p-AKT-S473-CellBody-Intensity-MeanIntensityEdge-ex555nm-log2-predicted |
| | p-AKT-S473-CellBody-Intensity-MeanIntensity-ex555nm-log2-predicted |
| | p-AKT-S473-CellBody-Intensity-MedianIntensity-ex555nm-log2-predicted |
| | p-AKT-S473-CellBody-Intensity-MinIntensityEdge-ex555nm-log2-predicted |
| | p-AKT-S473-CellBody-Intensity-MinIntensity-ex555nm-log2-predicted |
| | p-AKT-S473-CellBody-Intensity-StdIntensityEdge-ex555nm-log2-predicted |

*Continued on next page*

| Features used for E/M landscape UMAP | Features used for signaling landscape PHATE |
|---|---|
| | p-AKT-S473-CellBody-Intensity-StdIntensity-ex555nm-log2-predicted |
| | p-AKT-S473-CellBody-Intensity-UpperQuartileIntensity-ex555nm-log2-predicted |
| | p-AKT-S473-CellBody-RadialDistribution-FracAtD-ex555nm-1of4-log2-predicted |
| | p-AKT-S473-CellBody-RadialDistribution-FracAtD-ex555nm-2of4-log2-predicted |
| | p-AKT-S473-CellBody-RadialDistribution-FracAtD-ex555nm-3of4-log2-predicted |
| | p-AKT-S473-CellBody-RadialDistribution-FracAtD-ex555nm-4of4-log2-predicted |
| | p-AKT-S473-CellBody-RadialDistribution-MeanFrac-ex555nm-2of4-log2-predicted |
| | p-AKT-S473-CellBody-RadialDistribution-MeanFrac-ex555nm-4of4-log2-predicted |
| | p-AKT-S473-CellBody-RadialDistribution-RadialCV-ex555nm-4of4-log2-predicted |
| | p-AKT-S473-CellBody-RadialDistribution-ZernikeMagnitude-ex555nm-0-0-log2-predicted |
| | p-AKT-S473-CellBody-RadialDistribution-ZernikeMagnitude-ex555nm-1-1-log2-predicted |
| | p-AKT-S473-CellBody-RadialDistribution-ZernikeMagnitude-ex555nm-2-0-log2-predicted |
| | p-AKT-S473-CellBody-RadialDistribution-ZernikeMagnitude-ex555nm-2-2-log2-predicted |

*Continued on next page*

| Features used for E/M landscape UMAP | Features used for signaling landscape PHATE |
|---|---|
| | p-AKT-S473-CellBody-RadialDistribution-ZernikeMagnitude-ex555nm-3-1-log2-predicted |
| | p-AKT-S473-CellBody-RadialDistribution-ZernikeMagnitude-ex555nm-3-3-log2-predicted |
| | p-AKT-S473-CellBody-RadialDistribution-ZernikeMagnitude-ex555nm-4-2-log2-predicted |
| | p-AKT-S473-CellBody-RadialDistribution-ZernikeMagnitude-ex555nm-4-4-log2-predicted |
| | p-AKT-S473-CellBody-RadialDistribution-ZernikeMagnitude-ex555nm-5-1-log2-predicted |
| | p-AKT-S473-CellBody-RadialDistribution-ZernikeMagnitude-ex555nm-5-3-log2-predicted |
| | p-AKT-S473-CellBody-RadialDistribution-ZernikeMagnitude-ex555nm-5-5-log2-predicted |
| | p-AKT-S473-CellBody-RadialDistribution-ZernikeMagnitude-ex555nm-6-4-log2-predicted |
| | p-AKT-S473-CellBody-RadialDistribution-ZernikeMagnitude-ex555nm-6-6-log2-predicted |
| | p-AKT-S473-CellBody-RadialDistribution-ZernikeMagnitude-ex555nm-7-3-log2-predicted |

*Continued on next page*

| Features used for E/M landscape UMAP | Features used for signaling landscape PHATE |
|---|---|
| | p-AKT-S473-CellBody-RadialDistribution-ZernikeMagnitude-ex555nm-7-5-log2-predicted |
| | p-AKT-S473-CellBody-RadialDistribution-ZernikeMagnitude-ex555nm-7-7-log2-predicted |
| | p-AKT-S473-CellBody-RadialDistribution-ZernikeMagnitude-ex555nm-8-4-log2-predicted |
| | p-AKT-S473-CellBody-RadialDistribution-ZernikeMagnitude-ex555nm-8-6-log2-predicted |
| | p-AKT-S473-CellBody-RadialDistribution-ZernikeMagnitude-ex555nm-8-8-log2-predicted |
| | p-AKT-S473-CellBody-RadialDistribution-ZernikeMagnitude-ex555nm-9-5-log2-predicted |
| | p-AKT-S473-CellBody-RadialDistribution-ZernikeMagnitude-ex555nm-9-7-log2-predicted |
| | p-AKT-S473-CellBody-RadialDistribution-ZernikeMagnitude-ex555nm-9-9-log2-predicted |
| | p-AKT-S473-CellBody-Texture-AngularSecondMoment-ex555nm-3-00-256-log2-predicted |
| | p-AKT-S473-CellBody-Texture-AngularSecondMoment-ex555nm-3-01-256-log2-predicted |

*Continued on next page*

| Features used for E/M landscape UMAP | Features used for signaling landscape PHATE |
|---|---|
| | p-AKT-S473-CellBody-Texture-AngularSecondMoment-ex555nm-3-02-256-log2-predicted |
| | p-AKT-S473-CellBody-Texture-AngularSecondMoment-ex555nm-3-03-256-log2-predicted |
| | p-AKT-S473-CellBody-Texture-Contrast-ex555nm-3-00-256-sqrt-predicted |
| | p-AKT-S473-CellBody-Texture-Contrast-ex555nm-3-01-256-sqrt-predicted |
| | p-AKT-S473-CellBody-Texture-Contrast-ex555nm-3-02-256-sqrt-predicted |
| | p-AKT-S473-CellBody-Texture-Contrast-ex555nm-3-03-256-sqrt-predicted |
| | p-AKT-S473-CellBody-Texture-DifferenceEntropy-ex555nm-3-00-256-raw-predicted |
| | p-AKT-S473-CellBody-Texture-DifferenceEntropy-ex555nm-3-01-256-raw-predicted |
| | p-AKT-S473-CellBody-Texture-DifferenceEntropy-ex555nm-3-02-256-raw-predicted |
| | p-AKT-S473-CellBody-Texture-DifferenceEntropy-ex555nm-3-03-256-raw-predicted |
| | p-AKT-S473-CellBody-Texture-DifferenceVariance-ex555nm-3-00-256-log2-predicted |
| | p-AKT-S473-CellBody-Texture-DifferenceVariance-ex555nm-3-01-256-log2-predicted |

*Continued on next page*

| Features used for E/M landscape UMAP | Features used for signaling landscape PHATE |
|---|---|
| | p-AKT-S473-CellBody-Texture-DifferenceVariance-ex555nm-3-02-256-log2-predicted |
| | p-AKT-S473-CellBody-Texture-DifferenceVariance-ex555nm-3-03-256-log2-predicted |
| | p-AKT-S473-CellBody-Texture-Entropy-ex555nm-3-00-256-raw-predicted |
| | p-AKT-S473-CellBody-Texture-Entropy-ex555nm-3-01-256-raw-predicted |
| | p-AKT-S473-CellBody-Texture-Entropy-ex555nm-3-02-256-raw-predicted |
| | p-AKT-S473-CellBody-Texture-Entropy-ex555nm-3-03-256-raw-predicted |
| | p-AKT-S473-CellBody-Texture-InverseDifferenceMoment-ex555nm-3-00-256-raw-predicted |
| | p-AKT-S473-CellBody-Texture-InverseDifferenceMoment-ex555nm-3-01-256-raw-predicted |
| | p-AKT-S473-CellBody-Texture-InverseDifferenceMoment-ex555nm-3-02-256-raw-predicted |
| | p-AKT-S473-CellBody-Texture-InverseDifferenceMoment-ex555nm-3-03-256-raw-predicted |
| | p-AKT-S473-CellBody-Texture-SumAverage-ex555nm-3-00-256-log2-predicted |
| | p-AKT-S473-CellBody-Texture-SumAverage-ex555nm-3-01-256-log2-predicted |
| | p-AKT-S473-CellBody-Texture-SumAverage-ex555nm-3-02-256-log2-predicted |

*Continued on next page*

| Features used for E/M landscape UMAP | Features used for signaling landscape PHATE |
| --- | --- |
| | p-AKT-S473-CellBody-Texture-SumAverage-ex555nm-3-03-256-log2-predicted |
| | p-AKT-S473-CellBody-Texture-SumEntropy-ex555nm-3-00-256-raw-predicted |
| | p-AKT-S473-CellBody-Texture-SumEntropy-ex555nm-3-01-256-raw-predicted |
| | p-AKT-S473-CellBody-Texture-SumEntropy-ex555nm-3-02-256-raw-predicted |
| | p-AKT-S473-CellBody-Texture-SumEntropy-ex555nm-3-03-256-raw-predicted |
| | p-AKT-S473-Cytoplasm-Intensity-IntegratedIntensityEdge-ex555nm-log2-predicted |
| | p-AKT-S473-Cytoplasm-Intensity-IntegratedIntensity-ex555nm-log2-predicted |
| | p-AKT-S473-Cytoplasm-Intensity-LowerQuartileIntensity-ex555nm-log2-predicted |
| | p-AKT-S473-Cytoplasm-Intensity-MADIntensity-ex555nm-log2-predicted |
| | p-AKT-S473-Cytoplasm-Intensity-MaxIntensityEdge-ex555nm-log2-predicted |
| | p-AKT-S473-Cytoplasm-Intensity-MaxIntensity-ex555nm-log2-predicted |
| | p-AKT-S473-Cytoplasm-Intensity-MeanIntensityEdge-ex555nm-log2-predicted |
| | p-AKT-S473-Cytoplasm-Intensity-MeanIntensity-ex555nm-log2-predicted |
| | p-AKT-S473-Cytoplasm-Intensity-MedianIntensity-ex555nm-log2-predicted |
| | p-AKT-S473-Cytoplasm-Intensity-MinIntensityEdge-ex555nm-log2-predicted |

*Continued on next page*

| Features used for E/M landscape UMAP | Features used for signaling landscape PHATE |
|---|---|
| | p-AKT-S473-Cytoplasm-Intensity-MinIntensity-ex555nm-log2-predicted |
| | p-AKT-S473-Cytoplasm-Intensity-StdIntensityEdge-ex555nm-log2-predicted |
| | p-AKT-S473-Cytoplasm-Intensity-StdIntensity-ex555nm-log2-predicted |
| | p-AKT-S473-Cytoplasm-Intensity-UpperQuartileIntensity-ex555nm-log2-predicted |
| | p-AKT-S473-Nuclei-Intensity-IntegratedIntensityEdge-ex555nm-log2-predicted |
| | p-AKT-S473-Nuclei-Intensity-IntegratedIntensity-ex555nm-log2-predicted |
| | p-AKT-S473-Nuclei-Intensity-LowerQuartileIntensity-ex555nm-log2-predicted |
| | p-AKT-S473-Nuclei-Intensity-MADIntensity-ex555nm-log2-predicted |
| | p-AKT-S473-Nuclei-Intensity-MaxIntensityEdge-ex555nm-log2-predicted |
| | p-AKT-S473-Nuclei-Intensity-MaxIntensity-ex555nm-log2-predicted |
| | p-AKT-S473-Nuclei-Intensity-MeanIntensityEdge-ex555nm-log2-predicted |
| | p-AKT-S473-Nuclei-Intensity-MeanIntensity-ex555nm-log2-predicted |
| | p-AKT-S473-Nuclei-Intensity-MedianIntensity-ex555nm-log2-predicted |
| | p-AKT-S473-Nuclei-Intensity-MinIntensityEdge-ex555nm-log2-predicted |
| | p-AKT-S473-Nuclei-Intensity-MinIntensity-ex555nm-log2-predicted |

*Continued on next page*

| Features used for E/M landscape UMAP | Features used for signaling landscape PHATE |
|---|---|
| | p-AKT-S473-Nuclei-Intensity-StdIntensityEdge-ex555nm-log2-predicted |
| | p-AKT-S473-Nuclei-Intensity-StdIntensity-ex555nm-log2-predicted |
| | p-AKT-S473-Nuclei-Intensity-UpperQuartileIntensity-ex555nm-log2-predicted |
| | p-AKT-S473-Nuclei-Texture-AngularSecondMoment-ex555nm-3-00-256-log2-predicted |
| | p-AKT-S473-Nuclei-Texture-AngularSecondMoment-ex555nm-3-01-256-log2-predicted |
| | p-AKT-S473-Nuclei-Texture-AngularSecondMoment-ex555nm-3-02-256-log2-predicted |
| | p-AKT-S473-Nuclei-Texture-AngularSecondMoment-ex555nm-3-03-256-log2-predicted |
| | p-AKT-S473-Nuclei-Texture-Contrast-ex555nm-3-00-256-sqrt-predicted |
| | p-AKT-S473-Nuclei-Texture-Contrast-ex555nm-3-01-256-sqrt-predicted |
| | p-AKT-S473-Nuclei-Texture-Contrast-ex555nm-3-02-256-sqrt-predicted |
| | p-AKT-S473-Nuclei-Texture-Contrast-ex555nm-3-03-256-sqrt-predicted |
| | p-AKT-S473-Nuclei-Texture-DifferenceEntropy-ex555nm-3-00-256-raw-predicted |

*Continued on next page*

| Features used for E/M landscape UMAP | Features used for signaling landscape PHATE |
|---|---|
| | p-AKT-S473-Nuclei-Texture-DifferenceEntropy-ex555nm-3-01-256-raw-predicted |
| | p-AKT-S473-Nuclei-Texture-DifferenceEntropy-ex555nm-3-02-256-raw-predicted |
| | p-AKT-S473-Nuclei-Texture-DifferenceEntropy-ex555nm-3-03-256-raw-predicted |
| | p-AKT-S473-Nuclei-Texture-DifferenceVariance-ex555nm-3-00-256-log2-predicted |
| | p-AKT-S473-Nuclei-Texture-DifferenceVariance-ex555nm-3-01-256-log2-predicted |
| | p-AKT-S473-Nuclei-Texture-DifferenceVariance-ex555nm-3-02-256-log2-predicted |
| | p-AKT-S473-Nuclei-Texture-DifferenceVariance-ex555nm-3-03-256-log2-predicted |
| | p-AKT-S473-Nuclei-Texture-Entropy-ex555nm-3-00-256-raw-predicted |
| | p-AKT-S473-Nuclei-Texture-Entropy-ex555nm-3-01-256-raw-predicted |
| | p-AKT-S473-Nuclei-Texture-Entropy-ex555nm-3-02-256-raw-predicted |
| | p-AKT-S473-Nuclei-Texture-Entropy-ex555nm-3-03-256-raw-predicted |
| | p-AKT-S473-Nuclei-Texture-InverseDifferenceMoment-ex555nm-3-00-256-raw-predicted |

*Continued on next page*

| Features used for E/M landscape UMAP | Features used for signaling landscape PHATE |
|---|---|
| | p-AKT-S473-Nuclei-Texture-InverseDifferenceMoment-ex555nm-3-01-256-raw-predicted |
| | p-AKT-S473-Nuclei-Texture-InverseDifferenceMoment-ex555nm-3-02-256-raw-predicted |
| | p-AKT-S473-Nuclei-Texture-InverseDifferenceMoment-ex555nm-3-03-256-raw-predicted |
| | p-AKT-S473-Nuclei-Texture-SumEntropy-ex555nm-3-00-256-raw-predicted |
| | p-AKT-S473-Nuclei-Texture-SumEntropy-ex555nm-3-01-256-raw-predicted |
| | p-AKT-S473-Nuclei-Texture-SumEntropy-ex555nm-3-02-256-raw-predicted |
| | p-AKT-S473-Nuclei-Texture-SumEntropy-ex555nm-3-03-256-raw-predicted |
| | p-AKT-S473-Nuclei-Texture-SumVariance-ex555nm-3-00-256-sqrt-predicted |
| | p-AKT-S473-Nuclei-Texture-SumVariance-ex555nm-3-03-256-sqrt-predicted |
| | p-AKT-S473-Nuclei-Texture-Variance-ex555nm-3-00-256-sqrt-predicted |
| | p-AKT-S473-Nuclei-Texture-Variance-ex555nm-3-01-256-sqrt-predicted |
| | p-AKT-S473-Nuclei-Texture-Variance-ex555nm-3-02-256-sqrt-predicted |
| | p-AKT-S473-Nuclei-Texture-Variance-ex555nm-3-03-256-sqrt-predicted |
| | p-AKT-S473-Ratio-NC-ex555nm-IntegInt-log2-predicted |

| | |
|---|---|
| | p-AKT-S473-Ratio-NC-ex555nm-MeanInt-log2-predicted |

*Continued on next page*
